# Induced degradation of SNAP-fusion proteins

**DOI:** 10.1101/2024.07.18.603056

**Authors:** Savina Abraham Pol, Sara Liljenberg, Jack Barr, Gina Simon, Luis Wong-Dilworth, Danielle L. Paterson, Vladimir P. Berishvili, Francesca Bottanelli, Farnusch Kaschani, Markus Kaiser, Mariell Pettersson, Doris Hellerschmied

## Abstract

Self-labeling protein tags are an efficient means to visualize, manipulate, and isolate engineered fusion proteins with suitable chemical probes. The SNAP-tag, which covalently conjugates to benzyl-guanine and -chloropyrimidine derivatives is used extensively in fluorescence microscopy, given the availability of suitable SNAP-ligand-based probes. Here, we extend the applicability of the SNAP-tag to targeted protein degradation. We developed a set of SNAP PROteolysis TArgeting Chimeras (SNAP-PROTACs), which recruit the VHL or CRBN-ubiquitin E3 ligases to induce the degradation of SNAP-fusion proteins. Endogenous tagging enabled the visualization and the selective depletion of a SNAP-clathrin light chain fusion protein using SNAP-PROTACs. The addition of PROTACs to the SNAP-tag reagent toolbox facilitates the comprehensive analysis of protein function with a single gene tagging event.

## INTRODUCTION

Studying the function of proteins inside cells and within organisms is key to our understanding of cellular processes and complex biological systems. To selectively visualize, isolate, or perturb proteins of interest (POIs), cell biologists often resort to the generation of fusion proteins (1). For example, the genetic engineering of POIs with fluorescent protein domains is applied to study POI expression and localization, while the fusion of the POI with distinct modifying enzymes enables proximity biotinylation, thereby mapping the surroundings of the POI (1). To create a POI with multiple capabilities, the use of self-labeling protein tags (SLPs) is an attractive strategy (2, 3). SLPs are small protein domains, which covalently conjugate to their corresponding ligands (2, 3). These ligands can be functionalized with various chemical probes e.g. fluorescent moieties, relaying the technology development onto chemical synthesis. Following this chemical genetic strategy, a single genetic engineering event, linking the SLP to the POI, can provide access to a variety of technologies, provided the appropriate chemical probes are available (3). The advancement of genetic engineering strategies, such as CRISPR-Cas9-based gene editing, has even extended our ability to characterize endogenously tagged POIs at their natural expression level and regulation (4–6).

The SNAP-and CLIP-tags are versatile SLPs, originally derived from human *O*^6^-alkylguanine DNA acetyltransferase (7–10). While the SNAP-tag reacts with benzyl-guanine (SNAP1 ligand) or benzyl-chloropyrimidine (SNAP2 ligand) derivatives, the orthogonal CLIP-tag reacts with benzyl-cytosine (CLIP ligand) derivatives (7, 9, 11). Engineered SNAP-fusion proteins are extensively used to visualize POIs in high resolution (live) cell microscopy studies (12, 13). For this application, a multitude of (wash-free) fluorescent dyes spanning the near-UV, visible, and near-IR spectrum are available (14–16). SNAP-conjugating compounds serving as reporters on cellular metabolites and the protein homeostasis state of cells are also available (15, 17). Resins for pull-down studies enable the efficient isolation of SNAP-fusion proteins. Proximity labeling with a SNAP-conjugating photo-proximity probe enables mapping the cellular environment of SNAP-fusion proteins (18). Heterobifunctional compounds containing a SNAP ligand can be used to induce protein dimerization with other orthogonally tagged POIs (19).

A powerful chemical biology approach that relies on heterobifunctional compounds is targeted protein degradation (TPD) using PROteolysis TArgeting Chimeras (PROTACs) (20, 21) PROTACs are heterobifunctional small molecules that recruit the cellular ubiquitination machinery to a POI to induce its ubiquitination and subsequent degradation (20). The most frequently recruited ubiquitination enzymes are the Cullin-RING ubiquitin ligases, comprising either the van-Hippel-Lindau (VHL) or Cereblon (CRBN) substrate adaptors (22). The predominant recruitment of the VHL and CRBN substrate adaptors by the majority of the currently published PROTACs is driven by the availability of highly specific small molecule VHL and CRBN ligands (23). Equally, recruitment of the POI depends on the availability of a suitable ligand. Direct engagement of the POI by an active PROTAC provides the basis for TPD as a promising therapeutic modality (24). When ligands directly interacting with the POI are lacking, protein domains with available high affinity ligands can be leveraged as efficient recruiting elements. Combined with endogenous tagging of the POI, TPD approaches thereby provide an important strategy for target validation in drug discovery programs and loss-of-function studies in basic research (25, 26). For example, genetically engineered fusion proteins with the FKBP^F36V^ domain as part of the dTag system can be efficiently degraded by the use of VHL-or CRBN-recruiting PROTACs (27, 28). Similarly, the DHFR domain and the Bromodomain of BRD4 were turned into efficient tag-PROTAC systems (29, 30). In addition to these non-covalent tag-targeting PROTACs, covalent HaloPROTACs induce the acute and efficient degradation of HaloTag fusion proteins (31, 32). The self-labeling HaloTag covalently conjugates to synthetic chloroalkane ligands (33, 34). Functionalization of this ligand with VHL-recruiting ligands afforded highly specific covalent HaloPROTACs (31, 32).

PROTACs targeting POI-fusion proteins emerged as an attractive alternative to conventional RNAi-based knockdown or genetic knock-out approaches, as they address limitations inherent to these traditional methods (26, 35). Compared to genetic knock-outs, which represent an irreversible perturbation, TPD offers temporal control of POI depletion. Accordingly, TPD can circumvent cellular adaptation to prolonged POI depletion, a common occurrence in permanent genetic knock-outs, masking important loss-of-function phenotypes and confounding downstream analysis. Compared to RNAi-based approaches, which prevent the synthesis of new protein, TPD directly targets the pre-existing POI pool and frequently offers enhanced selectivity.

Here we describe the development of SNAP-PROTACs, thereby extending the applicability of the self-labeling SNAP-tag from protein isolation, visualization, and proximity biotinylation to TPD.

## DESIGN

The SNAP-tag is a small 20 kDa protein domain that covalently conjugates to benzyl-guanine and –chloropyrimidine derivatives under release of a leaving group (**Figure 1A** and **2A**) (9, 11). The phenyl ring (**Figure 2A**), which is covalently attached to the active site cysteine of the SNAP-tag, offers an ideal attachment point for the generation of SNAP-PROTACs. To develop versatile SNAP-PROTACs that can be applied to a wide range of cell types, we chose to recruit the VHL and CRBN E3 ligases using high affinity small molecule ligands. We explored different linker length and exit vectors from the E3 ligase ligands, as these define degradation potency and selectivity (36–38). For the VHL-recruiting PROTACs, we kept the amine exit vector constant (39) and varied the linker length (**Figure 1B** and **Scheme S1**). For the CRBN-recruiting PROTACs, we included two exit vectors (**Figure 4A**) and varied the linker length as well as composition (**Scheme S3**). In addition to flexible aliphatic linkers, we included piperidine-and piperazine-based linkers, commonly used to make PROTACs more rigid (**Scheme S3**). In an attempt to optimize the labeling efficiency of the SNAP-tag, we explored the SNAP-ligand with a focus on changing the electronics of its benzylic position (**Figure 2A**). To explore synthetic diversity, we also introduced handles that allow new chemistry for conjugation to the SNAP-ligand (**Figure 2B**). The resulting SNAP-PROTACs can easily be implemented and applied in cell lines expressing endogenously tagged SNAP-fusion proteins, readily available in many laboratories from previous use in imaging experiments. Here, we apply an efficient strategy for endogenous tagging in the haploid HAP1 cell line (40) and demonstrate the SNAP-PROTAC-induced degradation of a SNAP-clathrin light chain fusion protein.

**Figure 1.**
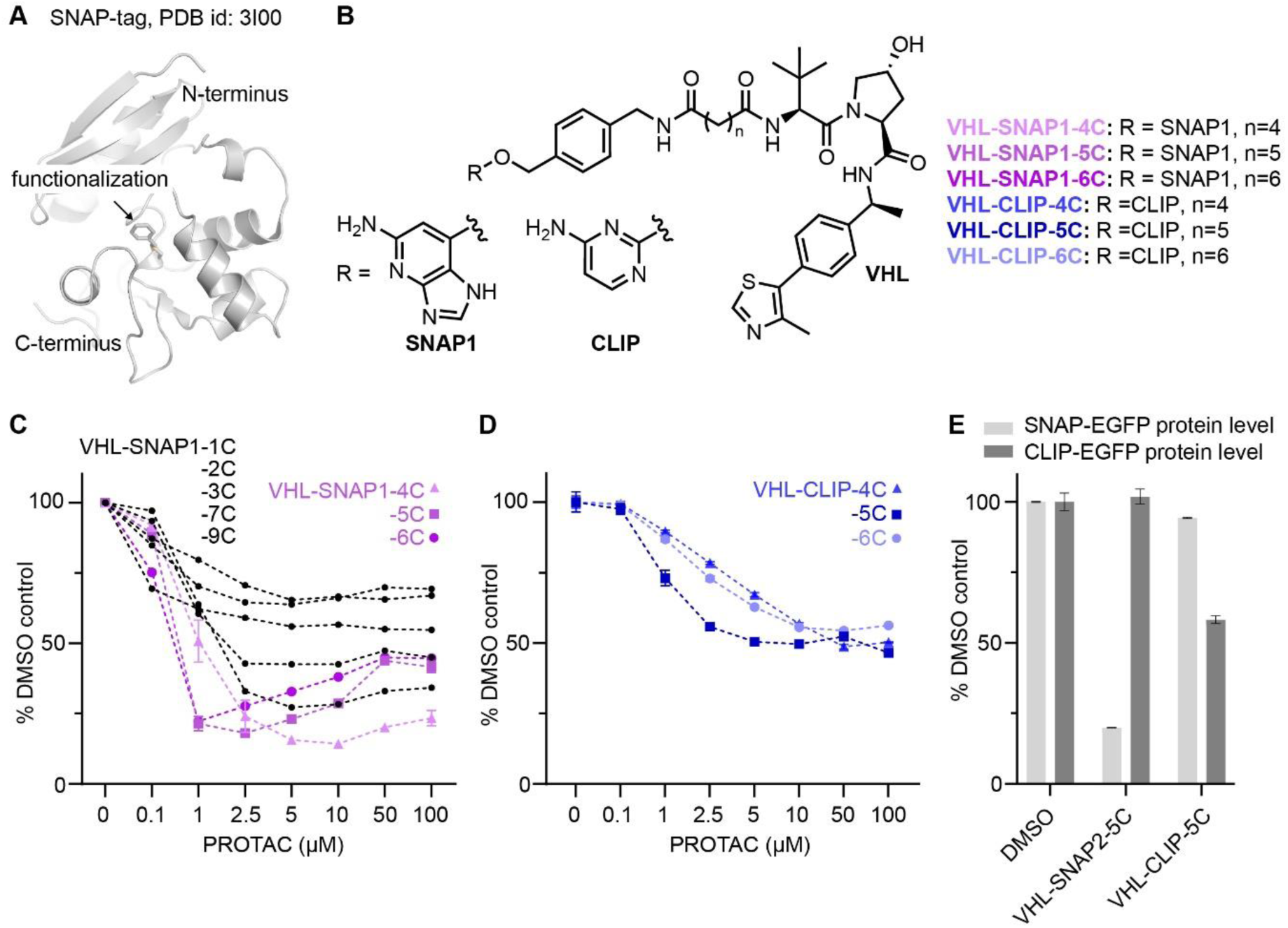
Development of VHL-recruiting SNAP-and CLIP-tag targeting PROTACs. (**A**) Crystal structure of benzylated SNAP-tag. (**B**) Chemical structures of selected PROTACs comprising the SNAP1-or CLIP ligands, linker, and VHL-recruiting ligand (related to scheme S1) (**C**) Flow cytometry analysis of SNAP-EGFP levels in HEK293 SNAP-EGFP cells treated with different concentrations of VHL-SNAP1-PROTACS for 24 h. (**D**) Flow cytometry analysis of CLIP-EGFP levels in HEK293 CLIP-EGFP cells treated with different concentrations of VHL-CLIP-PROTACS for 24 h. (**E**) Analysis of cross-reactivity of SNAP-and CLIP-targeting PROTACs. HEK293 cells expressing either SNAP-EGFP or CLIP-EGFP were treated for 24 h with 2.5 µM VHL-CLIP-5C or 1 µM VHL-SNAP2-5C. Decrease in EGFP-levels was measured by flow cytometry. (n = 3, data represent mean +/- s.d.)

**Figure 2.**
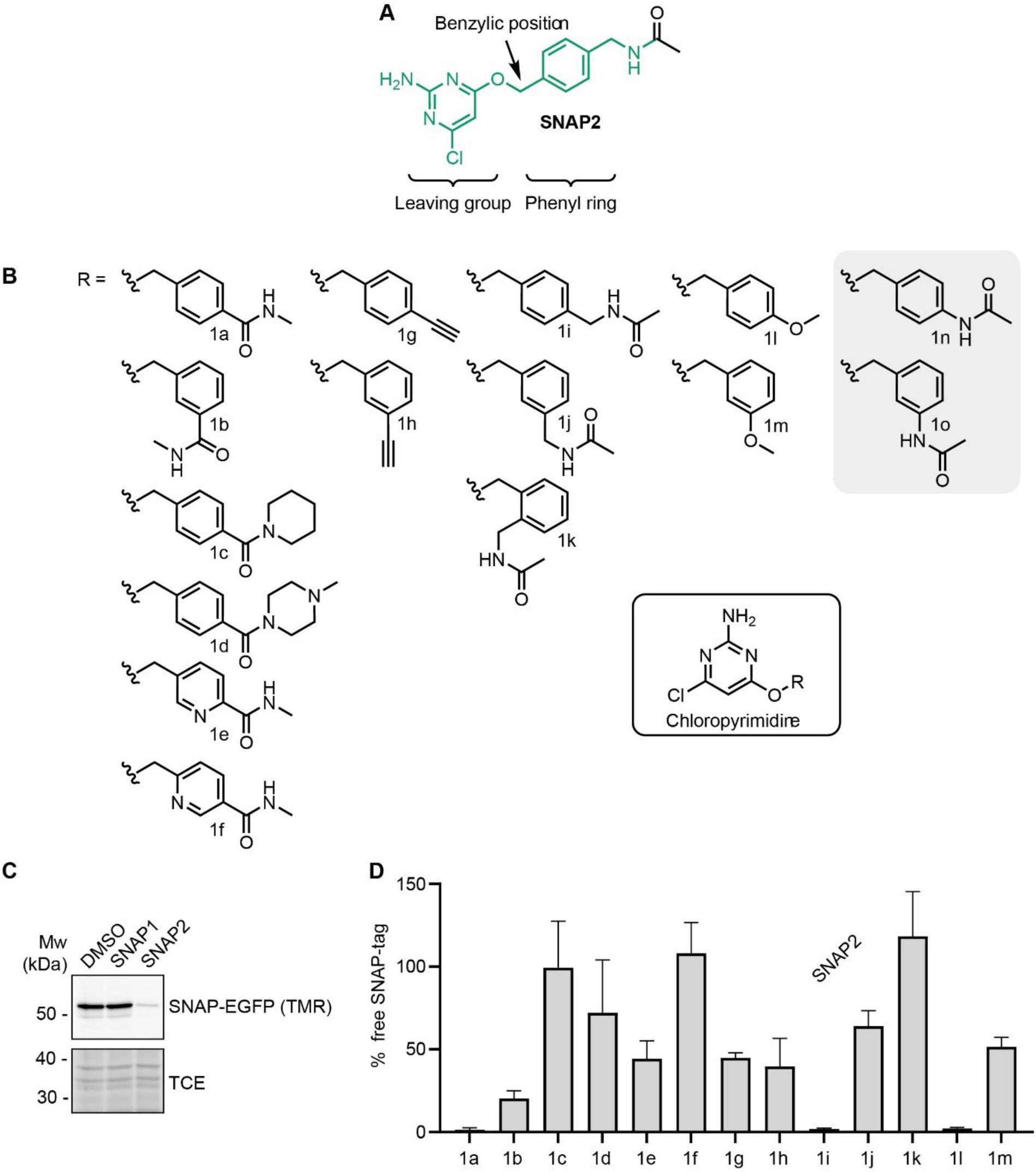
Optimization of the SNAP-tag recruiting ligand. (**A**) Structure of the SNAP2 ligand. (**B**) Chemical structures of the 15 chloropyrimidine-based SNAP ligands considered in this study. Ligands **1n** and **1o** were never synthesized. (**C**) SDS-PAGE of SNAP-EGFP-expressing HEK293 cells treated with SNAP1-or SNAP2 ligand for 15 min. Non-engaged SNAP-EGFP was labeled with SNAP-TMR dye to indirectly assess binding of the ligands. TCE signal was used as a loading control. (**D**) Quantification of the TMR-signal in HEK293 SNAP-EGFP cells treated for 15 min with the SNAP ligands shown in B (n = 4, data represent mean +/- s.d.).

## RESULTS

### Development of VHL-SNAP-PROTACs

To identify PROTACs that afford rapid and efficient degradation of SNAP-fusion proteins, we initially synthesized a set of compounds recruiting the VHL E3 ligase. We used the VHL-ligand depicted in **Figure 1B** and explored alkyl-linkers of varying length, extending from the amine exit vector, in combination with the SNAP1 ligand (**Scheme S1**). To screen the VHL-SNAP1-PROTAC series in a high-throughput manner, we generated a SNAP-EGFP-expressing HEK293 cell line, which enables quantification of protein levels by flow cytometry (**Figure 1B** and **S1A**). In a dose response PROTAC treatment for 24 h we observed a reduction of SNAP-EGFP levels for all tested VHL-SNAP1-PROTACs (**Figure 1C**). PROTACs with a linker length of n = 4-6 carbons proved most efficient in inducing SNAP-EGFP degradation.

On the basis of the VHL-SNAP-PROTACs, we also generated CLIP-tag targeting PROTACs with a linker length of n = 4-6 carbons (**Figure 1B**) and assessed their degradation efficiency by flow cytometry in a CLIP-EGFP-expressing HEK293 cell line (**Figure 1D** and **S1B**). The maximum degradation of CLIP-EGFP was ∼50% at 5-100 µM PROTAC concentration. The overall low potency of the CLIP-PROTACs may be due to their reduced solubility in growth medium, as strong precipitation was observed at concentrations of 50-100 µM. Nevertheless, to explore degradation selectivity, we tested the cross-reactivity of SNAP-and CLIP-targeting PROTACs (**Figure 1E**), given that minor cross-reactivity of TMR functionalized SNAP and CLIP-ligands was previously reported (41). Treating SNAP-EGFP expressing cells with 2.5 µM **VHL-CLIP-5C** resulted in a minor decrease (∼5%) of protein levels, while **VHL-SNAP2-5C** did not induce degradation of CLIP-EGFP at 1 µM concentration for 24 h.

Overall, our initial screening efforts revealed a linker length (n = 4-6 carbons) that enables productive degradation of the SNAP-EGFP fusion protein by the VHL E3 ligase, with minimal cross-reactivity with the CLIP-tag.

### Improving SNAP-ligand chemistry

We next wanted to improve the SNAP-PROTACs by optimizing the SNAP ligand. Previous work focused on protein engineering of the SNAP-tag and evaluating the influence of the guanine or pyrimidine leaving group of the SNAP ligand to increase reactivity (**Figure 2A**) (11, 41, 42). We therefore turned to the phenyl ring of the SNAP ligand (**Figure 2A**) with the goal of improving SNAP labeling efficiency. In addition, we wanted to expand the chemistry that could be utilized for SNAP-PROTAC synthesis and that potentially could further increase cell permeability. With these criteria in mind, we selected 15 SNAP ligands (**Figure 2B**) with varied substituents on the phenyl ring to afford differing electronic impact on the benzylic position, in combination with the previously described more cell-permeable chloropyrimidine-based SNAP2 ligand (**1i**) (11). In our design, we included reversed amides (**1a**-**f**), relative to the reference compound **1i**, allowing the exploration of more electron-deficient benzylic positions, as well as new chemistry. Amides from primary (**1a**-**b, e-f**) and secondary amines (**1c**-**d**) were also included, as many commercial building blocks and intermediate collections have terminal amines (43). Alkyne (**1g**-**h**) and ether (**1l**-**m**) substituents have the potential to improve cell permeability, by removal of the amide, and provide new chemistry. Differences in the substitution pattern on the phenyl ring (*ortho*, *meta*, or *para*, e.g. **1i**-**k**) affect the reactivity of the benzylic position. In the context of a PROTAC, different exit vectors can potentially change Removal of the methylene bridge of **1i** would provide **1n**, which would yield a more electron rich benzylic position, as well as reduced flexibility of the exit vector. We were able to synthesize 13 of the 15 selected ligands (**Figure 2B**). **1n** and **1o** could not be obtained, since we observed immediate degradation of any formed product during synthesis. We speculate that this instability may arise from promotion of a S_N_1 type pathway in electron rich substrates that generates a highly reactive benzylic carbocation *in situ*.

Labeling of the SNAP-tag depends on the formation of a SNAP-tag-ligand complex. Accordingly, Wilhelm et al. have previously seen a correlation between ligand binding affinities and SNAP-tag labeling kinetics (41). To categorize the 13 obtained chloropyrimidines (**1a-m**), we therefore set out to predict binding affinities. The synthesized compounds belong to a congeneric series (**Figure 2B**) and are all close analogues, which makes relative Free Energy Perturbation (FEP) calculations a suitable tool to assess binding affinities. We conducted FEP calculations starting from a crystal structure of the SNAP-tag in complex with the benzylguanine (PDB code: 3KZZ), assuming that benzylguanines share a similar binding mode with benzyl chloropyrimidines. Interestingly, four compounds (**1a**, **1c**, **1d**, and **1j**) are predicted to have affinities comparable to or better than the reference compound **1i** (**Table S1**). Compound **1k** was predicted not to form a productive protein-substrate complex due to steric clashes with the SNAP-tag (**Table S1**).

We tested the ability of the obtained compounds to engage the SNAP-EGFP protein in HEK293 cells, given our final goal to use them as target-recruiting elements in PROTACs. We indirectly monitored their engagement of the SNAP-tag, by labeling unengaged SNAP-EGFP protein in cell lysates with a SNAP-TMR dye. Initially we compared SNAP-TMR-labeling of the SNAP-EGFP protein upon treating cells with the SNAP1-and SNAP2-ligands. While faster reaction kinetics are described for SNAP1-versus SNAP2-ligands in experiments using purified SNAP-protein (41), in cells, the more cell-permeable SNAP2 ligand outperformed SNAP1 (**Figure 2C**). Upon 15 min incubation at 1 µM compound concentration, the SNAP2 ligand yielded a nearly complete block of SNAP-EGFP labeling with the SNAP-TMR dye. In fact, from the 13 compounds shown in **Figure 2B**, the SNAP2 ligand, reference compound **1i**, most potently blocked labeling with the SNAP-TMR dye (**Figure 2D**). Notably, two compounds, **1a** and **1l**, blocked SNAP-TMR labeling to a similar extent (**Figure 2D**). Compound **1l**, while engaging the SNAP-EGFP protein in cells, is unstable. Characterization of chemical stability (**Table S2**) showed a short half-life for **1l** at pH 7.4 (t_1/2_ ∼10 h) compared to **1i** and **1a** (t_1/2_ >149 days). To our surprise **1c** and **1d**, were inactive in cells. We further tried to rationalize this discrepancy between in-cell data and FEP calculations. However, **1c** showed comparable chemical stability to the reference compound **1i** (**Table S2**) and molecular docking showed that the piperidine substituent of **1c** is solvent exposed, excluding any steric clashes (**Figure S2**).

The *para*-substituted **1g** and *meta*-substituted compounds **1b**, **1h**, **1j**, and **1m**, showed reduced but robust engagement of the SNAP-tag (40-80%) in our assay. Of these, **1b**, carrying a reversed amide, showed the highest potency for blocking SNAP-TMR labeling. In line with the computational prediction, *ortho*-substituted **1k**, did not impair SNAP-TMR labeling, suggesting that this compound cannot engage with the SNAP-tag.

In summary, we found that the methylene-linked amide, as in **1i**, harbors the required sweet spot of reactivity and stability for efficient and selective SNAP-tag labeling. Increasing electron density in the phenyl ring results in compounds that still label the SNAP-tag but are unstable in buffer conditions (e.g. **1l**). A reversed amide in the *para* position, as in **1a**, is however feasible and enables conjugation to the SNAP2 ligand in a so far undescribed manner, by a simple amide coupling reaction to the SNAP-targeting phenyl core component. In general, we found that substitutions at the *para* position (**1a, 1i, 1l**) afford highest SNAP labeling, however substitutions at the *meta* position are also well tolerated (e.g. **1b**) and could potentially be explored for the generation of PROTACs.

### PROTACs with an improved SNAP-targeting ligand

To improve the lead **VHL-SNAP1-PROTACs**, we synthesized matched-pairs using the previously reported more cell permeable SNAP2 ligand (**1i**) (**Figure 3A**) (11). SNAP1-and SNAP2-derived PROTACs have similar molecular weight, we did however observe a smaller experimental polar surface area (EPSA) (14 Å^2^ smaller) and higher ChromLogD values (∼1.5 units higher) for the SNAP2-derived PROTACs (n = 4-6 carbons) (**Table S3**). According to emerging guidelines for achieving oral absorption of beyond rule of 5 compounds, size and polarity influence passive cell permeability (45, 46). With this in mind, cell permeability is predicted to be higher for VHL-SNAP2-PROTACs. Indeed, all VHL-SNAP2-PROTACs induced SNAP-EGFP protein degradation at lower concentrations than the matching VHL-SNAP1-PROTACs (**Figure 3B**), while the maximum degradation that could be achieved remained unchanged. A more pronounced hook-effect at lower PROTAC concentration was observed for **VHL-SNAP2-PROTAC-4C** and **-6C**, hinting at a higher intracellular PROTAC concentration, and thus improved cell permeability.

**Figure 3.**
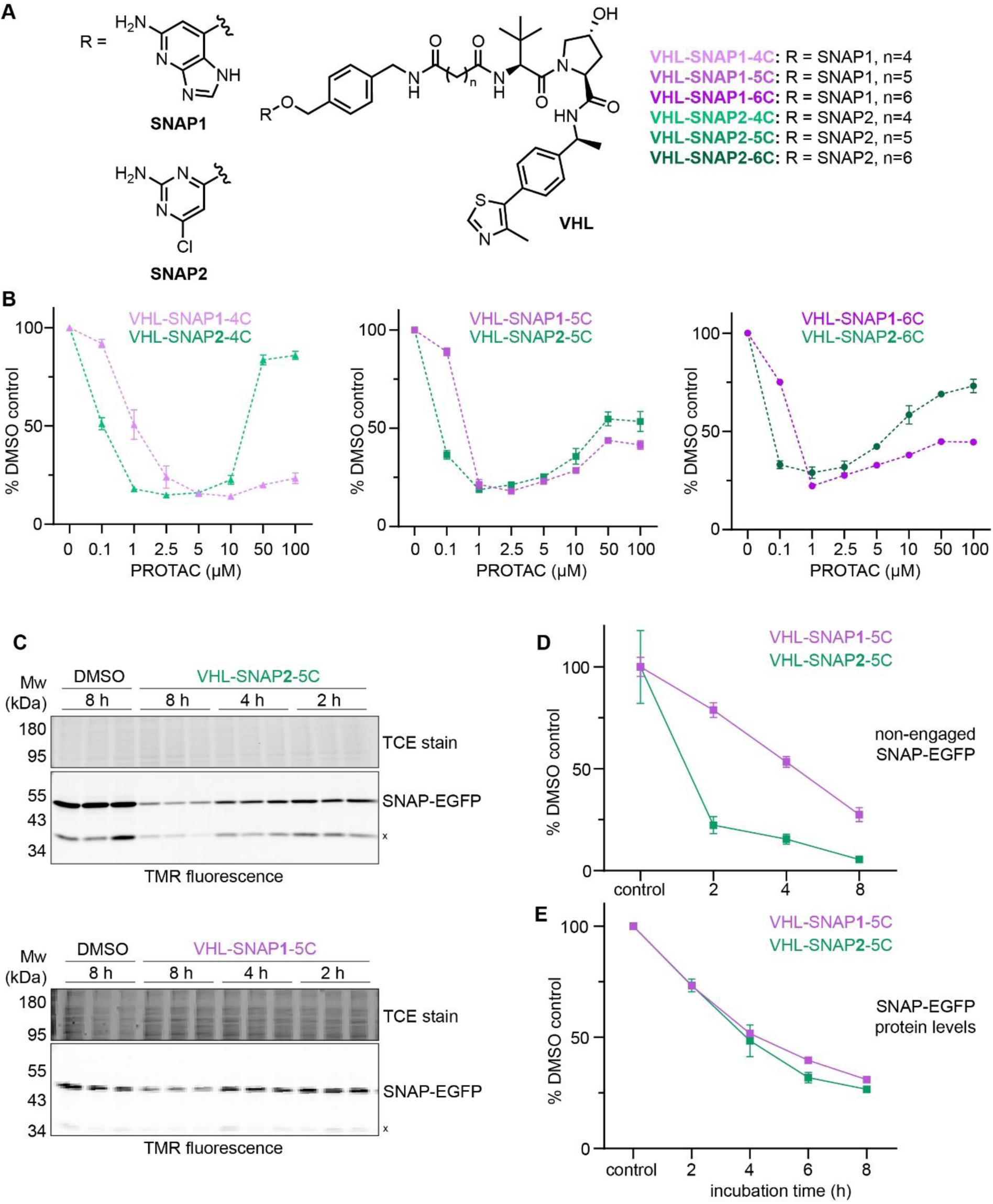
Optimization of VHL-SNAP-PROTACs. (**A**) Chemical structures of PROTACs comprising SNAP1 or SNAP2-ligands, aliphatic linkers, and VHL recruiting ligand. (**B**) Dose response of VHL-SNAP1-or -SNAP2-based PROTACs in HEK293 SNAP-EGFP cells treated for 24 h. SNAP-EGFP levels were quantified by flow cytometry. (n = 3, data represent mean +/- s.d.). (**C**) SDS-PAGE of HEK293 SNAP-EGFP cells after time course with either 1 µM VHL-SNAP2-5C or 2.5 µM VHL-SNAP1-5C. Non-engaged SNAP-EGFP was labeled with SNAP-TMR dye. (**D**) Quantification of non-engaged SNAP-EGFP from (C). (**E**) Quantification of SNAP-EGFP protein levels by flow cytometry after time course treatment with either 1 µM VHL-SNAP2-5C or 2.5 µM VHL-SNAP1-5C. (n = 3, data represent mean +/- s.d.)

For further characterization, we chose the VHL-SNAP1/2-PROTAC-pair containing the 5-carbon linker. By applying a similar strategy, as shown in **Figure 2C** – labeling of unengaged SNAP-EGFP protein with a SNAP-TMR dye – we assessed differences in cell permeability of the PROTACs. **VHL-SNAP2-5C** at 1 µM treatment concentration showed faster target engagement than **VHL-SNAP1-5C** at 2.5 µM (**Figure 3C**), corroborating the higher cell permeability contributed by the SNAP2 ligand (11). However, while over 75% of SNAP-EGFP protein was engaged with **VHL-SNAP2-5C** after 2 h of treatment, degradation showed slower kinetics (**Figure 3D**). In fact, time course analyses of SNAP-EGFP degradation by flow cytometry showed only minor differences between SNAP1-or SNAP2-based VHL-PROTACs (**Figure 3E** and **Figure S3**). For the following experiments, we continued working with the VHL-SNAP2-PROTAC-based compounds, which showed faster SNAP-tag engagement and a trend towards faster degradation.

Accordingly, we developed potent and specific VHL-recruiting PROTACs for the degradation of SNAP fusion proteins. The three VHL-SNAP2-PROTACs (-4C, -5C, -6C) induce efficient degradation of the SNAP-EGFP protein and can be explored in a target-specific manner, when applying this technology to other SNAP-fusion proteins. Notably, the three compounds do not show cytotoxicity up to 50 µM treatment concentration (**Table S3**). Additional data on physicochemical properties of the lead compounds are listed in **Table S3**.

### Development of CRBN-SNAP-PROTACs

To further increase the versatility of SNAP-PROTACs as chemical biology tools, we explored PROTACs recruiting the CRBN E3 ligase. Previous work showed that the target spectrum (27, 36) and cell line-specific activity (47) of PROTACs is E3 ligase-dependent, motivating our efforts to include CRBN-ligands in developing SNAP-PROTACs. We explored different linker lengths and compositions as well as two exit vectors of the CRBN ligand (**Figure 4A** and **S4A**), as these can affect ternary complex formation and hence degradation efficiency (38). Since SNAP2 ligand-based VHL-PROTACs showed faster target engagement (**Figure 3**), we excluded the SNAP1 ligand from our design. From the resulting CRBN-PROTAC series **CRBN5-SNAP2-0C-PIP** and **CRBN5-SNAP2-1C-PIP** showed the most potent degradation of the SNAP-EGFP protein (**Figure 4B** and **S4B**), with a D_max_ of ∼75% at 1 µM PROTAC concentration. At concentrations of 50-100 µM **CRBN5-SNAP2-0/1C-PIP**, cytotoxic effects were obvious from visual inspection of the cells by bright field microscopy. Data on cytotoxicity and physicochemical properties are listed in **Table S4**. We also assessed the kinetics of SNAP-EGFP degradation induced by **CRBN5-SNAP2-0C/1C-PIP** at 1 µM concentration (**Figure 4C**), where both CRBN-PROTACs showed only slightly slower degradation kinetics than the VHL-PROTACs. Hence, in addition to the VHL-recruiting PROTACs, we developed **CRBN-SNAP2-0C/1C-PIP** as efficient tools for TPD of SNAP-fusion proteins.

**Figure 4.**
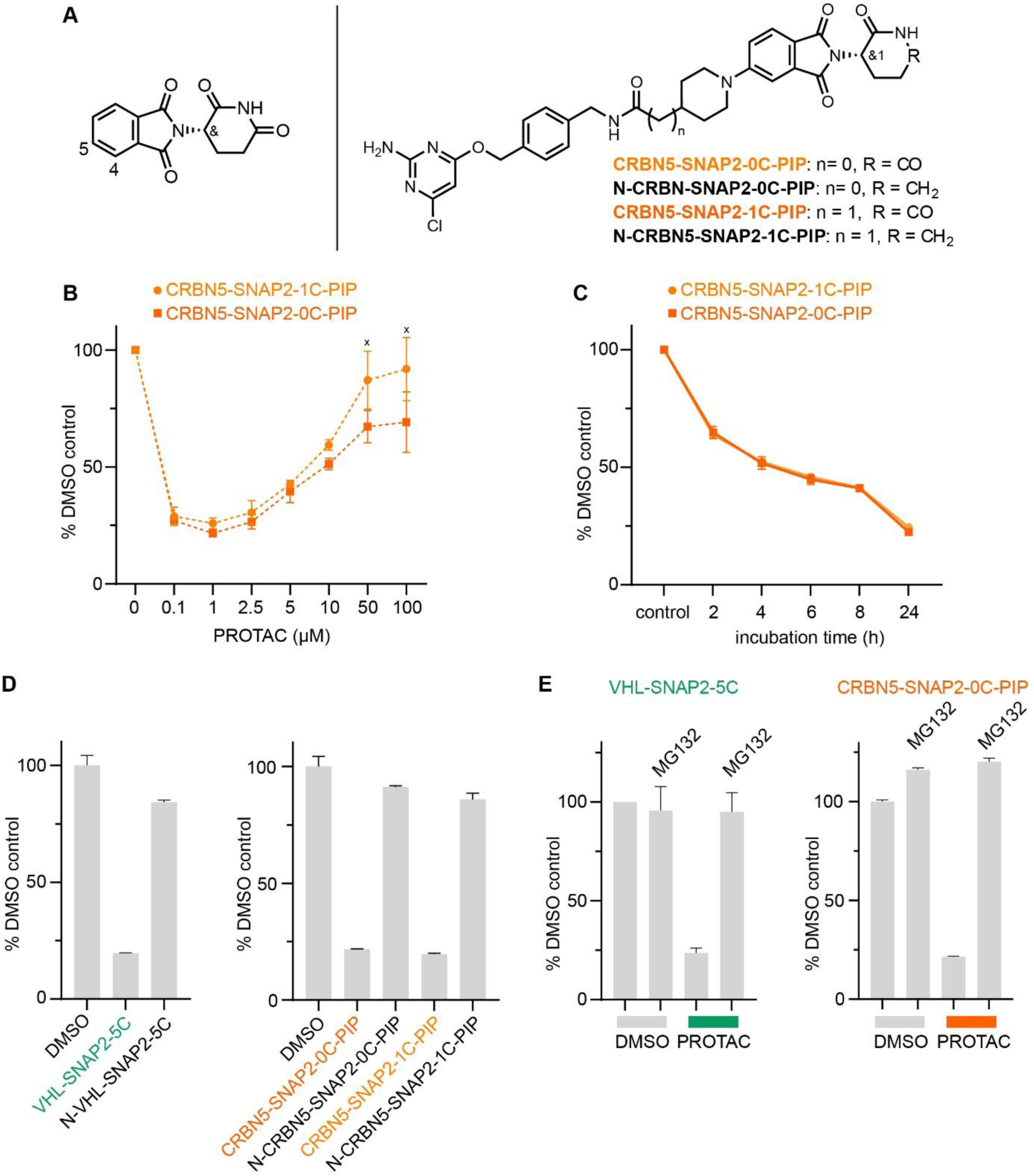
Development of CRBN-recruiting PROTACs and mode of action analysis. (**A**) Chemical structure of thalidomide (left panel) with the positions for the two different exit vector attachment points indicated. Chemical structures of selected CRBN-recruiting PROTACs (right panel). (**B**) Dose response and (**C**) time course analysis of CRBN5-SNAP2-0C/1C-PIP in HEK293 SNAP-EGFP cells. SNAP-EGFP levels were quantified by flow cytometry. (n = 3, data represent mean +/- s.d.). ‘x’ marks concentrations were cell death was observed. (**D**) Comparison of SNAP-EGFP degradation by active PROTACs or control compounds that do not recruit VHL or CRBN. HEK293 SNAP-EGFP cells were treated with 1 µM compound for 24 h and SNAP-EGFP levels were assessed by flow cytometry. (n = 3, data represent mean +/- s.d.). (**E**) Flow cytometry analysis of SNAP-EGFP levels in HEK293 SNAP-EGFP cells treated with 1 µM VHL-SNAP2-5C for 8 h or 1 µM CRBN5-SNAP2-0C-PIP for 24 h. Cells were co-treated with 10 µM MG-132 or DMSO.

### Mode-of-action of VHL-and CRBN-SNAP-PROTACs

To address the mode-of-action of the developed SNAP-PROTACs, we explored whether target degradation is induced by the formation of a ternary complex between the SNAP-PROTAC, the SNAP-fusion protein, and the ubiquitin E3 ligase. We first confirmed that all lead SNAP-PROTACs still bind the respective VHL and CRBN E3 ligases *in vitro* (**Table S3** and **S4**). To determine the PROTAC mode-of-action in cells, we focused on **VHL-SNAP2-5C** and **CRBN5-SNAP2-0C/1C-PIP**. To study the dependence of PROTAC-induced degradation on E3 ligase engagement, we synthesized control compounds that do not engage the VHL or CRBN E3 ligases. We used the previously reported epimer of the VHL ligand to generate an inactive analogue of the VHL-PROTAC (**N-VHL-SNAP2-5C**) (**Figure 3A**) (48, 49). For the CRBN negative control compounds (**N-CRBN5-SNAP2-0C/1C-PIP**, **Figure 4A**), we used a warhead, where CRBN binding is impaired by exchanging the imide for an amide. This exchange results in a ∼1000-fold increase in IC_50_ values in TR-FRET experiments (**Table S4**). All control compounds at 1 µM concentration induced only a minor reduction in SNAP-EGFP levels at the 24 h treatment time point, demonstrating that they are largely inactive, compared to the active PROTACs (**Figure 4D**). The activity of the VHL-based PROTAC is also reduced in the presence of free VHL ligand, further corroborating the dependence on the VHL E3 ligase (**Figure S4C**). Finally, we assessed the dependency of SNAP-EGFP degradation on target engagement. We blocked the active site of the SNAP-tag by treating cells with the SNAP2-ligand (**1i**) 30 min prior to addition of 1 µM **VHL-SNAP2-5C**. SNAP-EGFP degradation was completely abolished under these conditions, confirming the dependence on the PROTAC-SNAP-tag interaction (**Figure S4D**). The PROTAC-induced degradation of SNAP-EGFP was also abolished by co-treating cells with the proteasome inhibitor MG132, demonstrating that target degradation is proteasome-dependent (**Figure 4E**). Taken together, these experiments support that the tested VHL-and CRBN-PROTACs engage their respective ubiquitin E3 ligases and the SNAP-fusion protein to induce the proteasomal degradation of the latter.

### A cellular model for clathrin light chain visualization and degradation

To test the applicability of SNAP-PROTACs in depleting endogenously tagged proteins, we generated a CRISPR-Cas9 knock-in cell line expressing endogenously SNAP-tagged clathrin light chain isoform a (SNAP-CLCa^EN^, EN = endogenous) in the haploid human cell line HAP1 (40). In mammalian cells, clathrin-coated vesicles mediate intracellular trafficking in the secretory and endocytic pathways (50, 51). The clathrin complex is a triskelion consisting of three clathrin heavy chains, each bound to one clathrin light chain (52). While the clathrin heavy chains constitute a fundamental structural component of the vesicle coat, the clathrin light chains primarily play regulatory roles (53). Mammalian cells express two types of clathrin light chains: CLCa and CLCb, with CLCa showing higher relative abundance in most tissues (54). Also in HAP1 cells we observed higher levels of CLCa compared to CLCb, as detected by a pan-CLC antibody (**Figure 5A**). We introduced an N-terminal SNAP-tag into the endogenous locus of the CLCa gene (CTLA), thereby generating a fusion protein with ∼55 kDa (**Figure 5A**). We tested the induced degradation of SNAP-CLCa^EN^ with the **VHL-SNAP2-5C** (1 µM) and **CRBN5-SNAP2-PIP-1C** (0.1 µM) PROTACs at a short (2 h) and a long (24 h) treatment time point (**Figure 5BC**). Both PROTACs induced the depletion of SNAP-CLCa^EN^ nearly below detection limit of the Western blot within 24 h (**Figure 5BC**). We continued working with the VHL-recruiting PROTAC, since at the 2 h-time point **VHL-SNAP2-5C** outperformed **CRBN5-SNAP2-PIP-1C**. In a time course experiment, **VHL-SNAP2-5C** induced maximum depletion of SNAP-CLCa^EN^ within 4-6 h (**Figure 5D**).

**Figure 5.**
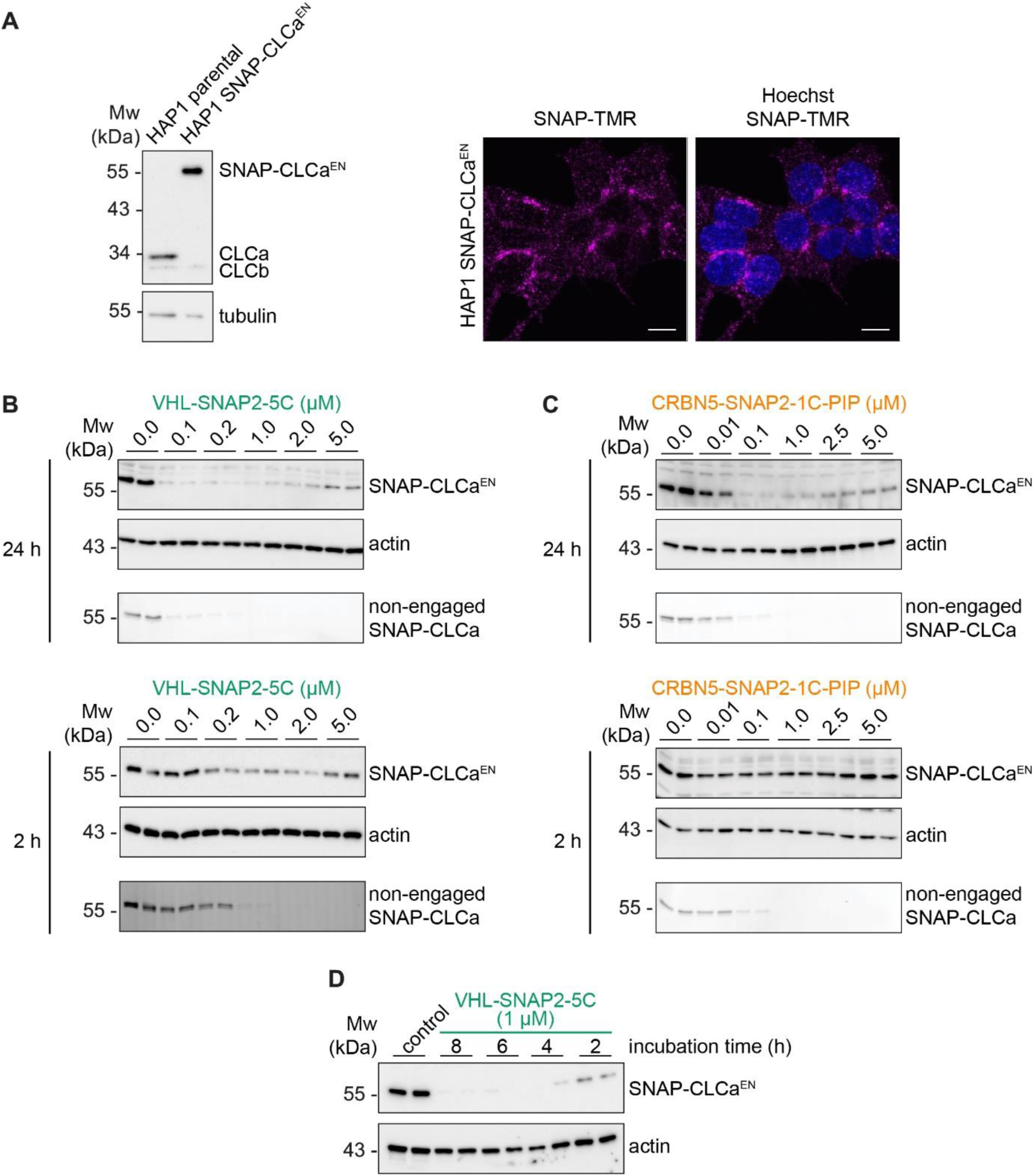
Degradation of endogenously tagged SNAP-CLCa. (**A**) (left) Western blot of HAP1-SNAP-CLCa^EN^ and parental HAP1 cells. CLC signal was detected with a pan-CLC antibody. (right) Confocal images of HAP1 SNAP-CLCa^EN^ cells labeled with SNAP-TMR dye. Scale bars are 10 µm. Dose response of VHL-SNAP2-5C (**B**) or CRBN5-SNAP2-1C-PIP (**C**) in HAP1 SNAP-CLCa^EN^ cells for 24 h (top) and 2 h (bottom). Non-engaged SNAP-CLCa was assessed with TMR-labeling. (**D**) Time course analysis of HAP1 SNAP-CLCa^EN^ cells treated with 1 µM VHL-SNAP2-5C.

Based on this initial characterization, we performed a quantitative mass spectrometry experiment of SNAP-CLCa^EN^ HAP1 cells upon 6 h and 24 h treatment with 1 µM **VHL-SNAP2-5C**. The proteomics data showed a ∼3-fold decrease of SNAP-CLCa levels after 6 h and ∼6-fold decrease after 24 h PROTAC treatment (**Figure 6**). In contrast, the levels of the clathrin heavy chain and of CLCb are not significantly affected by the PROTAC treatment. At the 24 h treatment time point, we observed an increase in levels of the endoplasmic reticulum (ER) protein quality control factors SEC61G, Derlin, and UBQLN2, indicating that long-term depletion of CLCa may affect (ER) protein homeostasis. We also observed higher levels for the Prefoldin chaperone subunits -1, -5, and -6. The Prefoldin complex is a hetero-hexameric chaperone important for the quality control of the cytoskeleton proteins actin and tubulin (55, 56). Clathrin light chains connect clathrin-coated vesicles to the actin cytoskeleton (57–59). Previous work further established that depleting CLCa and CLCb by siRNA leads to actin overassembly and accumulation in patches (60). In imaging experiments, we assessed the integrity of the actin cytoskeleton upon PROTAC-mediated depletion of SNAP-CLCa^EN^ for 24 h using phalloidin (**Figure S5**). In these experiments, we did not observe any bulk structural differences in the actin cytoskeleton upon SNAP-CLCa^EN^ degradation, suggesting that depletion of both clathrin light chains is required for a phenotype to manifest in imaging experiments.

**Figure 6.**
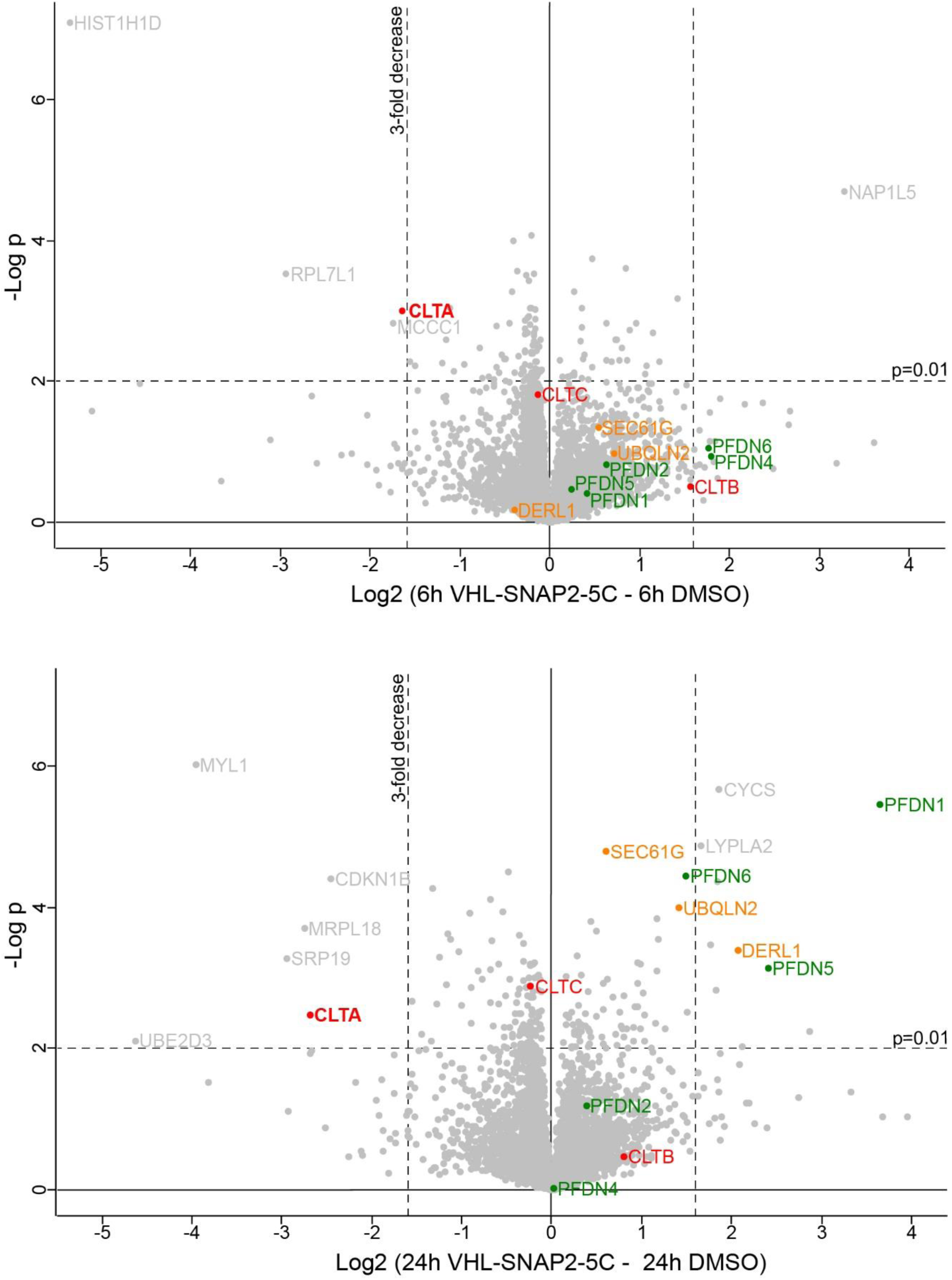
Characterization of SNAP-CLCa^EN^ depletion. Mass spectrometry analysis of cell lysates from HAP1 SNAP-CLCa^EN^ cells treated with 1 µM VHL-SNAP2-5C for 6 h or 24 h. Volcano plots show effect of VHL-SNAP2-5C on protein levels in HAP1 SNAP-CLCa^EN^ cells relative to DMSO control. Each experiment with four technical replicates, each data point represents mean value. Statistical significance was determined using a two-sided Student’s t test.

## DISCUSSION

The use of SNAP-fusion proteins has facilitated the characterization of a multitude of POIs. In this work, we targeted a component of the clathrin vesicle coat in mammalian cells, by generating a CRISPR-Cas9 knock-in cell line expressing an endogenously SNAP-tagged CLCa fusion protein. As we could show for SNAP-CLCa^EN^, an engineered POI can easily be visualized with SNAP-dyes and depleted with SNAP-PROTACs (**Figure 5** and **6**). SNAP-dyes are suitable for super-resolution microscopy and allow tracking vesicles and tubules coated with SNAP-CLCa^EN^ with high precision (13). Depletion experiments can easily be applied in the same cell lines, provided the SNAP-tag is homozygously integrated. Interfering with transport processes in the secretory and endocytic pathway by depletion of vesicle coat proteins or regulatory proteins is an important strategy to understand the structural and regulatory roles of POIs. As vesicle transport processes are fast-paced, acute depletion of coat components and regulatory factors is preferred over long-term genetic knock-out approaches. With our TPD set-up, we identified differences between short-term and long-term depletion of CLCa, an approach not feasible with a genetic knock-out strategy. Specifically, our proteomics data highlight differences between depletion of SNAP-CLCa^EN^ from HAP1 cells with **VHL-SNAP2-5C** for 6 h and 24 h. While at the 6 h time point we did not find any group of proteins significantly increased, we observed higher protein levels of Prefoldin chaperone subunits at the 24 h time point, hinting at an effect on the cytoskeleton upon CLCa depletion. Prefoldin, a multi-subunit chaperone, facilitates the folding of actin and tubulin subunits (55, 56). Its upregulation is in line with previous work describing a role for clathrin light chains in regulating local actin assembly (60). Our imaging data show that depletion of CLCa is not sufficient to induce previously observed phenotypes of actin cytoskeleton disorganization, yet we anticipate that the established SNAP-CLCa depletion system when applied in polarized cell lines could help define non-redundant roles of CLCa and CLCb.

Our experimental workflow for SNAP-CLCa^EN^ serves as a blueprint for applying SNAP-PROTACs to endogenously tagged SNAP-fusion proteins. In our example, we used **VHL-SNAP2-5C** to induce efficient SNAP-CLCa^EN^ degradation. However, we anticipate target-specific differences in SNAP-PROTAC-induced degradation, as observed for other induced degradation systems leveraging protein tags (27, 61). To this end, we developed three VHL-and two CRBN-recruiting SNAP-PROTACs that can be screened to evaluate the most efficient target degradation. This evaluation can be carried out with an overexpressed SNAP-fusion protein, prior to, or in parallel to the generation of a homozygous knock-in cell line, an experimental workflow that is typically more time-consuming. In this work, we used HEK293 and HAP1 cell lines, however we expect the SNAP-PROTAC approach to be widely applicable, since the VHL and CRBN E3 ligases are active in a wide range of cell lines (47).

A limitation of SNAP-PROTACs may be their covalent nature, as they cannot act in a catalytic manner, thus requiring stoichiometric PROTAC concertation to induce efficient degradation. However, they represent an efficient means to easily integrate loss-of-function experiments with SNAP-fusion proteins. The SNAP-tag now combines access to protein visualization, isolation, proximity biotinylation, and TPD in a single protein tag and thereby greatly reduces genetic engineering efforts that are often time-and cost-intensive. In general, the introduction of SNAP-PROTACs extends the tag-PROTAC systems and it may even be combined with previously described TPD approaches targeting the dTag, eDHFR, the BromoTag, or the HaloTag, as it is orthogonal to these (27–32). Previous work also highlights target-specific differences in the ability of tag-targeting PROTACs to induce the degradation of fusion proteins (27, 61). To this end, SNAP-PROTACs may offer access to targets which are currently difficult to degrade with existing tag-PROTAC systems.

One future directive to further expand the applicability of SNAP-PROTACs to different targets is the use of alternative SNAP-targeting ligands as recruiting elements. To this end, we identified additional SNAP-ligands, which conjugate to the SNAP-tag in cells (**Figure 2**). The activity of the reference compound **1i** and the top hits, **1a** and **1l**, were predicted based on FEP calculations. Contradictory to the FEP calculations, compound **1c** and **1d**, were inactive in cells. One explanation for the discrepancy could be that FEP calculations only consider the formation of the protein-substrate complex and do not take into account the stabilization of the transition states by the protein. FEP calculations further only take into account the binding affinity (i.e. K_D_), but not the kinetics of the enzymatic conjugation reaction. Taken together, we found that SNAP ligands carrying a reversed amide (**1a**) or an alkyne (**1g**) in the *para* position are well tolerated. We further showcased a larger scope of tolerated substitutions in the *meta* position, previously unexplored (**1b**, **1h**, **1j**, and **1m**, **Figure 2B**). All of these diverse functional handles can be explored in future design of SNAP-targeting probes and specifically in the design of SNAP-PROTAC.

## Methods

### Cloning

SNAP-HA-EGFP (SNAP-EGFP) was cloned into the multiple cloning site of pcDNA5/FRT (Invitrogen) using the HindIII and NotI restriction sites. The SNAP-tag was amplified from B4GT-SNAP-HT2 in pcDNA5/FRT/TO (62), HA-EGFP was amplified from Halo-HA-EGFP in pcDNA5/FRT (31) using accordingly designed primers. Upon restriction digest the two PCR fragments and the vector were ligated. To create CLIP-HA-EGFP (CLIP-EGFP) in pcDNA5/FRT, the SNAP-tag was removed from SNAP-HA-EGFP in pcDNA5/FRT by restriction digest with NheI and replaced with the CLIP-tag using the NEBuilder HiFi DNA Assembly Cloning Kit (NEB) with accordingly designed primers. The CLIP-tag sequence was amplified from EGFP-CLIP in pcDNA5/FRT, a kind gift from Kai Johnsson.

### Cell culture

HAP1 cells (63) were cultured in DMEM (Thermo Fisher Scientific) supplemented with 10% fetal bovine serum (FBS; Thermo Fisher Scientific), 100 U/ml penicillin and 100 μg/ml streptomycin (Invitrogen), referred to as growth medium. Parental HEK293 Flp-In T-REx cells (Thermo Fisher Scientific) were cultured in growth medium containing 100 µg/ml zeocin (Invivogen) and 15 µg/ml blasticidin (Invivogen). Cells were kept at 37 °C and 5% CO2 and regularly tested for mycoplasma infection using Venor^®^GeM OneStep mycoplasma detection kit (Minerva Biolabs). Stable HEK293 Flp-In cell lines constitutively expressing SNAP-EGFP or CLIP-EGFP were generated using the Flp-In T-REx system (Thermo Fisher Scientific) in combination with the pcDNA5/FRT vector according to the manufacturer’s protocol. Stable HEK293 SNAP-EGFP and CLIP-EGFP cells were selected and cultured in growth medium supplemented with 100 µg/ml hygromycin B (Invivogen).

### Generation of gene-edited SNAP-CLCa^EN^ knock-in cells

The guide RNA for targeting the CLTA gene (gene ID 1211) and the homology repair (HR) plasmid for SNAP-CLCa were previously described in (13). Transfection of guide RNA and HR plasmids into HAP1 cells was done using a NEPA21 electroporation system (Nepa Gene). For electroporation of HAP1 cells, 3 million cells were washed twice with Opti-MEM (Gibco) and resuspended in 90 µl Opti-MEM with 10 µg of DNA in an electroporation cuvette with a 2 mm gap. HAP1 cells were electroporated with a poring pulse of 200 V, 5 ms pulse length, 50 ms pulse interval, 2 pulses, with decay rate of 10% and + polarity; consecutively with a transfer pulse of 20 V, 50 ms pulse length, 50 ms pulse interval, 5 pulses, with a decay rate of 40% and ± polarity. To select for positive clones after electroporation, HAP1 cells were treated with 1 µg/ml puromycin (Gibco). After selection of positive clones, HAP1 cells were further sorted based on cell size and fluorescence to obtain homogenous population of edited haploid cells (64).

### In lysate labeling

Cultured cells were trypsinised, resuspended in growth medium and collected by centrifugation at 1.000 g for 5 min at RT. Pellets were washed with PBS before resuspension in lysis buffer (50 mM Tris pH 7.5, 100 mM NaCl, 0.1% Tween20, 1x protease inhibitor cocktail (Roche)). Lysed cells were incubated with 1.5 µM SNAP-TMR dye for 30 min at RT and then mixed in a 1:1 ratio with 100 mM Tris pH 7.5, 300 mM NaCl, 2 mM EDTA, 0.5% SDS, 2% NP-40, 1% Triton X-100, 1x protease inhibitor cocktail (Roche) and incubated on ice for 15 min. Next, the cells were spun down at 20.000 g for 10 min, the supernatant was collected and the protein concentration determined using the Pierce^TM^ BCA Protein Assay Kit (Thermo Scientific). The remaining supernatant was mixed with 4x SDS sample buffer (200 mM Tris pH 6.8, 8% SDS, 40% glycerol, 4% β-mercaptoethanol, bromophenol blue) and incubated at 95 °C for 5 min before analysis by SDS-PAGE. SDS-PAGE gels were supplemented with 0.5 % 2,2,2-Trichloroethanol (TCE) for visualization of proteins following electrophoresis. TCE and SNAP-TMR signals were detected with the stain-free (UV light) and Cy3 settings of the ChemiDoc MP imaging system (BioRad), respectively. Band quantification from SDS-PAGE gels was performed with Image Lab (BioRad). TMR signals were normalized to TCE signals as loading controls. Data were plotted and analysed in Prism (GraphPad).

### Western blot

Cells were harvested on ice in lysis buffer (50 mM Tris pH 7.5, 150 mM NaCl, 1 mM EDTA, 0.25% SDS, 1% NP-40, 0.5% Triton X-100, 1x protease inhibitor cocktail (Roche)). Lysates were centrifuged for 10 min at 20.000 g at 4 °C and further processed as described above. Protein extracts were applied to SDS-PAGE gels followed by transfer to a nitrocellulose membrane for 90 min at 75 V. Membranes were blocked in 5% non-fat dry milk in TBS supplemented with 0.1% Tween-20 for 1 h prior to incubation with the primary antibody dilutions. Membranes were probed for CLC (Merck, Cat#AB9884, 1:000), actin (Santa Cruz Biotechnology, Cat#sc-47778, 1:1000), or tubulin (Sigma-Aldrich, Cat#T9026, 1:10.000). Incubation was performed overnight at 4 °C and followed by incubation with HRP-conjugated secondary antibodies diluted 1:20.000 in TBS supplemented with 0.1% Tween-20 (anti-mouse-HRP, Invitrogen, Cat#31444 or anti-rabbit-HRP, Invitrogen, #31460). Secondary antibodies were incubated for 45 min at RT before detection at the ChemiDoc MP imaging system (BioRad) using ECL start (Cytiva) or ECL prime (Cytiva) western blotting detection reagents. Band intensities were analyzed using Image Lab (BioRad) and further processed using Prism (Graphpad).

### Flow cytometry

Cultured cells were trypsinised, resuspended in growth medium and collected by centrifugation at 1.000 g for 5 min. Growth medium was removed and cells were resuspended in PBS, 1% FBS, 1 mM EDTA and transferred to a 96-well plate. Cells were flowed at the Miltenyi MACSQuant VYB Flow Cytometer, measuring 20.000 events per sample. Flow cytometry data (fcs-files) were analysed in Kaluza (Beckmann Coulter) and plotted in Prism (Graphpad).

### Fluorescence microscopy

For microscopy experiments cells were grown on fibronectin coated coverslips and, after the indicated treatment, fixed in 4% paraformaldehyde in PBS. Cells were permeabilized, and blocked in 10% FBS in PBS + 0.01% Triton X-100, for 45 min at RT, followed by incubation with Alexa Fluor-633 Phalloidin (Invitrogren, Cat#A22284,1:2000) and Hoechst for 45 min at RT. Coverslips were mounted on microscopy slides in Vectashield mounting medium (Vector Laboratories). Images were acquired at a Leica TCS SP8 HCS A using a 63x oil objective and processed using Fiji (65).

### Sample preparation for LC/MS

Single-Pot Solid-Phase-enhanced Sample Preparation (SP3) for LC/MS sample preparation for LC/MS/MS is based on the SP3 protocol (66). 15 µg protein extracts were taken up in 100 µL 1× SP3 lysis buffer (final concentrations: 1% (wt/vol) SDS; 10 mM TCEP; 200 μL 40 mM chloracetamide; 250 mM HEPES pH 8) and heated for 5 min at 90 °C. After cooling the samples to room temperature 7 units Benzonase (Merck) were added to each sample and incubated for 30 min at 37°C. Next the Benzonase treated samples were mixed with 150 µg hydrophobic (#65152105050250) and 150 µg hydrophilic (#45152105050250) SeraMag Speed Beads (Cytiva) (bead to protein ratio 10 to 1) and gently mixed. Then 100 µL 100% vol/vol Ethanol (EtOH) was added before incubation for 20 min at 24°C shaking vigorously. The beads were collected on a magnet and the supernatant aspirated. The beads were then washed 4 times with 180 µL 80 % EtOH (collection time on the magnet minimum of 4 min). The beads were finally taken up in 100 µL 25 mM ammoniumbicarbonate (ABC) containing 1 µg Trypsin (Protein:Trypsin ratio 30:1). To help bead dissociation, samples were incubated for 5 min in a sonification bath (preheated to 37°C). Samples were incubated overnight, shaking vigorously (1300 rpm). Next day samples were acidified with formic acid (FA, final 1% vol/vol) before collection on a magnet. The supernatants were transferred to a fresh Eppendorf tube, before removing trace beads using a magnet for 5 min. The tryptic digests were then desalted on home-made C18 StageTips as described (67). Briefly, peptides were immobilized and washed on a 2 disc C18 StageTip. Samples were then dried using a vacuum concentrator (Eppendorf) and the peptides were taken up in 0.1% formic acid solution (10 μL) and directly used for LC-MS/MS experiments.

## LC-MS/MS

Experiments were performed on an Orbitrap Fusion Lumos (Thermo) that was coupled to an EASY-nLC 1200 liquid chromatography (LC) system (Thermo). The LC was operated in the one-column mode. The analytical column was a fused silica capillary (75 µm × 41 cm) with an integrated frit emitter (CoAnn Technologies) packed in-house with Kinetex C18-XB core shell 1.7 µm resin (Phenomenex). The analytical column was encased by a column oven (Sonation) and attached to a nanospray flex ion source (Thermo). The column oven temperature was adjusted to 50 °C during data acquisition. The LC was equipped with two mobile phases: solvent A (0.2% formic acid, FA, 99.9% H_2_O) and solvent B (0.2% formic acid, FA, 80% Acetonitrile, ACN, 19.8% H_2_O). All solvents were of UPLC grade (Honeywell). Peptides were directly loaded onto the analytical column with a maximum flow rate that would not exceed the set pressure limit of 980 bar (usually around 0.5 – 0.7 µL/min). Peptides were subsequently separated on the analytical column by running a 105 min gradient of solvent A and solvent B (start with 3% B; gradient 3% to 9% B for 6:30 min; gradient 9% to 30% B for 62:30 min gradient 30% to 50% B for 24 min; gradient 50% to 100% B for 2:30 min and 100% B for 9:30 min) at a flow rate of 300 nl/min. The mass spectrometer was operated using Tune v3.3.2782.28. The mass spectrometer was set in the positive ion mode. Precursor ion scanning was performed in the Orbitrap analyzer (FTMS; Fourier Transform Mass Spectrometry) in the scan range of *m/z* 300-1500 and at a resolution of 240000 with the internal lock mass option turned on (lock mass was 445.120025 *m/z*, polysiloxane) (68) Product ion spectra were recorded in a data dependent fashion in the ITMS at “rapid” scan rate. The ionization potential (spray voltage) was set to 2.5 kV. Peptides were analyzed using a repeating cycle consisting of a full precursor ion scan (AGC standard; max acquisition time “Auto”) followed by a variable number of product ion scans (AGC 300% and acquisition time auto) where peptides are isolated based on their intensity in the full survey scan (threshold of 4000 counts) for tandem mass spectrum (MS2) generation that permits peptide sequencing and identification. Cycle time between MS1 scans was 3 sec. Fragmentation was achieved by stepped Higher Energy Collision Dissociation (sHCD) (NCE 27, 32, 40). During MS2 data acquisition dynamic ion exclusion was set to 20 seconds and a repeat count of one. Ion injection time prediction, preview mode for the FTMS, monoisotopic precursor selection and charge state screening were enabled. Only charge states between +2 and +7 were considered for fragmentation.

### Peptide and Protein Identification using MaxQuant

RAW spectra were submitted to an Andromeda (69) search in MaxQuant (2.0.2.0.) using the default settings (70) Label-free quantification and match-between-runs was activated (71). The MS/MS spectra data were searched against the Uniprot *H. sapiens* reference proteome (ACE_0719_UP000005640_9606_full.fasta; 79071 entries) where the original P09496|CLCa sequence was replaced by the SNAP-CLCa version used in this project. All searches included a contaminants database search (as implemented in MaxQuant, 245 entries). The contaminants database contains known MS contaminants and was included to estimate the level of contamination. Andromeda searches allowed oxidation of methionine residues (16 Da) and acetylation of the protein N-terminus (42 Da). Carbamidomethylation on Cysteine (57) was selected as static modification. Enzyme specificity was set to “Trypsin/P”. The instrument type in Andromeda searches was set to Orbitrap and the precursor mass tolerance was set to ±20 ppm (first search) and ±4.5 ppm (main search). The MS/MS match tolerance was set to ±0.5 Da. The peptide spectrum match FDR and the protein FDR were set to 0.01 (based on target-decoy approach). For protein quantification unique and razor peptides were allowed. Modified peptides were allowed for quantification. The minimum score for modified peptides was 40. Label-free protein quantification was switched on, and unique and razor peptides were considered for quantification with a minimum ratio count of 2. Retention times were recalibrated based on the built-in nonlinear time-rescaling algorithm. MS/MS identifications were transferred between LC-MS/MS runs with the “match between runs” option in which the maximal match time window was set to 0.7 min and the alignment time window set to 20 min. The quantification is based on the “value at maximum” of the extracted ion current. At least two quantitation events were required for a quantifiable protein. Further analysis and filtering of the results was done in Perseus v1.6.10.0. (72). Comparison of protein group quantities (relative quantification) between different MS runs is based solely on the LFQs as calculated by MaxQuant, MaxLFQ algorithm (71).

### Free Energy Calculations

FEP calculations were conducted on a series of analogs of the SNAP2 ligand with diverse substitutions in the phenyl ring. The unsubstituted benzyl chloropyrimidine was chosen as the reference ligand due to its shared substructure with all the assessed molecules. For compounds capable of binding in two different orientations due to a 180-degree rotation of a pyridine ring or a phenyl ring with a meta-substituent (1b, 1d, 1f, 1h, 1j, and 1m), both conformations were simulated. Throughout the simulations, all ligands exhibited low root-mean-square deviation (RMSD) values relative to their initial conformations.

Protein structures were retrieved from the Protein Data Bank (PDB) at www.rcsb.org and subsequently imported into Maestro (Schrödinger Release 2023-4: Schrödinger, LLC: NewYork, NY, 2023). Protein Preparation Wizard in Maestro were used to process these structures. LigPrep with Epik were used for assigning protonation states (73). To generate 3D conformations within the binding site, Glide core-constrained docking, based on the co-crystallized ligand as a reference, was executed (74). Poses resulting from this procedure underwent visual inspection. In cases where multiple potential binding modes were identified (especially for molecules with a single m-substitution), the second symmetrical conformation was manually generated by adjusting the corresponding torsion.

Prospective free energy calculations were subsequently carried out using the Schrödinger FEP+ method within Schrödinger Suite version 2023-4 (75). The OPLS4 forcefield, using the custom parameters generated via the Force Field Builder, was employed (76). Calculations adhered to default settings, with the sampling extended to 10 ns per λ-window and a total of 48 λ-windows per transformation. Optimal topology perturbation map was biased towards the ligand with the unsubstituted phenyl ring. The results of calculations were analyzed using the FEP+ GUI.

## Supporting information

Supplemental files combined

## Data availability

The mass spectrometry proteomics data have been deposited to the ProteomeXchange Consortium via the PRIDE (77) partner repository (https://www.ebi.ac.uk/pride/archive/) with the dataset identifier PXD049283.

## Acknowledgements

We thank Lars Borgards, Tim Vanselow, Yannick Schnurbusch, Lea Robin Krift, Anne Derichs, and Lorena Lindermann for support of cell biology experiments. We also thank Svenja Heimann and Jenny Bormann for help with sample preparation for mass spectrometry. We thank Malin Lemurell for supporting the project. This work was funded by the Sofja Kovaleveskaja Award by the Alexander von Humboldt Foundation endowed by the Federal Ministry of Education and Research and by the Deutsche Forschungsgemeinschaft (DFG, German Research Foundation) – SFB1430 – Project-ID 424228829 (sub-projects: A07 (D.H.), B01 (M.K.), Z03 (F.K.)). L.W.D.’s work is funded by the DFG – Project-ID 278001972 – TRR 186 and Freie Universität Berlin. We acknowledge the Imaging Center Campus Essen (ICCE), Center of Biotechnology (ZMB), University of Duisburg-Essen, for providing the imaging equipment and support in microscope usage and image analysis. Leica TCS SP8 HCS A microscope was funded by the Deutsche Forschungsgemeinschaft (DFG, German Research Foundation) – Project-ID 219183055

## Author contributions

S.A-P. and G.S. performed cell biology experiments. J.B., S.L., D.L.P. and M.P. performed chemical synthesis. F.K. and M.K. performed MS experiments. L.W.D. and F.B. engineered the HAP1 cell line. V.P.B. performed FEP calculations. M.P. and D.H. conceived, outlined, and supervised the project. M.P. and D.H. prepared the manuscript with the help of G.S., S.L., and V.P.B., and input from all co-authors.

## Key resources table

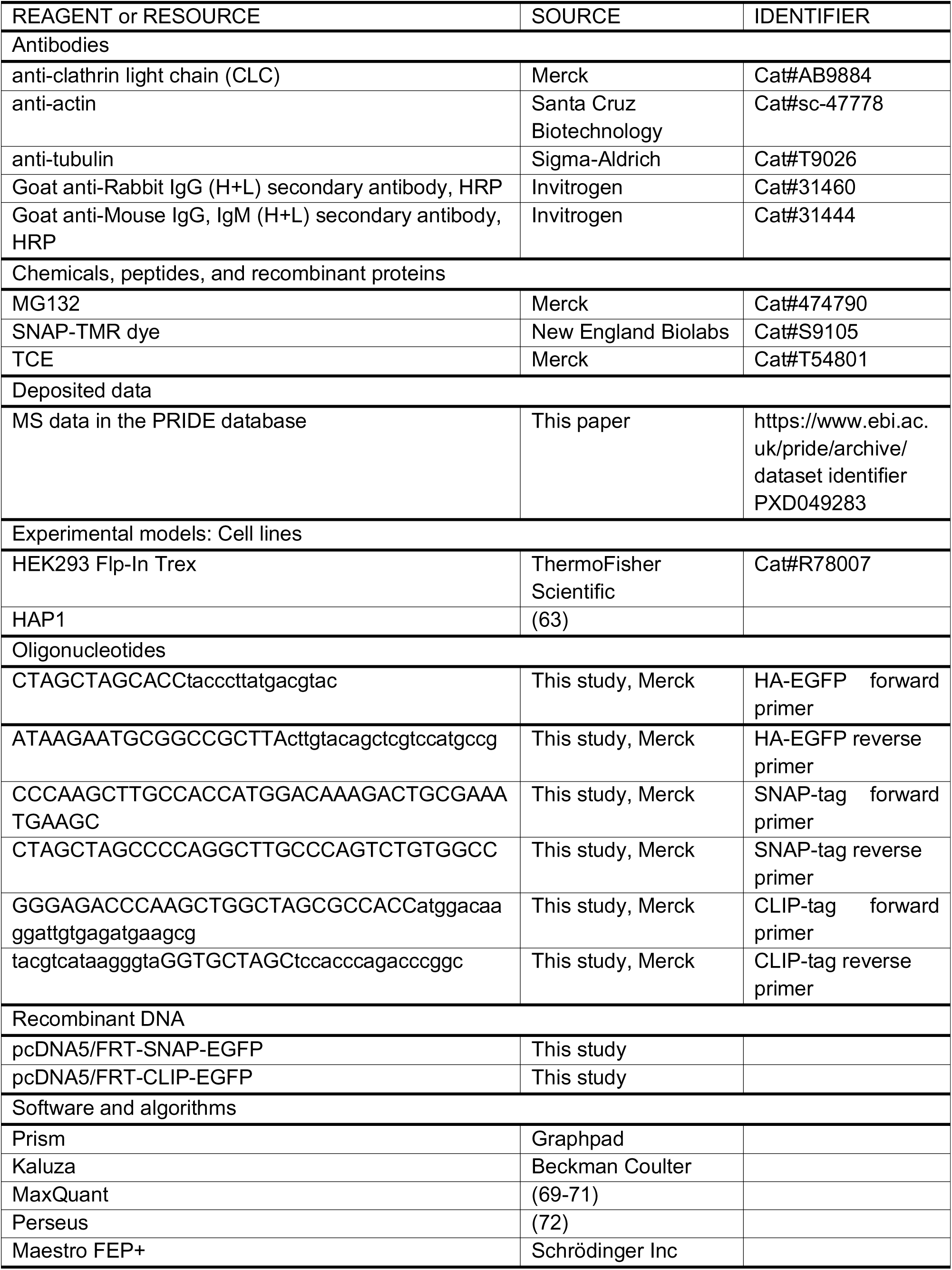

