## Supplemental files combined for "Induced degradation of SNAP-fusion proteins"

**Content:** Supplementary Figures 1-4, Supplementary Tables 1-3, Scheme S1-S6,Synthetic protocols for described compounds

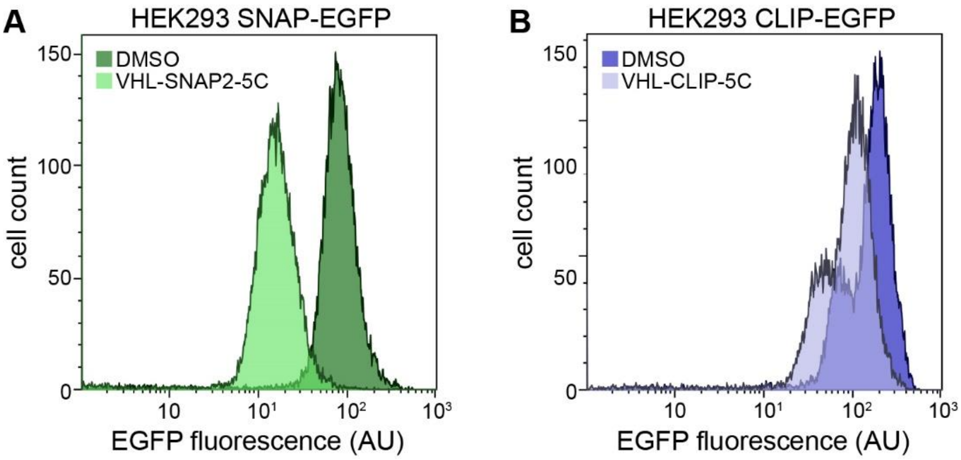


**Figure S1. Characterization of stable HEK293 SNAP-EGFP and CLIP-EGFP cell lines.** (**A**) Flow cytometry analysis (histograms) of HEK293 SNAP-EGFP cells treated with 1 µM VHL-SNAP2-5C or DMSO for 24 h. (**B**) Flow cytometry analysis (histograms) of HEK293 CLIP-EGFP cells treated with 5 µM VHL-CLIP-5C or DMSO for 24 h.

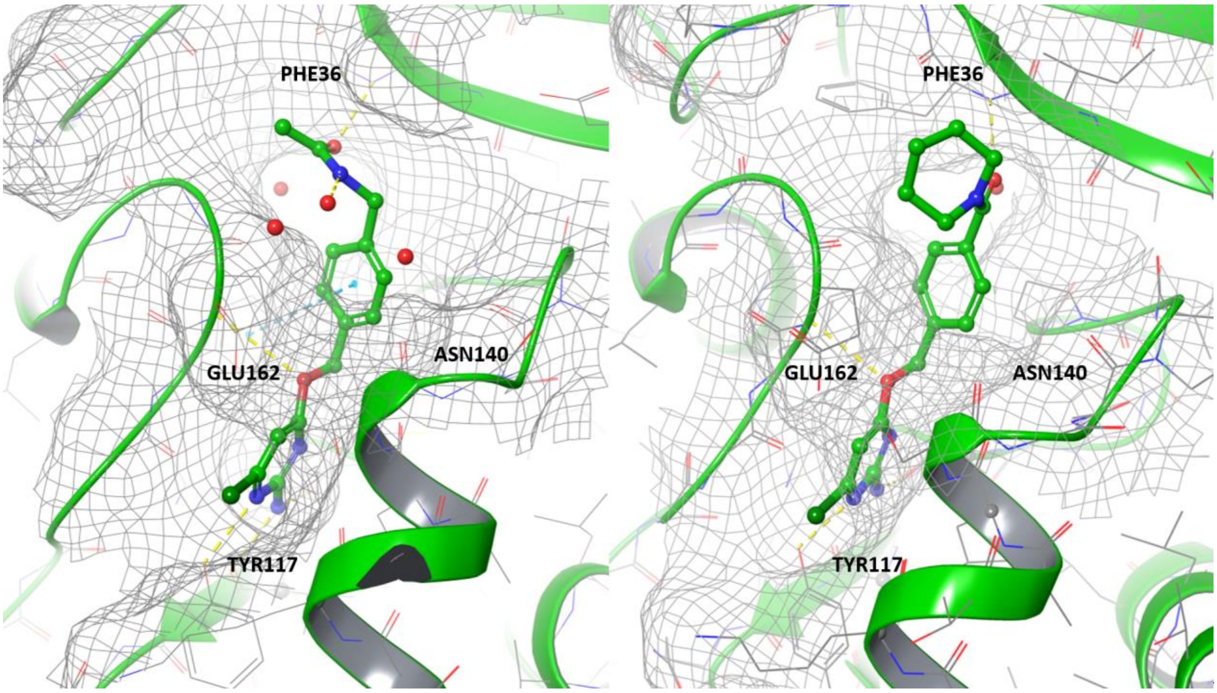


**Figure S2.** Representative conformations of the ligand **1i** (left) and **1c** (right) bound to SNAP protein. Showing that piperidine of ligand 1c is solvent exposed.

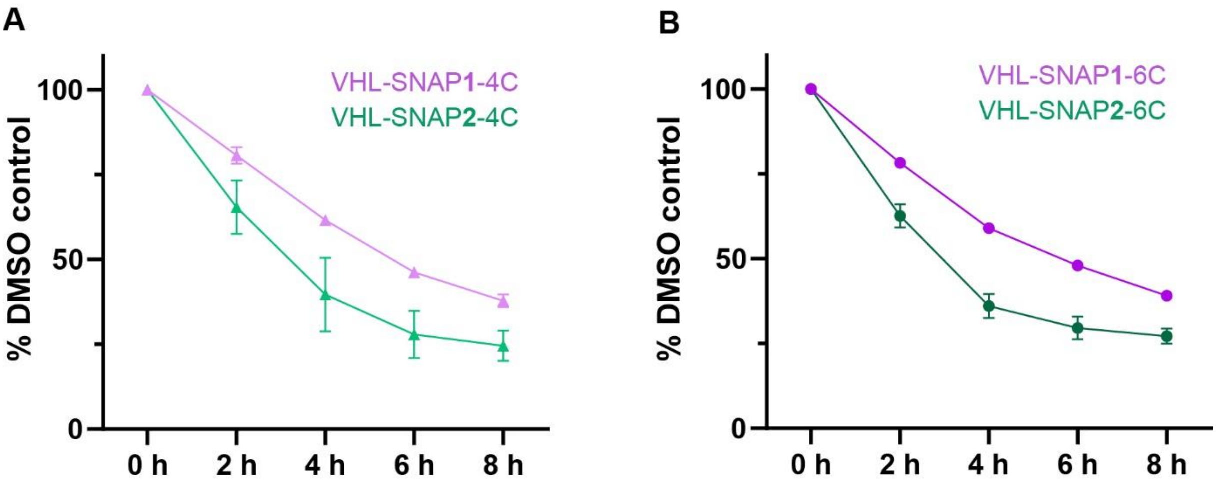


**Figure S3. Time course analysis of EGFP-SNAP degradation upon VHL-SNAP1/2-PROTAC treatment.** (**A**) Time course analysis of HEK293 SNAP-EGFP cells treated with 5 µM VHL-SNAP1-4C or 1 µM VHL-SNAP2-4C, concentrations at which maximal degradation was observed at 24 h. SNAP-EGFP degradation was measured by flow cytometry. (**B**) Time course analysis of HEK293 SNAP-EGFP cells treated with 1 µM VHL-SNAP1-6C or VHL-SNAP2-6C, the concentration at which maximal degradation was observed at 24 h. SNAP-EGFP degradation was measured by flow cytometry. (n = 3, data represent mean +/- s.d.).

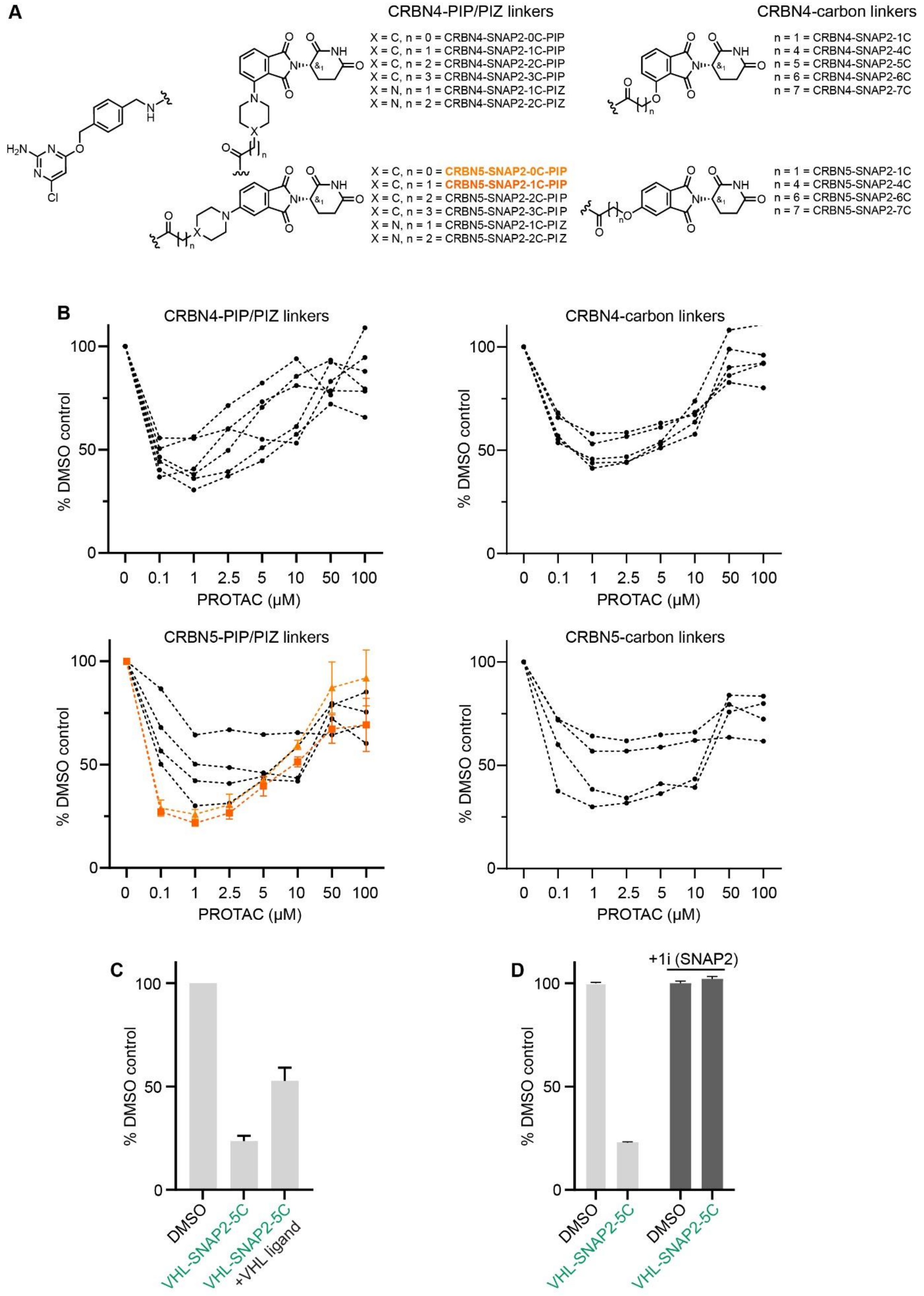


**Figure S4. Screening of CRBN-SNAP2-PROTAC series.** (**A**) Chemical structures of CRBN-SNAP2-PROTACs. (**B**) Dose response of SNAP2-based CRBN-recruiting PROTACs. HEK293 SNAP-EGFP cells were treated for 24 h with the indicated PROTACs and SNAP-

EGFP degradation was assessed by flow cytometry. (**C**) HEK293 SNAP-EGFP cells were treated with 1 µM VHL-SNAP2-5C for 8 h with or without additional pre-treatment with 10 µM VHL-ligand for 30 min. SNAP-EGFP levels were assessed by flow cytometry. (**D**) HEK293 SNAP-EGFP cells were treated with 1 µM VHL-SNAP2-5C for 24 h with or without additional pre-treatment with 1 µM SNAP2-ligand for 30 min. SNAP-EGFP levels were assessed by flow cytometry. (n = 3, data represent mean +/- s.d.).

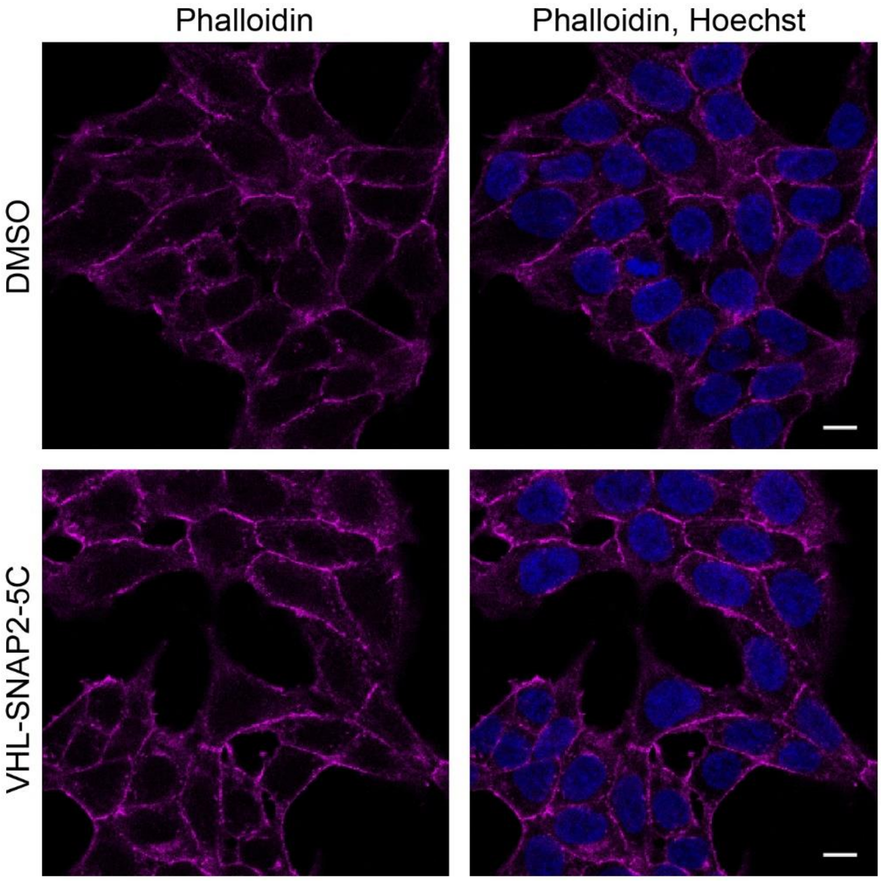


**Figure S5. Analysis of actin cytoskeleton upon SNAP-Clca$^{EN}$ depletion.** Confocal images of HAP1 SNAP-CLCa$^{EN}$ cells treated with 1 µM VHL-SNAP2-5C for 24 h and stained with Phalloidin to assess actin cytoskeleton organization. Images were taken with 63x magnification. Scale bars are 10 µm.

**Table S1. ΔΔG values for ligands predicted by FEP using benzyl chloropyrimidine as a reference.**

| Ligand ID | Predicted ΔΔG (kcal/mol) |
|---|---|
| 1a | -1.3 ± 0.55 |
| 1b (meta 1) | -0.5 ± 0.55 |
| 1b (meta 2) | -0.9 ± 0.56 |
| 1c | -1.6 ± 0.62 |
| 1d (protonated) | -1.9 ± 0.62 |
| 1d (not protonated) | -1.5 ± 0.57 |
| 1e (orientation 1) | 0.4 ± 0.45 |
| 1e (orientation 2) | -0.3 ± 0.58 |
| 1f (orientation 1) | 0.2 ± 0.42 |
| 1f (orientation 2) | 1.2 ± 0.42 |
| 1g | -0.4 ± 0.56 |
| 1h (orientation 1) | -0.6 ± 0.80 |
| 1h (orientation 2) | -0.6 ± 0.80 |
| 1i | -1.7 ± 0.45 |
| 1j (orientation 1) | -1.4 ± 0.82 |
| 1j (orientation 2) | -2.9 ± 0.82 |
| 1l | -0.7 ± 0.43 |
| 1m (orientation 1) | -0.5 ± 0.55 |
| 1m (orientation 2) | -0.1 ± 0.55 |
| Reference benzyl chloropyrimidine | 0.0 ± 0.40 |

**Table S2**. Chemical stability data for selected compound

| Entry | Compound | pH = 1<br>t ½ (days) | pH = 7.4<br>t ½ (days) | pH = 10<br>t ½ (days) |
|---|---|---|---|---|
| 1 | **1a** | 15.7 | >149 | >149 |
| 2 | **1c** | 4.5 | >149 | >149 |
| 3 | **1i** | N.D | >149 | >149 |
| 4 | **1l** | N.D | 0.51 | 0.52 |

**Table S3**. Purity, physicochemical properties, VHL binding and Cytotoxic of selected PROTACs

| Entry | Compound | UV purity (%) | MW | Solubility pH=7.4 (µM) | ChromLogD pH=7.4 | ePSA (Å) | VHL binding $IC_{50}$ (µM) | Cytotoxicity $IC_{50}$ (µM) |
|---|---|---|---|---|---|---|---|---|
| 1 | **VHL-SNAP1-4C** | >99 | 825.4 | 64 | 1.6 | 122 | 0.40 | >50 |
| 2 | **VHL-SNAP2-4C** | 99 | 819.3 | 18 | 3.0 | 110 | 0.25 | >49 |
| 3 | **VHL-SNAP1-5C** | 90 | 839.4 | 13 | 1.7 | 124 | 0.24 | >49 |
| 4 | **VHL-SNAP2-5C** | 97 | 833.4 | 17 | 3.2 | 111 | 0.33 | >50 |
| 5 | **VHL-SNAP1-6C** | 96 | 853.4 | <31 | 1.9 | 123 | 0.67 | >40 |
| 6 | **VHL-SNAP2-6C** | 99 | 847.4 | <9 | 3.4 | 111 | 0.39 | >50 |
| 7 | **N-VHL-SNAP2-5C** | 96 | 819.4 | <13 | 2.9 | 112 | >100 | N.D |
| 8 | **VHL-CLIP-4C** | 94 | 785.0 | 24.5 | 2.1 | 112 | 0.16 | >50 |
| 9 | **VHL-CLIP-5C** | 98 | 799.0 | 28.0 | 2.3 | 114 | 0.22 | >50 |
| 10 | **VHL-CLIP-6C** | 96 | 813.0 | 6.5 | 2.5 | 112 | 0.24 | 34 |

N.D = Not determined. In house protocol was used for determination of physiochemical properties. VHL binding was determined using fluorescence polarization (FP) assay using purified VHL and a FAM-labelled HIF1α derived probe. Cytotoxicity was determined in a human monocytic cell line (THP1) using Resazurin reagent.

**Table S4**. Purity, physicochemical properties, CRBN binding and Cytotoxic of selected PROTACs

| Entry | Compound | UV purity (%) | MW | Solubility pH=7.4 (µM) | ChromLogD pH=7.4 | ePSA (Å) | CRBN binding $IC_{50}$ (µM) | Cytotoxicity $IC_{50}$ (µM) |
|---|---|---|---|---|---|---|---|---|
| 1 | **CRBN5-SNAP2-0C-PIP** | 91 | 632.1 | 1.4 | 2.8 | 126.9 | 0.007 | >50 |
| 2 | **CRBN5-SNAP2-1C-PIP** | 97 | 646.1 | 3.8 | 2.9 | 124.8 | 0.010 | >50 |
| 3 | **N-CRBN5-SNAP2-1C-PIP** | 93 | 632.1 | <1 | 3.3 | 120.9 | 10.2 | N.D |
| 4 | **N-CRBN5-SNAP2-OC-PIP** | 91 | 618.1 | <0.7 | 3.2 | 123.6 | 6.9 | N.D |

N.D = Not determined. In house protocol was used for determination of physicochemical properties. CRBN binding was determined in a time-resolved-FRET competition assay using a Cy5-labeled CRBN ligand. Cytotoxicity was determined in a human monocytic cell line (THP1) using Resazurin reagent.

### Synthesis procedures

#### General reagent and purification information

All commercial reagents were obtained from Sigma-Aldrich, Combi-Blocks, Enamine, Fluorochem, Abosyn or AstaTech. Solvents were used as obtained from suppliers. Normal phase flash column chromatography (NPFCC) and reversed phase flash chromatography (RPFCC) was performed using prepacked Silica HC D 20 µm columns and C18 D 100 Å 30 µm columns, respectively, on a Biotage Selekt system. For RPFCC acidic (A: $H_2O$/formic acid 100/0.1, B = MeCN) mobile phase was used. Purification by preparative reverse phase HPLC was performed on a Kromasil C8 column (10 µm, 21.2x250 ID mm) using an acidic mobile phase (A: $H_2O$/MeCN/formic acid 95/5/0.2, B: MeCN). Alternatively, preparative reverse phase HPLC was performed on a XBridge C18 column (10 µm 18x250 ID mm) using a basic mobile phase (A: $H_2O$/MeCN/$NH_3$ 95/5/0.2, B: MeCN). Purification by SFC was performed on a Waters Prep 100 SFC system with a 3010 mass detector and a 2998 photodiode arrau detector on a Waters BEH 2-EP column (5 µM, 30x250 mm)

#### General analytical information

All new compounds were characterized by NMR spectroscopy and mass spectrometry (MS). NMR spectra were recorded on a Bruker Ultrashield 500 MHz spectrometer with a Bruker Cryo Platform. NMR data are reported as follows: chemical shift in reference to the residual solvent peak (δ ppm), multiplicity (s = singlet, br s = broad singlet, d = doublet, t = triplet, q = quartet, dd = doublet of doublets, td = triplet of doublets, m = multiplet), coupling constant (Hz), and integration. $^{1}$H NMR residual solvent peaks in respective deuterated solvents for chloroform-*d* at 7.26 ppm, DMSO-$d^6$ at 2.50 ppm, methanol-$d^4$at 3.31 ppm and acetone- $d^6$. $^{13}$C NMR residual solvent peaks in respective deuterated solvents for chloroform-*d* at 77.16 ppm, DMSO-$d^6$ at 39.52 ppm, methanol-$d^4$ at 49.00 ppm. HPLC-MS analysis was performed on a Waters Acquity UPLC system using an acidic mobile phase with a HSS C18 column (1.8 µm, 50 × 2.1 mm) or using a basic mobile phase with a BEH C18 column (1.7 µm, 50 × 2.1 mm).

#### VHL recruiting PROTACs

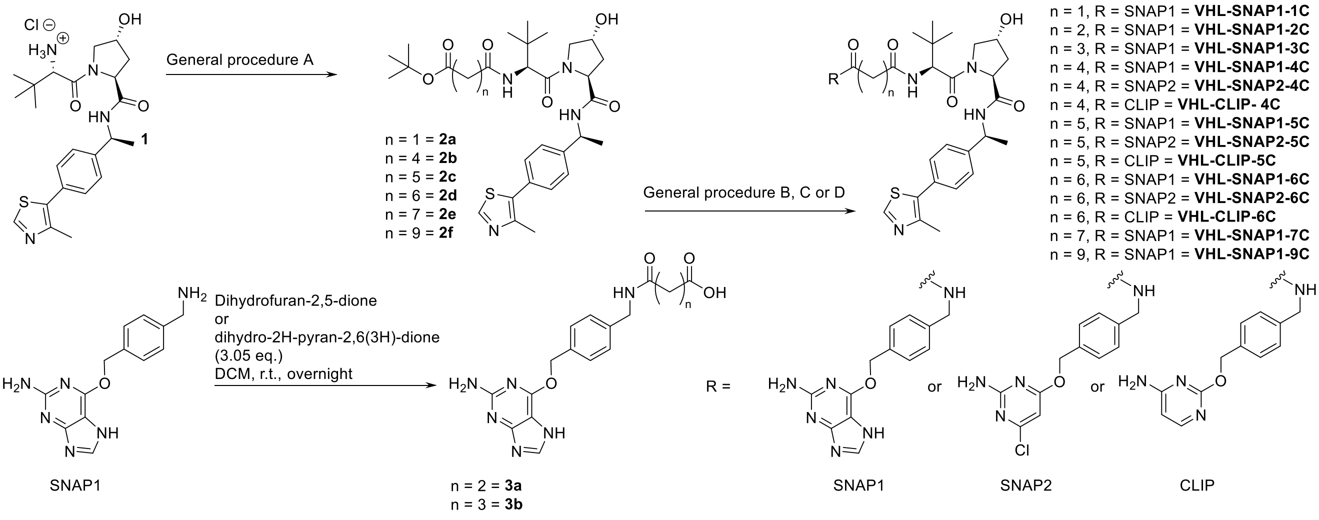


**Scheme S1**: Overview of synthesis of VHL-SNAP1, SNAP2 or CLIP-PROTACs

**General procedure A**

(2*S*,4*R*)-1-((*S*)-2-Amino-3,3-dimethylbutanoyl)-4-hydroxy-*N*-((*S*)-1-(4-(4-methylthiazol-5-yl)phenyl)ethyl)pyrrolidine-2-carboxamide hydrochloride **(1)** (1.1 eq.), the corresponding mono *tert*-butoxy diacid (1 eq.), PyOxim (1.5 eq.) and OXYMA Pure (0.5 eq.) were taken up in DMF (0.1 M) and DIPEA (5 eq.) added. The reaction was stirred at room temperature until completion monitored by LCMS. The reaction mixture was diluted with EtOAc and washed once with saturated $NaHCO_3$ and twice with brine. The organic layer was dried over $MgSO_4$, filtered, concentrated, and purified by NPFCC to afford products **2a**,**2d**-**h**.

***Tert*-butyl-3-(((*S*)-1-((2*S*,4*R*)-4-hydroxy-2-(((*S*)-1-(4-(4-methylthiazol-5-yl)phenyl)ethyl)carbamoyl)pyrrolidin-1-yl)-3,3-dimethyl-1-oxobutan-2-yl)amino)-3-oxopropanoate (2a)**

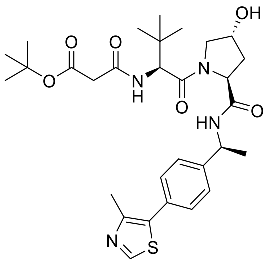

**1** (60 mg, 0.12 mmol) and 3-(*tert*-butoxy)-3-oxopropanoic acid (1.0 eq.) were used to synthesise **2a** according to general procedure A. The reaction was stirred for 3 h. The crude product was purified by NPFCC (2-15% methanol in EtOAc over 7 min). **2a** was obtained as a pale-yellow oil (24 mg, 33%). **$^1$H NMR** (500 MHz, Methanol-*d$_4$*) δ: 1.09 (s, 9H), 1.50 (s, 9H), 1.53 (d, *J* = 7.0 Hz, 3H), 1.98 (ddd, *J* = 13.3, 9.0, 4.6 Hz, 1H), 2.22 (ddt, *J* = 13.1, 7.7, 2.0 Hz, 1H), 2.50 (s, 3H), 3.25 (d, *J* = 15.8 Hz, 1H), 3.39 (d, 1H), 3.78 (dd, *J* = 11.0, 3.9 Hz, 1H), 3.89 (dt, *J* = 11.2, 1.8 Hz, 1H), 7.42–7.49 (m, 4H), 8.89 (s, 1H). **MS** (m/z): $[M + H]^+$ calcd. for $C_{30}H_{43}N_4O_6S$, 587.3; found 587.5.

***Tert*-butyl-6-(((*S*)-1-((2*S*,4*R*)-4-hydroxy-2-(((*S*)-1-(4-(4-methylthiazol-5-yl)phenyl)ethyl)carbamoyl)pyrrolidin-1-yl)-3,3-dimethyl-1-oxobutan-2-yl)amino)-6-oxohexanoat (2b)**

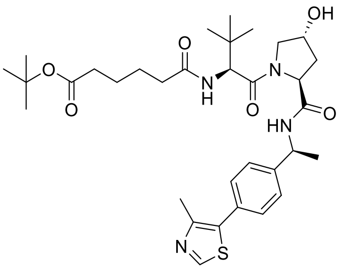

**1** (90 mg, 0.19 mmol) and 6-(*tert*-butoxy)-6-oxohexanoic acid (1.0 eq.) were used to synthesise **2b** according to general procedure A. The reaction was stirred overnight. The crude product was purified by NPFCC (0-20% methanol in EtOAc over 7 min). **2b** was obtained as a pale-yellow oil (96 mg, 90%). **$^1$H NMR** (500 MHz, Chloroform-*d*) δ: 1.04 (s, 9H), 1.43 (s, 9H), 1.46 (d, *J* = 6.9 Hz, 3H), 1.53–1.66 (m, 4H), 2.05 (ddt, *J* = 14.4, 8.2, 2.1 Hz, 1H), 2.13–2.25 (m, 4H), 2.45–2.54 (m, 4H), 3.50 (s, 1H), 3.60 (dd, *J* = 11.3, 3.8 Hz, 1H), 4.07 (dt, *J* = 11.4, 1.9 Hz, 1H), 4.48–4.51 (m, 1H), 4.55 (d, *J* = 8.6 Hz, 1H), 4.71 (t, *J* = 7.9 Hz, 1H), 5.08 (p, *J* = 7.1 Hz, 1H), 6.34 (d, *J* = 8.6 Hz, 1H), 7.38 (dd, 4H), 7.46 (d, *J* = 7.9 Hz, 1H), 8.67 (s, 1H). **MS** (m/z): $[M + H]^+$ calcd. for $C_{33}H_{49}N_4O_6S$, 629.3; found 629.4.

***Tert*-butyl-7-(((*S*)-1-((2*S*,4*R*)-4-hydroxy-2-(((*S*)-1-(4-(4-methylthiazol-5-yl)phenyl)ethyl)carbamoyl)pyrrolidin-1-yl)-3,3-dimethyl-1-oxobutan-2-yl)amino)-7-oxoheptanoate (2c)**

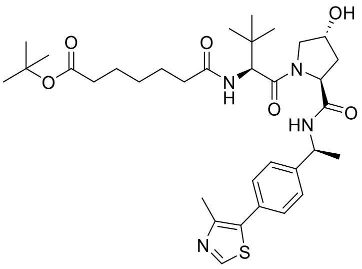

**1** (0.10 g, 0.21 mmol) and 7-(*tert*-butoxy)-7-oxoheptanoic acid (1.0 eq.) were used to synthesise **2c** according to general procedure A. The reaction was stirred overnight. The crude product was purified by NPFCC (0-20% methanol in EtOAc over 7 min). **2c** was obtained as a pale-yellow oil (94 mg, 77%). **$^1$H NMR** (500 MHz, Chloroform-*d*) δ: 1.04 (s, 9H), 1.28–1.36 (m, 2H), 1.43 (s, 9H), 1.47 (d, *J* = 6.9 Hz, 3H), 1.54–1.63 (m, 4H), 2.06 (ddt, *J* = 13.0, 8.2, 2.0 Hz, 1H), 2.15–2.25 (m, 4H), 2.53 (s,

4H), 3.12 (s, 1H), 3.60 (dd, *J* = 11.4, 3.7 Hz, 1H), 4.10 (dt, *J* = 11.5, 1.9 Hz, 1H), 4.52 (d, *J* = 8.5 Hz, 2H), 4.72 (t, *J* = 7.9 Hz, 1H), 5.08 (p, *J* = 7.1 Hz, 1H), 6.12 (d, *J* = 8.5 Hz, 1H), 7.35–7.42 (m, 4H), 7.46 (d, *J* = 7.9 Hz, 1H), 8.67 (s, 1H). **MS** (m/z): $[M + H]^+$ calcd. for $C_{34}H_{51}N_4O_6S$, 643.4; found 643.4.

#### ***Tert*-butyl-8-(((*S*)-1-((2*S*,4*R*)-4-hydroxy-2-(((*S*)-1-(4-(4-methylthiazol-5-yl)phenyl)ethyl)carbamoyl)pyrrolidin-1-yl)-3,3-dimethyl-1-oxobutan-2-yl)amino)-8-oxooctanoate (2d)**

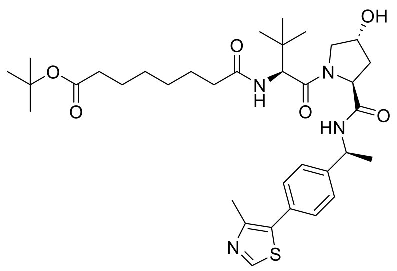

**1** (0.10 g, 0.21 mmol) and 8-(*tert*-butoxy)-8-oxooctanoic acid (1.0 eq.) were used to synthesise **2d** according to general procedure A. The reaction was stirred overnight. The crude product was purified by NPFCC (0-20% methanol in EtOAc over 7 min). **2d** was obtained as a pale-yellow oil (96 mg, 78%). **$^1$H NMR** (500 MHz, Chloroform-*d*) δ: 1.04 (s, 9H), 1.23–1.34 (m, 5H), 1.43 (s, 9H), 1.47 (d, *J* = 6.9 Hz, 3H), 1.53–1.63 (m, 4H), 2.06 (ddt, *J* = 13.0, 8.3, 2.1 Hz, 1H), 2.13–2.22 (m, 4H), 2.52 (s, 4H), 3.60 (dd, *J* = 11.3, 3.8 Hz, 1H), 4.10 (dt, *J* = 11.5, 1.9 Hz, 1H), 4.48–4.56 (m, 2H), 4.72 (t, *J* = 7.9 Hz, 1H), 5.08 (p, *J* = 7.0 Hz, 1H), 6.14 (d, *J* = 8.6 Hz, 1H), 7.34–7.43 (m, 4H), 7.47 (d, *J* = 7.9 Hz, 1H), 8.68 (s, 1H). **MS** (m/z): $[M + H]^+$ calcd. for $C_{35}H_{53}N_4O_6S$, 657.4; found 657.5.

#### ***Tert*-butyl-9-(((*S*)-1-((2*S*,4*R*)-4-hydroxy-2-(((*S*)-1-(4-(4-methylthiazol-5-yl)phenyl)ethyl)carbamoyl)pyrrolidin-1-yl)-3,3-dimethyl-1-oxobutan-2-yl)amino)-9-oxononanoate (2e)**

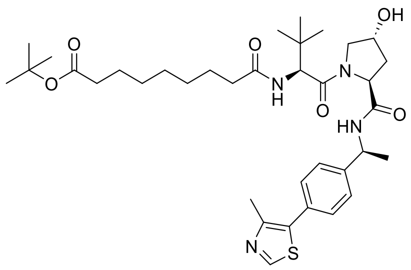

**1** (60 mg, 0.12 mmol) and 9-(*tert*-butoxy)-9-oxononanoic acid (1.0 eq.) were used to synthesise **2e** according to general procedure A. The reaction was stirred for 3 h. The crude was purified by NPFCC (2-15% methanol in EtOAc over 7 min). **2e** was obtained as a pale-yellow oil (57 mg, 68%). **$^1$H NMR** (500 MHz, Methanol-*d$_4$*) δ: 1.07 (s, 9H), 1.36 (d, *J* = 3.3 Hz, 6H), 1.47 (s, 9H), 1.53 (d, *J* = 7.0 Hz, 3H), 1.57–1.69 (m, 4H), 1.99 (ddd, *J* = 13.3, 9.0, 4.6 Hz, 1H), 2.17–2.38 (m, 5H), 2.50 (s, 3H), 3.78 (dd, *J* = 11.0, 4.0 Hz, 1H), 3.91 (dt, *J* = 11.2, 1.8 Hz, 1H), 4.45–4.47 (m, 1H), 4.61 (t, *J* = 8.3 Hz, 1H), 4.66 (d, *J* = 9.0 Hz, 1H), 4.99–5.09 (m, 1H), 7.42–7.50 (m, 4H), 7.83 (d, *J* = 9.0 Hz, 1H), 8.56 (d, *J* = 7.5 Hz, 1H), 8.90 (s, 1H). **MS** (m/z): $[M + H]^+$ calcd. for $C_{36}H_{55}N_4O_6S$, 671.4; found 671.9.

#### ***Tert*-butyl-11-(((*S*)-1-((2*S*,4*R*)-4-hydroxy-2-(((*S*)-1-(4-(4-methylthiazol-5-yl)phenyl)ethyl)carbamoyl)pyrrolidin-1-yl)-3,3-dimethyl-1-oxobutan-2-yl)amino)-11-oxoundecanoate (2f)**

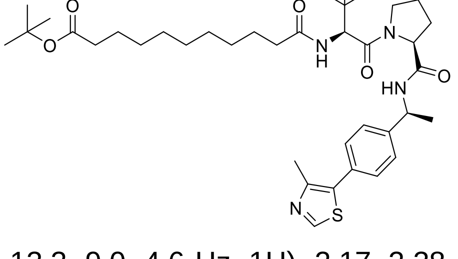

**1** (60 mg, 0.12 mmol) and 11-(*tert*-butoxy)-11-oxoundecanoic acid (1.0 eq.) were used to synthesise **2f** according to general procedure A. The reaction was stirred for 3 h. The crude product was purified by NPFCC (2-15% methanol in EtOAc over 7 min). **2f** was obtained as a pale-yellow oil (45 mg, 57%). **$^1$H NMR** (500 MHz, Methanol-*d$_4$*) δ: 1.07 (s, 9H), 1.17 (d, *J* = 6.2 Hz, 1H), 1.30–1.38 (m, 10H), 1.46 (s, 9H), 1.53 (d, *J* = 7.0 Hz, 3H), 1.55–1.71 (m, 4H), 1.98 (ddd, *J* = 13.3, 9.0, 4.6 Hz, 1H), 2.17–2.38 (m, 5H), 2.50 (s, 3H), 3.77 (dd, *J* = 11.0, 4.0 Hz, 1H), 3.87–3.95 (m, 1H), 4.45–4.46 (m, 1H), 4.60 (t, *J* = 8.2 Hz, 1H), 4.66 (d, *J* = 9.0 Hz, 1H), 5.03 (p, *J* = 7.0 Hz, 1H), 7.41–7.49 (m, 4H), 7.82

(d, $J$ = 8.9 Hz, 1H), 8.56 (d, $J$ = 7.5 Hz, 1H), 8.89 (s, 1H). **MS** (m/z): $[M + H]^+$ calcd. for $C_{38}H_{59}N_4O_6S$, 699.4; found 699.6.

**4-((4-(((2-Amino-7*H*-purin-6-yl)oxy)methyl)benzyl)amino)-4-oxobutanoic acid (3a)**

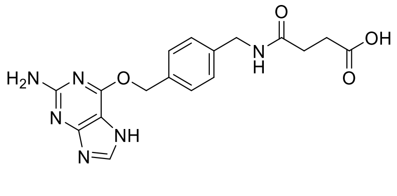


6-((4-(Aminomethyl)benzyl)oxy)-7*H*-purin-2-amine (SNAP1 ligand) (50 mg, 0.18 mmol) and dihydrofuran-2,5-dione (56 mg, 0.55 mmol, 3.05 eq.) were dissolved in DCM (3 mL) and stirred overnight at room temperature. The obtained solid was filtered off and washed with DCM. The crude product was purified by preparative HPLC (acidic, 0-30% B in A over 20 min) to afford **3a** as a fluffy white solid (29 mg, 42%). **$^1$H NMR** (500 MHz, DMSO-$d_6$) δ: 2.35 (t, $J$ = 6.8 Hz, 2H), 2.43 (t, $J$ = 6.8 Hz, 2H), 4.24 (d, $J$ = 5.8 Hz, 2H), 5.43 (s, 2H), 5.73 (s, 2H), 6.25 (s, 1H), 7.25 (d, $J$ = 7.9 Hz, 2H), 7.42 (d, $J$ = 7.8 Hz, 2H), 7.78 (s, 1H), 8.35 (t, $J$ = 6.0 Hz, 1H). **MS** (m/z): $[M + H]^+$ calcd. for $C_{17}H_{19}N_6O_4$, 371.2; found 371.3.

**5-((4-(((2-Amino-7*H*-purin-6-yl)oxy)methyl)benzyl)amino)-5-oxopentanoic acid (3b)**

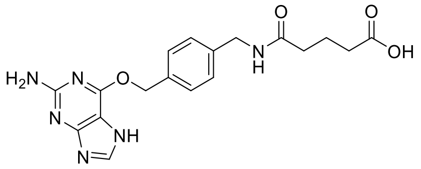


6-((4-(Aminomethyl)benzyl)oxy)-7*H*-purin-2-amine (SNAP1 ligand) (50 mg, 0.18 mmol) and dihydro-2*H*-pyran-2,6(3*H*)-dione (63 mg, 0.55 mmol, 3.05 eq.) were dissolved in DCM (3 mL) and stirred overnight at room temperature. The obtained solid was filtered off and washed with DCM. The crude product was purified by preparative HPLC (acidic, 0-30% B in A over 20 min) to afford **3b** as a fluffy white solid (29 mg, 29%).**$^1$H NMR** (500 MHz, DMSO-$d_6$) δ: 1.72 (p, $J$ = 7.4 Hz, 2H), 2.15 (t, $J$ = 7.4 Hz, 2H), 2.20 (t, $J$ = 7.4 Hz, 2H), 4.24 (d, $J$ = 5.9 Hz, 2H), 5.44 (s, 2H), 6.26 (s, 1H), 7.25 (d, $J$ = 7.8 Hz, 2H), 7.44 (d, $J$ = 7.8 Hz, 2H), 7.79 (s, 1H), 8.31 (t, $J$ = 5.9 Hz, 1H). **MS** (m/z): $[M + H]^+$ calcd. for $C_{18}H_{21}N_6O_4$, 385.2; found 385.3

**General procedure B**

The corresponding ester was dissolved in DCM (40 mL) and HCl (dioxane, 4 M, 30 eq.) was added. The reaction was stirred at room temperature until completion monitored by LCMS. The solvent was removed under reduced pressure and the crude was used in the next step without purification.

The crude acid (1.1 eq.), the amine (6-((4-(aminomethyl)benzyl)oxy)-7H-purin-2-amine (SNAP1 ligand), 4-((4-(aminomethyl)benzyl)oxy)-6-chloropyrimidin-2-amine (SNAP2 ligand) or 2-((4-(aminomethyl)benzyl)oxy)pyrimidin-4-amine (CLIP ligand) (1 eq.) and HATU (1.2 eq.) were taken up in DMF (0.1 M) and DIPEA (10 eq.) added. The reaction mixture was stirred at room temperature until completion monitored by LCMS. The mixture was diluted in EtOAc, washed once with saturated $NaHCO_3$ and extracted twice with EtOAc. The combined organic layer was dried over $MgSO_4$, filtered, concentrated, and purified by preparative HPLC to afford the desired PROTACs.

**General procedure C**

The corresponding ester was dissolved in DCM (40 mL) and HCl (dioxane, 4 M 30 eq.) was added. The reaction was stirred at room temperature until completion monitored by LCMS. The solvent was removed under reduced pressure and the crude was used in the next step without purification.

The crude acid (1.1 eq.), the amine (SNAP1, SNAP2 or CLIP ligand) (1 eq.), PyOxim (1.5 eq.) and OXYMA Pure (0.5 eq.) were taken up in DMF (0.1 M) and DIPEA (10 eq.) added. The reaction mixture was stirred at room temperature until completion monitored by LCMS. The mixture was diluted in EtOAc, washed once with saturated $NaHCO_3$ and extracted twice with EtOAc. The combined organic layer was dried over $MgSO_4$, filtered, concentrated, and purified by preparative HPLC to afford the desired PROTACs.

**General procedure D**

The corresponding carboxylic acid (1.1 eq.), **1** (1 eq.) and HATU (1.2 eq.) were taken up in DMF (0.1 M) and DIPEA (10 eq.) was added. The reaction mixture was stirred at room temperature until completion monitored by LCMS. The mixture was diluted in EtOAc, washed once with saturated $NaHCO_3$ and extracted twice with EtOAc. The combined organic layer was dried over $MgSO_4$, filtered, concentrated, and purified by preparative HPLC to afford the desired PROTACs.

***N*$^1$-(4-(((2-Amino-7*H*-purin-6-yl)oxy)methyl)benzyl)-*N*$^3$-((*S*)-1-((2*S*,4*R*)-4-hydroxy-2-(((*S*)-1-(4-(4-methylthiazol-5-yl)phenyl)ethyl)carbamoyl)pyrrolidin-1-yl)-3,3-dimethyl-1-oxobutan-2-yl)malonamide (VHL-SNAP1-1C)**

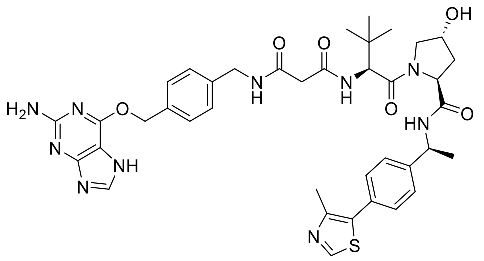

**2a** (22 mg, 0.04 mmol) and SNAP1 ligand were used to synthesise **VHL-SNAP1-1C** according to general procedure B with the exception that 3.0 eq. of HATU and 3.0 eq. of SNAP1 ligand were used. The reaction was stirred for 3 h. The crude product was purified by preparative HPLC (basic, 5-45% B in A over 20 min). **VHL-SNAP1-1C** was obtained as a fluffy white solid (3.8 mg, 13%). **$^1$H NMR** (500 MHz, Methanol-*d$_4$*) δ: 1.01 (s, 9H), 1.48 (d, *J* = 7.0 Hz, 3H), 1.88–1.97 (m, 2H), 2.17 (ddt, *J* = 13.1, 7.7, 2.1 Hz, 1H), 2.45 (s, 3H), 3.72 (dd, *J* = 11.0, 3.9 Hz, 1H), 3.86 (dd, *J* = 11.2, 1.8 Hz, 1H), 4.37–4.43 (m, 3H), 4.55 (dd, *J* = 9.1, 7.8 Hz, 1H), 4.59 (s, 1H), 4.99 (q, *J* = 7.0 Hz, 1H), 5.51 (s, 2H), 7.30 (d, 2H), 7.38–7.43 (m, 4H), 7.46 (d, 2H), 7.81 (s, 1H), 8.85 (s, 1H). **MS** (m/z): $[M + H]^+$ calcd. for $C_{39}H_{47}N_{10}O_6S$, 783.3; found 783.6.

***N*$^1$-(4-(((2-Amino-7*H*-purin-6-yl)oxy)methyl)benzyl)-*N*$^4$-((*S*)-1-((2*S*,4*R*)-4-hydroxy-2-(((*S*)-1-(4-(4-methylthiazol-5-yl)phenyl)ethyl)carbamoyl)pyrrolidin-1-yl)-3,3-dimethyl-1-oxobutan-2-yl)succinamide (VHL-SNAP1-2C)**

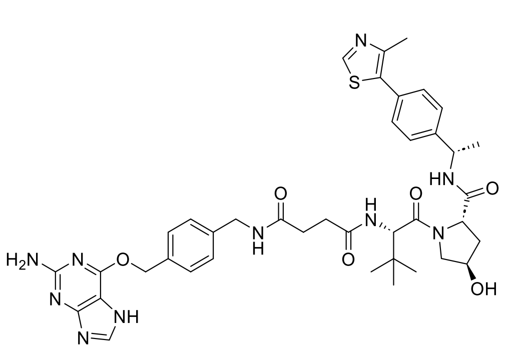

**3a** (5.8 mg, 0.02 mmol) and **1** were used to synthesise **VHL-SNAP1-2C** according to general procedure D. The reaction was stirred for 3 h. The crude product was purified by preparative HPLC (acidic, 15-55% B in A over 20 min). **VHL-SNAP1-2C** was obtained as a fluffy white solid (2.6 mg, 23%). **$^1$H NMR** (500 MHz, Methanol-*d$_4$*) δ: 1.04 (s, 9H), 1.52 (d, *J* = 7.0 Hz, 2H), 1.95 (ddd, *J* = 13.3, 9.0, 4.6 Hz, 1H), 2.20 (dd, *J* = 13.0, 8.0 Hz, 1H), 2.49 (s, 3H), 2.52–2.70 (m, 4H), 3.73 (dd, *J* = 11.1, 3.9 Hz, 1H), 3.87 (d, *J* = 11.1 Hz, 1H), 4.29–4.45 (3H, m), 4.54–4.63 (m, 2H), 5.02 (p, 1H), 5.55 (s, 2H), 7.32 (d, *J* = 7.6 Hz, 2H), 7.41–7.48 (m, 4H), 7.50 (d, *J* = 7.7 Hz, 2H), 7.86 (s, 1H), 8.89 (s, 1H). **MS** (m/z): $[M + H]^+$ calcd. for $C_{40}H_{49}N_{10}O_6S$, 797.4; found 797.6.

**$N^1$-(4-(((2-Amino-7*H*-purin-6-yl)oxy)methyl)benzyl)-$N^5$-((*S*)-1-((2*S*,4*R*)-4-hydroxy-2-(((*S*)-1-(4-(4-methylthiazol-5-yl)phenyl)ethyl)carbamoyl)pyrrolidin-1-yl)-3,3-dimethyl-1-oxobutan-2-yl)glutaramide (VHL-SNAP1-3C)**

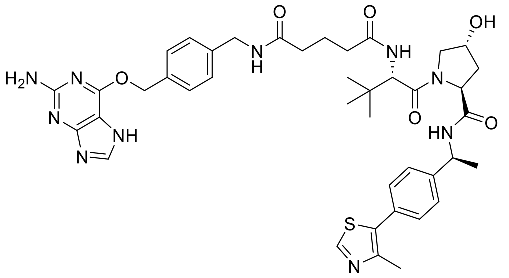

**3b** (9.4 mg, 0.02 mmol) and **1** were used to synthesise **VHL-SNAP1-3C** according to general procedure D. The reaction was stirred for 3 h. The crude product was purified by preparative HPLC (acidic, 15-55% B in A over 20 min). **VHL-SNAP1-3C** was obtained as a fluffy white solid (4.2 mg, 23%). **$^1$H NMR** (500 MHz, Methanol-*d$_4$*) δ: 1.05 (s, 9H), 1.50 (d, *J* = 7.0 Hz, 3H), 1.89–2.01 (m, 4H), 2.15–2.23 (m, 1H), 2.24–2.38 (m, 5H), 2.49 (s, 3H), 3.76 (dd, *J* = 11.0, 4.0 Hz, 1H), 3.90 (d, *J* = 11.0 Hz, 1H), 4.33–4.49 (m, 3H), 4.51–4.63 (m, 2H), 5.01 (p, *J* = 6.8 Hz, 1H), 5.56 (s, 2H), 7.32 (d, *J* = 7.9 Hz, 2H), 7.39 (dd, *J* = 8.2, 4.4 Hz, 1H), 7.41–7.47 (m, 4H), 7.51 (d, *J* = 7.9 Hz, 2H), 7.86 (s, 1H), 7.89 (d, *J* = 8.7 Hz, 1H), 8.56 (d, *J* = 7.6 Hz, 1H), 8.89 (s, 1H). **MS** (m/z): [M + H]$^+$ calcd. for $C_{41}H_{51}N_{10}O_6S$, 811.4; found 811.5.

**$N^1$-(4-(((2-Amino-7*H*-purin-6-yl)oxy)methyl)benzyl)-$N^6$-((*S*)-1-((2*S*,4*R*)-4-hydroxy-2-(((*S*)-1-(4-(4-methylthiazol-5-yl)phenyl)ethyl)carbamoyl)pyrrolidin-1-yl)-3,3-dimethyl-1-oxobutan-2-yl)adipamide (VHL-SNAP1-4C)**

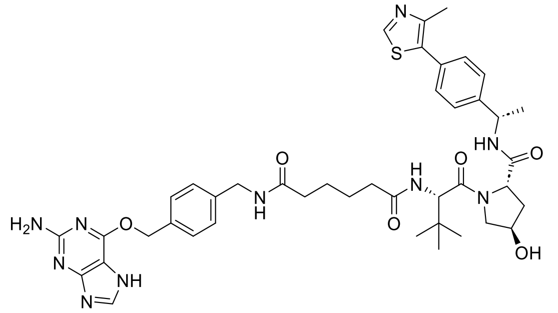

**2b** (23 mg, 0.04 mmol) and SNAP1 ligand were used to synthesise **VHL-SNAP1-4C** according to general procedure C. The reaction was stirred for 4 h. The crude product was purified by preparative HPLC (basic, 15-55% B in A over 20 min). **VHL-SNAP1-4C** was obtained as a fluffy white solid (18 mg, 53%) **$^1$H NMR** (500 MHz, Methanol-*d$_4$*) δ: 1.04 (s, 9H), 1.50 (d, *J* = 7.0 Hz, 3H), 1.62–1.68 (m, 3H), 1.92–2.01 (m, 2H), 2.19 (ddt, *J* = 13.2, 7.8, 2.0 Hz, 1H), 2.23–2.38 (m, 4H), 2.48 (s, 3H), 2.67 (s, 1H), 3.75 (dd, *J* = 11.0, 4.0 Hz, 1H), 3.89 (dt, *J* = 11.3, 1.7 Hz, 1H), 4.38 (s, 2H), 4.40–4.47 (m, 1H), 4.58 (t, *J* = 8.3 Hz, 1H), 4.62 (s, 1H), 5.01 (q, *J* = 6.9 Hz, 1H), 5.54 (s, 2H), 7.31 (d, *J* = 8.0 Hz, 2H), 7.40–7.46 (m, 4H), 7.49 (d, *J* = 8.1 Hz, 2H), 7.84 (s, 1H), 8.88 (s, 1H). **$^{13}$C NMR** (126 MHz, Methanol-*d$_4$*) δ: 15.8, 22.4, 26.5, 26.6, 27.1, 36.2, 36.4, 36.7, 38.8, 40.4, 43.8, 50.1, 58.0, 59.1, 60.6, 68.6, 71.0, 127.6, 128.7, 129.7, 130.5, 131.5, 133.3, 136.9, 140.1, 145.7, 149.1, 152.8, 161.7, 172.3, 173.2, 175.7, 175.7. **MS** (m/z): [M + H]$^+$ calcd. for $C_{42}H_{53}N_{10}O_6S$, 825.387; found 825.392.

**$N^1$-(4-(((2-Amino-7*H*-purin-6-yl)oxy)methyl)benzyl)-$N^7$-((*S*)-1-((2*S*,4*R*)-4-hydroxy-2-(((*S*)-1-(4-(4-methylthiazol-5-yl)phenyl)ethyl)carbamoyl)pyrrolidin-1-yl)-3,3-dimethyl-1-oxobutan-2-yl)heptanediamide (VHL-SNAP1-5C)**

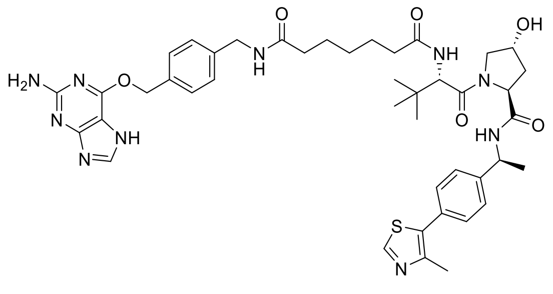

**2c** (29 mg, 0.05 mmol) and SNAP1 ligand were used to synthesise **VHL-SNAP1-5C** according to general procedure C. The reaction was stirred for 3 h. The crude product was purified by preparative HPLC (basic, 15-55% B in A over 20 min). **VHL-SNAP1-5C** was obtained as a fluffy white solid (6.6 mg, 16%). **$^1$H NMR** (500 MHz, Methanol-*d$_4$*) δ: 1.05 (s, 9H), 1.37 (p, *J* = 7.7

Hz, 2H), 1.52 (d, *J* = 7.0 Hz, 3H), 1.61–1.71 (m, 4H), 1.94–2.01 (m, 1H), 2.18–2.34 (m, 5H), 2.49 (s, 3H), 3.76 (dd, *J* = 11.0, 4.0 Hz, 1H), 3.89 (d, *J* = 11.1 Hz, 1H), 4.38 (s, 2H), 4.43–4.46 (m, 1H), 4.58 (t, *J* = 8.3 Hz, 1H), 4.63 (s, 1H), 5.01 (q, *J* = 7.1 Hz, 1H), 5.56 (s, 2H), 7.32 (d, *J* = 7.9 Hz, 2H), 7.41–7.48 (m, 4H), 7.51 (d, *J* = 8.1 Hz, 2H), 7.85 (s, 1H), 8.89 (s, 1H). $^{13}$**C NMR** (126 MHz, Methanol-*d₄*) δ: 13.2, 15.8, 19.3, 22.4, 26.6, 26.7, 27.1, 29.8, 36.4, 36.4, 36.9, 38.8, 43.8, 43.8, 50.1, 55.9, 58.0, 59.0, 60.6, 68.7, 71.0, 127.6, 128.7, 129.7 130.5, 131.5, 133.4, 136.9, 140.1, 145.7, 149.1, 152.8, 161.7, 172.3, 173.3, 175.9, 176.0. **MS** (m/z): $[M + H]^+$ calcd. for $C_{43}H_{55}N_{10}O_6S$, 839.403; found 839.401.

#### ***N*¹-(4-(((2-Amino-7*H*-purin-6-yl)oxy)methyl)benzyl)-*N*⁸-((*S*)-1-((2*S*,4*R*)-4-hydroxy-2-(((*S*)-1-(4-(4-methylthiazol-5-yl)phenyl)ethyl)carbamoyl)pyrrolidin-1-yl)-3,3-dimethyl-1-oxobutan-2-yl)octanediamide (VHL-SNAP1-6C)**

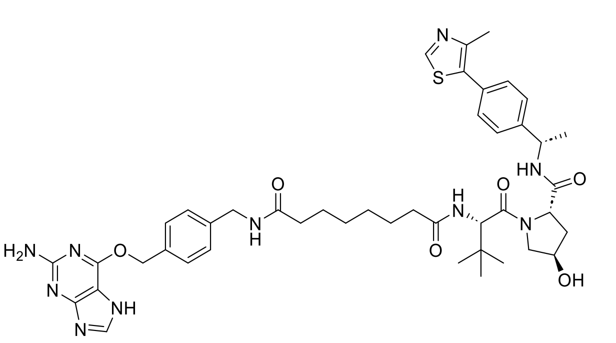

**2d** (33 mg, 0.05 mmol) and SNAP1 ligand were used to synthesise **VHL-SNAP1-6C** according to general procedure C. The reaction was stirred for 3 h. The crude product was purified by preparative HPLC (basic, 15-55% B in A over 20 min). **VHL-SNAP1-6C** was obtained as a fluffy white solid (15 mg, 31%). $^1$**H NMR** (500 MHz, Methanol-*d₄*) δ: 1.05 (s, 9H), 1.32–1.38 (m, 4H), 1.51 (d, *J* = 7.0 Hz, 3H), 1.56–1.71 (m, 4H), 1.92–2.01 (m, 1H), 2.15–2.34 (m, 5H), 2.49 (s, 3H), 2.67 (s, 1H), 3.75 (dd, *J* = 11.0, 4.0 Hz, 1H), 3.89 (dt, *J* = 11.2, 1.8 Hz, 1H), 4.38 (s, 2H), 4.41–4.47 (m, 1H), 4.59 (t, 1H), 4.63 (s, 1H), 5.01 (q, *J* = 7.0 Hz, 1H), 5.54 (s, 2H), 7.31 (d, *J* = 8.0 Hz, 2H), 7.40–7.46 (m, 4H), 7.50 (d, *J* = 8.0 Hz, 2H), 7.85 (s, 1H), 8.88 (s, 1H).$^{13}$**C NMR** (126 MHz, Methanol-*d₄*) δ: 15.79, 22.37, 26.85, 26.88, 27.05, 29.85, 29.90, 36.43, 36.53, 36.97, 38.76, 40.43, 43.81, 50.13, 58.00, 59.01, 60.58, 68.65, 70.96, 96.28, 106.42, 127.62, 128.69, 129.69, 129.76, 130.48, 131.50, 133.43, 136.94, 140.16, 145.65, 149.07, 152.83, 161.06, 172.31, 173.26, 175.95, 176.09. **MS** (m/z): $[M + H]^+$ calcd. for $C_{44}H_{57}N_{10}O_6S$, 853.418; found 853.420.

#### ***N*¹-(4-(((2-Amino-7*H*-purin-6-yl)oxy)methyl)benzyl)-*N*⁹-((*S*)-1-((2*S*,4*R*)-4-hydroxy-2-(((*S*)-1-(4-(4-methylthiazol-5-yl)phenyl)ethyl)carbamoyl)pyrrolidin-1-yl)-3,3-dimethyl-1-oxobutan-2-yl)nonanediamide (VHL-SNAP1-7C)**

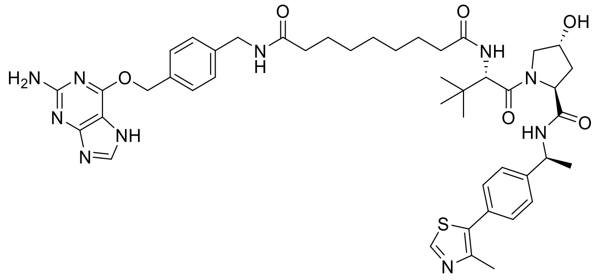

**2e** (26 mg, 0.04 mmol) and SNAP1 ligand were used to synthesise **VHL-SNAP1-7C** according to general procedure B. The reaction was stirred overnight. The crude product was purified by preparative HPLC (acidic, 15-55% B in A over 20 min). **VHL-SNAP1-7C** was obtained as a fluffy white solid (7.2 mg, 22%). $^1$**H NMR** (500 MHz, Methanol-*d₄*) δ: 1.01 (s, 9H), 1.24–1.35 (m, 6H), 1.47 (d, *J* = 7.0 Hz, 3H), 1.51–1.66 (m, 4H), 1.92 (ddd, *J* = 13.3, 9.0, 4.5 Hz, 1H), 2.12–2.31 (m, 5H), 2.44 (s, 3H), 3.72 (dd, *J* = 11.0, 4.0 Hz, 1H), 3.86 (dt, *J* = 11.1, 1.8 Hz, 1H), 4.34 (d, *J* = 4.7 Hz, 2H), 4.38–4.42 (m, 1H), 4.56 (t, *J* = 8.2 Hz, 1H), 4.60 (d, *J* = 9.0 Hz, 1H), 4.97 (q, 1H), 5.50 (s, 2H), 7.26 (d, *J* = 8.0 Hz, 2H), 7.36–7.42 (m, 4H), 7.45 (d, *J* = 8.0 Hz, 2H), 7.79–7.84 (m, 2H), 8.39 (t, *J* = 5.9 Hz, 1H), 8.58 (d, *J* = 7.5 Hz, 1H), 8.84 (s, 1H). **MS** (m/z): $[M + H]^+$ calcd. for $C_{45}H_{59}N_{10}O_6S$, 867.434; found 853.437.

#### $N^1$-(4-(((2-Amino-7*H*-purin-6-yl)oxy)methyl)benzyl)-$N^{11}$-((*S*)-1-((2*S*,4*R*)-4-hydroxy-2-(((*S*)-1-(4-(4-methylthiazol-5-yl)phenyl)ethyl)carbamoyl)pyrrolidin-1-yl)-3,3-dimethyl-1-oxobutan-2-yl)undecanediamide (VHL-SNAP1-9C)

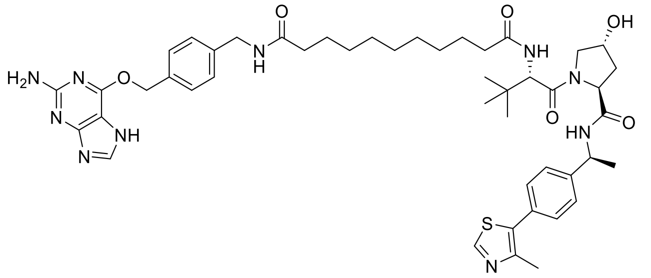

**2f** (22 mg, 0.03 mmol) and SNAP1 ligand were used to synthesise **VHL-SNAP1-9C** according to general procedure B. The reaction was stirred for 3 h. The crude product was purified by preparative HPLC (basic, 5-45% B in A over 20 min). **VHL-SNAP1-9C** was obtained as a fluffy white solid (9.6 mg, 31%). **$^1$H NMR** (500 MHz, Methanol-*d₄*) δ: 1.04 (s, 9H), 1.26–1.33 (m, 10H), 1.49 (d, *J* = 7.0 Hz, 3H), 1.54–1.65 (m, 4H), 1.90–1.99 (m, 1H), 2.16–2.33 (m, 5H), 2.47 (s, 3H), 3.74 (dd, *J* = 11.0, 4.0 Hz, 1H), 3.88 (dt, *J* = 11.3, 1.7 Hz, 1H), 4.36 (s, 2H), 4.41–4.45 (m, 1H), 4.55–4.61 (m, 1H), 4.62 (s, 1H), 5.00 (q, *J* = 7.0 Hz, 2H), 5.52 (s, 2H), 7.27–7.30 (m, 2H), 7.39–7.44 (m, 4H), 7.45–7.50 (m, 2H), 7.83 (s, 1H), 8.86 (s, 1H). **MS** (m/z): $[M + H]^+$ calcd. for $C_{47}H_{63}N_{10}O_6S$, 895.465; found 895.470.

#### $N^1$-(4-(((2-Amino-6-chloropyrimidin-4-yl)oxy)methyl)benzyl)-$N^6$-((*S*)-1-((2*S*,4*R*)-4-hydroxy-2-(((*S*)-1-(4-(4-methylthiazol-5-yl)phenyl)ethyl)carbamoyl)pyrrolidin-1-yl)-3,3-dimethyl-1-oxobutan-2-yl)adipamide (VHL-SNAP2-4C)

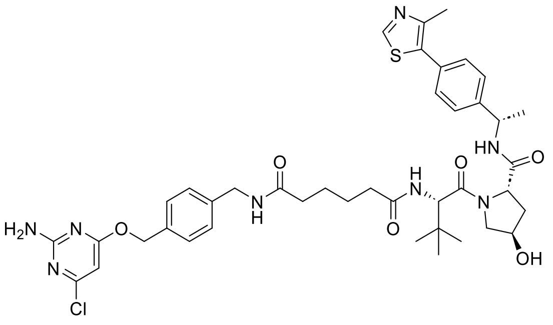

**2b** (22 mg, 0.04 mmol) and SNAP2 ligand were used to synthesise **VHL-SNAP2-4C** according to general procedure B. The reaction was stirred overnight. The crude product was purified by preparative HPLC (acidic, 15-55% B in A over 20 min). **VHL-SNAP2-4C** was obtained as a fluffy white solid (9.0 mg, 29%) **$^1$H NMR** (500 MHz, Methanol-*d₄*) δ: 1.04 (s, 9H), 1.49 (d, *J* = 7.0 Hz, 3H), 1.58–1.68 (m, 4H), 1.95 (ddd, *J* = 13.3, 9.0, 4.5 Hz, 1H), 2.14–2.35 (m, 5H), 2.48 (s, 3H), 3.74 (dd, *J* = 11.0, 3.9 Hz, 1H), 3.88 (d, *J* = 11.0 Hz, 1H), 4.36 (s, 2H), 4.42 (tt, *J* = 4.3, 2.0 Hz, 1H), 4.57 (t, *J* = 8.2 Hz, 1H), 4.60–4.64 (m, 1H), 5.00 (p, 1H), 5.33 (s, 2H), 6.08 (s, 1H), 7.28 (d, *J* = 7.9 Hz, 2H), 7.38 (d, *J* = 7.9 Hz, 2H), 7.40–7.48 (m, 4H), 7.87 (d, *J* = 8.9 Hz, 1H), 8.55 (d, *J* = 7.5 Hz, 1H), 8.93 (s, 1H). **$^{13}$C NMR** (126 MHz, Methanol-*d₄*) δ: 15.7, 22.4, 26.5, 26.6, 27.0, 36.3, 36.4, 36.7, 38.8, 43.8, 50.1, 50.2, 58.0, 59.1, 60.7, 69.0, 71.0, 96.6, 127.4, 127.6, 128.7, 129.6, 130.5, 131.3, 136.7, 140.1, 145.8, 148.8, 153.1, 161.7, 164.4, 172.3, 173.3, 175.7, 175.8. **MS** (m/z): $[M + H]^+$ calcd. for $C_{41}H_{52}ClN_8O_6S$, 819.342; found 819.343.

#### $N^1$-(4-(((2-Amino-6-chloropyrimidin-4-yl)oxy)methyl)benzyl)-$N^7$-((*S*)-1-((2*S*,4*R*)-4-hydroxy-2-(((*S*)-1-(4-(4-methylthiazol-5-yl)phenyl)ethyl)carbamoyl)pyrrolidin-1-yl)-3,3-dimethyl-1-oxobutan-2-yl)heptanediamide (VHL-SNAP2-5C)

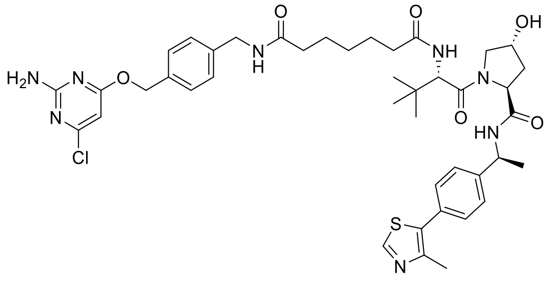

**2c** (17 mg, 0.03 mmol) and SNAP2 ligand were used to synthesise **VHL-SNAP2-5C** according to general procedure C. The reaction was stirred for 3 h. The crude product was purified by SFC (25-30% MeOH). **VHL-SNAP2-5C** was obtained as a white solid (10.8 mg, 46%). **$^1$H NMR** (500 MHz, Methanol-*d₄*) δ: 1.03 (s, 9H), 1.36 (h, *J* = 7.4 Hz, 2H), 1.50 (d, *J* = 7.0 Hz, 3H), 1.54–1.72 (m, 4H), 1.95 (ddd, *J* = 13.2, 9.0, 4.5 Hz, 1H), 2.14–2.34 (m, 5H), 2.47 (s, 3H), 2.66 (s, 1H), 3.74 (dd, *J*

= 11.0, 4.0 Hz, 1H), 3.87 (d, *J* = 11.1 Hz, 1H), 4.34 (d, *J* = 14.8 Hz, 2H), 4.40–4.45 (m, 1H), 4.53–4.60 (m, 1H), 4.62 (s, 1H), 5.00 (q, *J* = 7.1 Hz, 1H), 5.32 (s, 2H), 6.08 (s, 1H), 7.29 (d, *J* = 8.1 Hz, 2H), 7.37–7.46 (m, 6H), 8.86 (s, 1H). **$^{13}$C NMR** (126 MHz, Methanol-*d₄*) δ: 15.8, 22.4, 26.6, 26.7, 27.1, 29.8, 36.4, 36.5, 36.9, 38.8, 40.4, 43.8, 49.9, 50.1, 56.9, 58.0, 59.0, 60.6, 69.0, 71.0, 96.6, 127.6, 128.7, 129.6, 130.5, 131.5, 136.7, 140.2, 145.7, 148.9, 152.9, 161.8, 172.4, 173.2, 175.9, 176.0. **MS** (m/z): $[M + H]^+$ calcd. for $C_{42}H_{54}ClN_8O_6S$, 833.358; found 833.357.

#### ***N*¹-(4-(((2-amino-6-chloropyrimidin-4-yl)oxy)methyl)benzyl)-*N*⁸-((*S*)-1-((2*S*,4*R*)-4-hydroxy-2-(((*S*)-1-(4-(4-methylthiazol-5-yl)phenyl)ethyl)carbamoyl)pyrrolidin-1-yl)-3,3-dimethyl-1-oxobutan-2-yl)octanediamide (VHL-SNAP2-6C)**

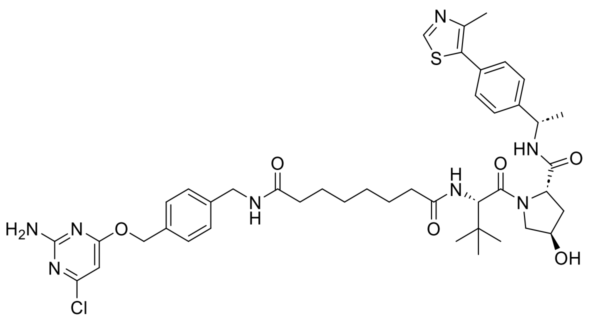

**2d** (17 mg, 0.03 mmol) and SNAP2 ligand were used to synthesise **VHL-SNAP2-6C** according to general procedure C. The reaction was stirred for 3 h. The crude product was purified by preparative HPLC (acidic, 15-55% B in A over 20 min). **VHL-SNAP2-6C** was obtained as a fluffy white solid (5.7 mg, 21%). **$^1$H NMR** (500 MHz, Methanol-*d₄*) δ: 1.03 (s, 9H), 1.31–1.37 (m, 4H), 1.50 (d, *J* = 7.1 Hz, 3H), 1.54–1.68 (m, 4H), 1.88–1.99 (m, 1H), 2.13–2.33 (m, 5H), 2.47 (s, 3H), 3.74 (dd, *J* = 11.0, 4.0 Hz, 1H), 3.85–3.91 (m, 1H), 4.36 (s, 2H), 4.42 (tt, *J* = 4.3, 2.0 Hz, 1H), 4.54–4.65 (m, 2H), 5.00 (q, *J* = 7.0 Hz, 1H), 5.32 (s, 2H), 6.08 (s, 1H), 7.26–7.3 (m, 2H), 7.36–7.45 (m, 6H), 8.87 (s, 1H). **$^{13}$C NMR** (126 MHz, Methanol-*d₄*) δ: 15.8, 22.4, 26.9, 26.9, 27.0, 29.9, 29.9, 36.4, 36.5, 37.0, 38.8, 43.8, 50.1, 58.0, 59.0, 60.6, 69.0, 71.0, 96.6, 127.6, 128.7, 129.6, 130.5, 131.5, 133.4, 136.7, 140.2, 145.7, 149.0, 152.9, 161.7, 164.4, 172.3, 173.3, 176.0, 176.1. **MS** (m/z): $[M + H]^+$ calcd. for $C_{43}H_{56}ClN_8O_6S$, 847.373; found 847.372.

#### ***N*¹-(4-(((4-Aminopyrimidin-2-yl)oxy)methyl)benzyl)-*N*⁶-((*S*)-1-((2*S*,4*R*)-4-hydroxy-2-(((*S*)-1-(4-(4-methylthiazol-5-yl)phenyl)ethyl)carbamoyl)pyrrolidin-1-yl)-3,3-dimethyl-1-oxobutan-2-yl)adipamide (VHL-CLIP-4C)**

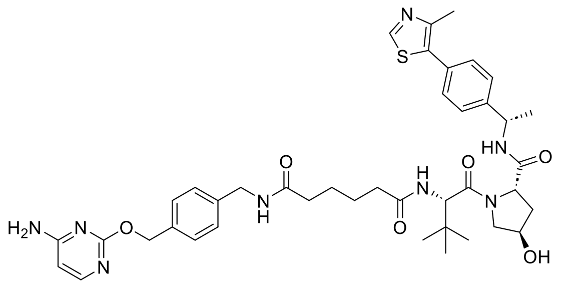

**2b** (49 mg, 0.08 mmol) and CLIP ligand were used to synthesise **VHL-CLIP-4C** according to general procedure B. The reaction was stirred for 3 h. The crude product was purified by preparative HPLC (acidic, 15-55% B in A over 20 min). **VHL-CLIP-4C** was obtained as a fluffy white solid (21.5 mg, 34%). **$^1$H NMR** (500 MHz, Methanol-*d₄*) δ 1.04 (s, 9H), 1.50 (d, *J* = 6.9 Hz, 3H), 1.60–1.69 (m, 4H), 1.95 (ddd, *J* = 13.3, 8.9, 4.5 Hz, 1H), 2.17 (d, *J* = 8.0 Hz, 1H), 2.23–2.38 (m, 4H), 2.47 (d, *J* = 9.5 Hz, 3H), 3.69–3.94 (m, 2H), 4.36 (d, *J* = 5.0 Hz, 2H), 4.42 (s, 1H), 4.53–4.64 (m, 2H), 5.00 (p, *J* = 6.5 Hz, 1H), 5.32 (s, 2H), 6.16 (d, *J* = 6.0 Hz, 1H), 7.28 (d, *J* = 7.9 Hz, 2H), 7.30–7.38 (m, 1H), 7.38–7.47 (m, 6H), 7.86 (dd, *J* = 17.4, 7.4 Hz, 2H), 8.40 (t, *J* = 5.9 Hz, 1H), 8.56 (d, *J* = 7.5 Hz, 1H), 8.87 (d, *J* = 7.8 Hz, 1H). **$^{13}$C NMR** (126 MHz, Methanol-*d₄*) δ 175.7, 175.7, 173.3, 172.3, 167.4, 165.5, 155.9, 152.9, 149.1, 145.7, 139.9, 137.2, 133.4, 131.5, 130.5, 129.3, 128.7, 127.6, 127.4, 100.3, 71.0, 69.3, 60.6, 59.1, 58.0, 50.1, 43.8, 38.8, 36.7, 36.4, 36.2, 27.1, 26.6, 26.5, 22.4, 15.8. **MS** (m/z): $[M + H]^+$ calcd. for $C_{41}H_{53}N_8O_6S$, 785.381; found 785.385.

#### ***N*[1]-(4-(((4-Aminopyrimidin-2-yl)oxy)methyl)benzyl)-*N*[7]-((*S*)-1-((2*S*,4*R*)-4-hydroxy-2-(((*S*)-1-(4-(4-methylthiazol-5-yl)phenyl)ethyl)carbamoyl)pyrrolidin-1-yl)-3,3-dimethyl-1-oxobutan-2-yl)heptanediamide (VHL-CLIP-5C)**

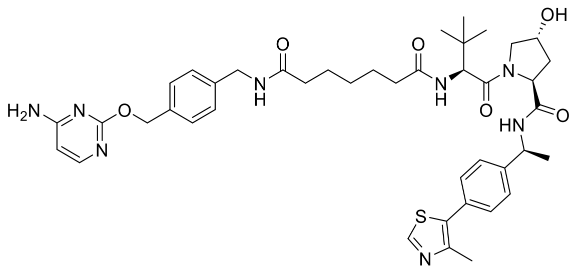

**2c** (50 mg, 0.08 mmol) and CLIP ligand were used to synthesise **VHL-CLIP-5C** according to general procedure B. The reaction was stirred or 3 h. The crude product was purified by preparative HPLC (acidic, 15-55% B in A over 20 min). **VHL-CLIP-5C** was obtained as a fluffy white solid (20 mg, 31%). **$^{1}$H NMR** (500 MHz, Methanol-*d$_4$*) δ 1.02 (d, *J* = 13.7 Hz, 9H), 1.36 (h, *J* = 7.7 Hz, 2H), 1.50 (d, *J* = 7.0 Hz, 3H), 1.57–1.73 (m, 4H), 1.95 (dtd, *J* = 13.3, 8.8, 4.6 Hz, 1H), 2.14–2.34 (m, 5H), 2.48 (s, 3H), 3.67–3.95 (m, 2H), 4.28–4.44 (m, 3H), 4.51–4.68 (m, 2H), 4.94–5.05 (m, 1H), 5.31 (s, 2H), 6.14 (d, *J* = 5.8 Hz, 1H), 7.28 (d, *J* = 8.0 Hz, 2H), 7.39–7.47 (m, 6H), 7.85 (d, *J* = 9.4 Hz, 2H), 8.39 (d, *J* = 6.4 Hz, 1H), 8.57 (d, *J* = 7.5 Hz, 2H), 8.88 (s, 1H). **$^{13}$C NMR** (126 MHz, Methanol-*d$_4$*) δ 176.0, 175.9, 173.3, 172.3, 167.3, 166.0, 157.0, 152.9, 149.1, 145.7, 139.8, 137.5, 133.4, 131.5, 130.5, 129.2, 128.6, 128.6, 127.6, 127.4, 100.3, 71.0, 69.1, 60.6, 59.0, 58.0, 50.1, 43.8, 38.8, 36.9, 36.4, 36.4, 29.8, 29.7, 27.1, 26.8, 26.7, 26.6, 22.4, 15.8. **MS** (m/z): $[M + H]^+$ calcd. for $C_{42}H_{55}N_8O_6S$, 799.397; found 799.397.

#### ***N*[1]-(4-(((4-Aminopyrimidin-2-yl)oxy)methyl)benzyl)-*N*[8]-((*S*)-1-((2*S*,4*R*)-4-hydroxy-2-(((*S*)-1-(4-(4-methylthiazol-5-yl)phenyl)ethyl)carbamoyl)pyrrolidin-1-yl)-3,3-dimethyl-1-oxobutan-2-yl)octanediamide (VHL-CLIP-6C)**

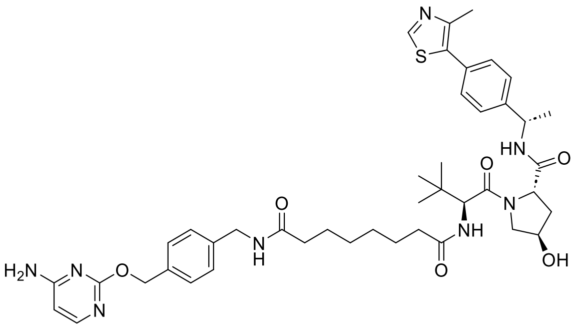

**2d** (51 mg, 0.08 mmol) and CLIP ligand were used to synthesise **VHL-CLIP-6C** according to general procedure B. The reaction was stirred for 3 h. The crude product was purified by preparative HPLC (acidic, 15-55% B in A over 20 min). **VHL-CLIP-6C** was obtained as a fluffy white solid (19.5 mg, 30%). **$^{1}$H NMR** (500 MHz, Methanol-*d$_4$*) δ: 1.04 (s, 9H), 1.32–1.39 (m, 4H), 1.52 (d, *J* = 7.0 Hz, 3H), 1.62 (ddd, *J* = 21.3, 14.6, 7.3 Hz, 4H), 1.96 (ddd, *J* = 13.2, 9.0, 4.5 Hz, 1H), 2.15–2.36 (m, 5H), 2.49 (s, 3H), 2.67 (s, 1H), 3.76 (dd, *J* = 11.0, 4.0 Hz, 1H), 3.90 (d, *J* = 11.0 Hz, 1H), 4.37 (s, 2H), 4.42–4.46 (m, 1H), 4.59 (dd, *J* = 10.0, 6.6 Hz, 1H), 4.62–4.67 (m, 1H), 5.02 (q, *J* = 7.0 Hz, 1H), 5.31 (d, *J* = 13.5 Hz, 2H), 6.16 (d, *J* = 5.9 Hz, 1H), 7.29 (d, *J* = 8.1 Hz, 2H), 7.40–7.48 (m, 6H), 7.86 (d, *J* = 5.9 Hz, 1H), 8.88 (s, 1H). **$^{13}$C NMR** (126 MHz, Methanol-*d$_4$*) δ 176.1, 176.0, 173.3, 172.3, 167.3, 165.8, 156.6, 152.9, 149.1, 145.7, 139.9, 137.4, 133.4, 131.5, 130.5, 129.2, 128.6, 128.6, 127.6, 127.4, 100.3, 71.0, 69.2, 60.6, 59.0, 58.0, 50.1, 43.8, 38.8, 37.0, 36.5, 36.4, 29.9, 29.9, 27.0, 26.9, 26.9, 22.4, 15.8. **MS** (m/z): $[M + H]^+$ calcd. for $C_{43}H_{57}N_8O_6S$, 813.412; found 813.413.

### Synthesis of control compounds (VHL ligand, SNAP ligand, CLIP ligand)

#### (2*S*,4*R*)-1-((*S*)-2-Acetamido-3,3-dimethylbutanoyl)-4-hydroxy-N-((*S*)-1-(4-(4-methylthiazol-5-yl)phenyl)ethyl)pyrrolidine-2-carboxamide (VHL ligand)

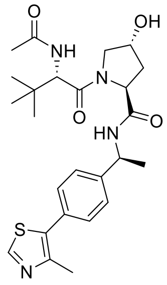

**1** (50 mg, 0.10 mmol), acetic acid (6.54 µL, 0.11 mmol, 1.1. eq.) and HATU (47 mg, 0.12 mmol, 1.2 eq.) were taken up in DMF (1 mL) and DIPEA (91 µl, 0.52 mmol, 5 eq.) added. The reaction was stirred at room temperature for 3 h. The mixture was diluted in EtOAc, washed once with saturated $NaHCO_3$ and extracted twice with EtOAc. The combined organic layer was dried over $MgSO_4$, filtered, and concentrated. The crude product was purified by NPFCC (5 g, 2-15% MeOH in EtOAC over 7 min). **VHL ligand** was obtained as white crystals (26.3 mg, 52%) $^1$**H NMR** (500 MHz, Methanol-*d₄*) δ: 1.05 (s, 9H), 1.50 (d, *J* = 7.2 Hz, 3H), 1.96 (ddd, *J* = 13.3, 9.0, 4.5 Hz, 1H), 2.00 (s, 3H), 2.19 (ddt, *J* = 13.2, 7.8, 2.0 Hz, 1H), 2.47 (s, 3H), 3.35 (s, 1H), 3.75 (dd, *J* = 11.0, 4.0 Hz, 1H), 3.88 (dt, *J* = 11.2, 1.8 Hz, 1H), 4.40–4.46 (m, 1H), 4.58 (t, *J* = 9.5, 7.1 Hz, 1H), 4.62 (d, *J* = 8.9 Hz, 1H), 5.01 (q, 1H), 7.40–7.46 (m, 4H), 7.94 (d, *J* = 8.9 Hz, 1H), 8.55 (d, *J* = 7.5 Hz, 1H), 8.87 (s, 1H).$^{13}$**C NMR** (126 MHz, Methanol-$d_4$) δ: 14.5, 15.8, 20.9, 27.0, 36.4, 38.8, 50.2, 58.0, 59.3, 61.5, 71.0, 127.6, 130.5, 131.5, 133.3, 145.7, 149.1, 152.8, 172.3, 173.2, 173.3. **MS** (m/z): $[M + H]^+$ calcd. for $C_{25}H_{35}N_4O_4S$, 487.238; found 487.240.

#### *N*-(4-(((2-Amino-6-chloropyrimidin-4-yl)oxy)methyl)benzyl)acetamide (SNAP ligand)

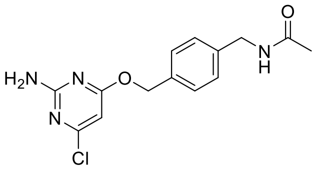

SNAP2 ligand (30 mg, 0.11 mmol), acetic acid (5.30 µl, 0.09 mmol, 0.9 eq.) and HATU (39.2 mg, 0.10 mmol, 1.0 eq.) were taken up in DMF (1.0 mL) and DIPEA (90 µl, 0.52 mmol, 5 eq.)) added. The reaction was stirred at room temperature for 3 h. The mixture was diluted in EtOAc, washed with saturated $NaHCO_3$ and extracted twice with EtOAc. The combined organic layer was dried with $MgSO_4$, filtered, and concentrated. The crude product was purified by preparative HPLC (basic, 15-20% B in A over 20 min). **SNAP ligand** was obtained as a white solid (12.4 mg, 39%). $^1$**H NMR** (500 MHz, Methanol-$d_4$) 2.00 (s, 3H), 4.37 (s, 2H), 5.36 (s, 2H), 6.12 (s, 1H), 7.31 (d, J = 8.1 Hz, 2H), 7.41 (d, J = 8.0 Hz, 2H). $NH_2$ not observed. $^{13}$**C NMR** (126 MHz, Methanol-$d_4$) δ 173.1, 172.4, 164.5, 161.8, 140.0, 136.8, 129.6, 128.7, 96.6, 69.0, 43.9, 22.5. **MS** (m/z): $[M + H]^+$ calcd. for $C_{14}H_{16}ClN_4O_2$, 307.096; found 307.098.

#### *N*-(4-(((4-Aminopyrimidin-2-yl)oxy)methyl)benzyl)acetamide (CLIP ligand)

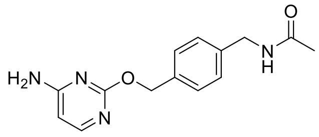

CLIP ligand (30 mg, 0.13 mmol), acetic acid (6.10 µl, 0.11 mmol, 0.9 eq.) and HATU (45.0 mg, 0.12 mmol, 1.0 eq.) was taken up in DMF (1.2 mL) and DIPEA (103 µl, 0.59 mmol, 5 eq.)) added. The reaction was stirred at room temperature for 3 h. The mixture was diluted in EtOAc, washed with saturated $NaHCO_3$ and extracted twice with EtOAc. The organic layer was dried with $MgSO_4$, filtered, and concentrated. The crude product was purified by preparative HPLC (basic, 5-45% B in A over 20 min). **CLIP ligand** was obtained as a white solid (14.2 mg, 44%). $^1$**H NMR** (500 MHz, Methanol-$d_4$) 1.99 (s, 3H), 4.36 (s, 2H), 5.32 (s, 2H), 6.15 (d, J = 5.9 Hz, 1H), 7.29 (d, J = 8.1 Hz, 2H), 7.39–7.45 (m, 2H), 7.86 (d, J = 5.9 Hz, 1H). $^{13}$**C NMR** (126 MHz, Methanol-$d_4$) 22.5, 44.0, 69.0, 100.3, 128.7, 129.2, 137.5, 139.6, 157.1, 166.1, 167.3, 173.1. **MS** (m/z): $[M + H]^+$ calcd. for $C_{14}H_{16}N_4O_2$, 273.1; found 273.1.

### CRBN recruiting PROTACs

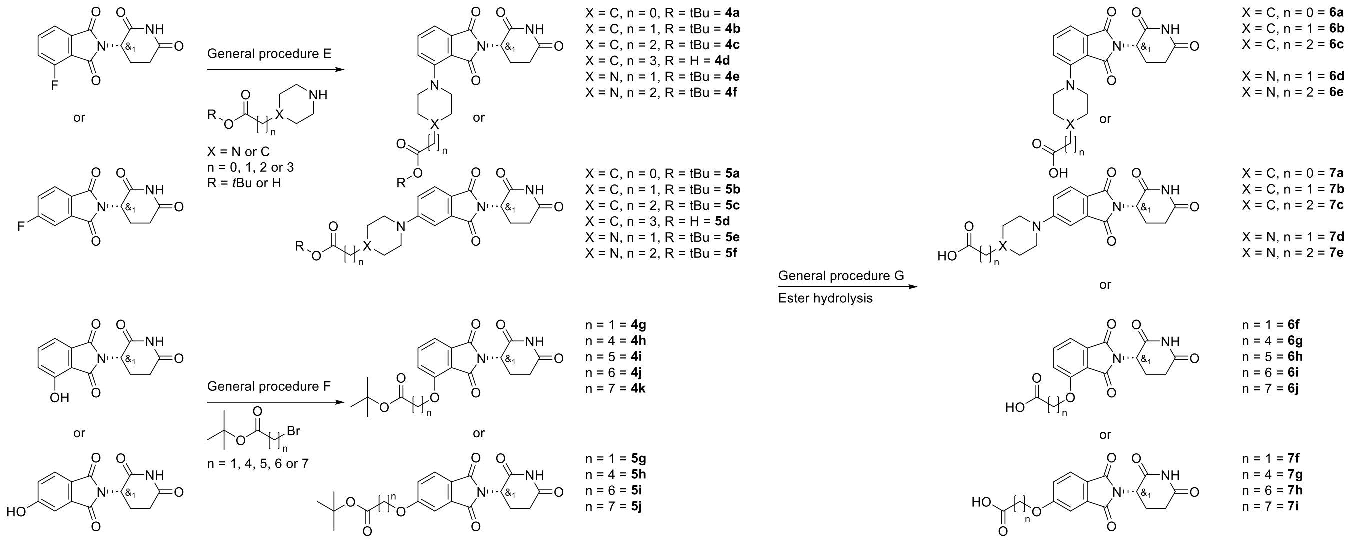

**Scheme S2**. Overview of synthetic route for CRBN intermediates.

#### General procedure E

*Rac*-2-(2,6-dioxopiperidin-3-yl)-4-fluoroisoindoline-1,3-dione (1.0 eq.) or *rac*-2-(2,6-dioxopiperidin-3-yl)-5-fluoroisoindoline-1,3-dione (1.0 eq.) and the corresponding amine (1.0 eq.) were dissolved in DMSO (3 mL) and DIPEA (1.2 eq.) added. The reaction was stirred at 100 °C until completion monitored by LCMS. The reaction mixture was diluted with EtOAc and was washed twice with brine. The combined organic layer was dried over $MgSO_4$, filtered, concentrated, and purified by NPFCC to afford products **4a-4f and 5a-5f**.

#### Rac-*tert*-butyl-1-(2-(2,6-dioxopiperidin-3-yl)-1,3-dioxoisoindolin-4-yl)piperidine-4-carboxylate (4a)

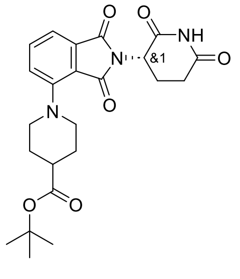

*Rac*-2-(2,6-dioxopiperidin-3-yl)-4-fluoroisoindoline-1,3-dione (300 mg, 1.09 mmol) and *tert*-butyl piperidine-4-carboxylate hydrochloride (1.3 eq.) were used to synthesise **4a** according to general procedure E. The reaction was stirred overnight. The crude was purified by NPFCC (10 g, 0-100% EtOAc in heptane over 10 CV) to afford **4a** as a yellow powder (371 mg, 77%). **$^1$H NMR** (500 MHz, Chloroform-*d*) δ 1.47 (s, 9H), 2.07–2.15 (m, 1H), 2.71–2.89 (m, 3H), 3.46 (tt, *J* = 8.9, 6.2 Hz, 1H), 4.33 (ddd, *J* = 8.7, 6.3, 4.1 Hz, 2H), 4.45 (t, *J* = 8.8 Hz, 2H), 4.92 (dd, *J* = 12.4, 5.4 Hz, 1H), 6.62 (d, *J* = 8.6 Hz, 1H), 7.19 (d, *J* = 7.1 Hz, 1H), 7.45–7.49 (m, 1H), 8.05 (s, 1H). **MS** (m/z): $[M + H]^+$ calcd. for $C_{23}H_{28}N_3O_6$, 442.2; found 442.3.

#### *Rac-tert*-butyl-2-(1-(2-(2,6-dioxopiperidin-3-yl)-1,3-dioxoisoindolin-4-yl)piperidin-4-yl)acetate (4b)

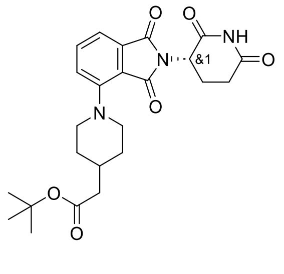

*Rac*-2-(2,6-dioxopiperidin-3-yl)-4-fluoroisoindoline-1,3-dione (300 mg, 1.09 mmol) and *tert*-butyl 2-(piperidin-4-yl)acetate (1.0 eq.) were used to synthesise **4b** according to general procedure E. The reaction was stirred for 1 h, and instead of EtOAc, DCM was used for the extraction. The crude was purified by NPFCC (10 g, 0-100% EtOAc in heptane over 10 CV) to afford **4a** as a yellow film (218 mg, 44%). **$^1$H NMR** (500 MHz, Chloroform-*d*) δ 1.46 (s, 9H), 1.64 (s, 2H), 1.89 (d, *J* = 12.2 Hz,

2H), 2.01 (s, 1H), 2.09–2.16 (m, 1H), 2.24 (d, *J* = 7.1 Hz, 2H), 2.68–2.77 (m, 1H), 2.81 (td, *J* = 12.7, 3.9 Hz, 1H), 2.86–2.92 (m, 1H), 3.06 (s, 1H), 3.71 (t, *J* = 11.3 Hz, 2H), 4.96 (dd, *J* = 12.4, 5.3 Hz, 1H), 5.30 (s, 2H), 7.45 (s, 1H), 7.61 (t, *J* = 7.6 Hz, 1H), 8.08 (s, 1H). **MS** (m/z): [M + H]$^+$ calcd. for $C_{24}H_{30}N_3O_6$, 456.2; found 456.5.

#### ***Rac-tert*-butyl-3-(1-(2-(2,6-dioxopiperidin-3-yl)-1,3-dioxoisoindolin-4-yl)piperidin-4-yl)propanoate (4c)**

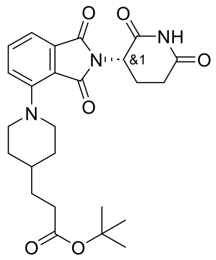

*Rac*-2-(2,6-dioxopiperidin-3-yl)-4-fluoroisoindoline-1,3-dione (300 mg, 1.09 mmol) and *tert*-butyl 3-(piperidin-4-yl)propanoate (1.0 eq.) were used to synthesise **4c** according to general procedure E. The reaction was stirred overnight. The crude was purified by NPFCC (25 g, 40-50% EtOAc in heptane over 10 CV) to afford **4c** as a yellow powder (447 mg, 88%).

**$^1$H NMR** (500 MHz, DMSO-*d$_6$*) δ 1.32 (q, *J* = 12.2 Hz, 2H), 1.39 (s, 10H), 1.50 (q, *J* = 7.2 Hz, 2H), 1.76 (d, *J* = 13.3 Hz, 2H), 2.03 (ddd, *J* = 9.8, 6.1, 2.0 Hz, 1H), 2.25 (t, *J* = 7.6 Hz, 2H), 2.51–2.62 (m, 2H), 2.79–2.92 (m, 3H), 3.68 (d, *J* = 11.8 Hz, 2H), 5.09 (dd, *J* = 12.7, 5.4 Hz, 1H), 7.32 (dd, *J* = 7.8, 5.3 Hz, 2H), 7.67 (dd, *J* = 8.4, 7.1 Hz, 1H), 11.09 (s, 1H). **MS** (m/z): [M + H]$^+$ calcd. for $C_{25}H_{32}N_3O_6$, 470.2; found 470.3.

#### ***Rac*-4-(1-(2-(2,6-dioxopiperidin-3-yl)-1,3-dioxoisoindolin-4-yl)piperidin-4-yl)butanoic acid (4d)**

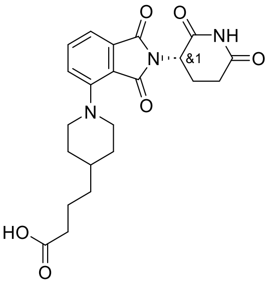

*Rac*-2-(2,6-dioxopiperidin-3-yl)-4-fluoroisoindoline-1,3-dione (300 mg, 1.09 mmol) and 4-(piperidin-4-yl)butanoic acid (1.2 eq.) were used to synthesise **4d** according to general procedure E. The reaction was stirred overnight. The crude was purified by RPFCC (30 g, 10-95% B in A over 10 CV) to afford **4d** as a yellow/orange film/foam (365 mg, 79%). **$^1$H NMR** (500 MHz, DMSO-*d$_6$*) δ 1.23–1.36 (m, 4H), 1.42 (ddq, *J* = 10.7, 7.3, 3.9 Hz, 1H), 1.55 (p, *J* = 7.5 Hz, 2H), 1.76 (d, *J* = 16.3 Hz, 2H), 2.00–2.04 (m, 1H), 2.21 (t, *J* = 7.3 Hz, 2H), 2.51–2.62 (m, 2H), 2.84 (t, *J* = 12.1 Hz, 3H), 3.68 (d, *J* = 12.4 Hz, 2H), 5.09 (dd, *J* = 12.8, 5.4 Hz, 1H), 7.32 (t, *J* = 7.2 Hz, 2H), 7.64–7.69 (m, 1H), 11.09 (s, 1H), 12.01 (s, 1H). ). **MS** (m/z): [M + H]$^+$ calcd. for $C_{22}H_{26}N_3O_6$, 428.2; found 428.2.

#### ***Rac-tert*-butyl-2-(4-(2-(2,6-dioxopiperidin-3-yl)-1,3-dioxoisoindolin-4-yl)piperazin-1-yl)acetate (4e)**

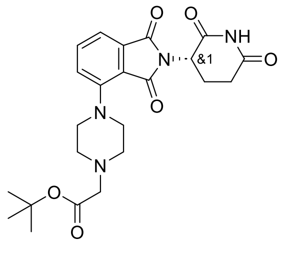

*Rac*-2-(2,6-dioxopiperidin-3-yl)-4-fluoroisoindoline-1,3-dione (300 mg, 1.09 mmol) and *tert*-butyl 2-(piperazin-1-yl)acetate (1.2 eq.) were used to synthesise **4e** according to general procedure **E**. Reaction was stirred for 4 h. The crude was purified by RPFCC (30 g, 20-50% B in A over 10 CV) to afford **4e** as a yellow powder (270 mg, 55%). **$^1$H NMR** (500 MHz, DMSO-*d$_6$*) δ 1.42 (s, 9H), 2.02 (ddd, *J* = 12.8, 5.6, 3.3 Hz, 1H), 2.53–2.62 (m, 2H), 2.70 (s, 4H), 2.87 (ddd, *J* = 16.7, 13.7, 5.4 Hz, 1H), 3.18 (s, 2H), 3.29 (s, 4H), 5.09 (dd, *J* = 12.8, 5.5 Hz, 1H), 7.35 (t, *J* = 7.6 Hz, 2H), 7.68–7.72 (m, 1H), 11.09 (s, 1H). **MS** (m/z): [M + H]$^+$ calcd. For $C_{23}H_{29}N_4O_6$, 457.2; found 457.3.

#### ***Rac-tert*-butyl-(*R*)-3-(4-(2-(2,6-dioxopiperidin-3-yl)-1,3-dioxoisoindolin-4-yl)piperazin-1-yl)propanoate (4f)**

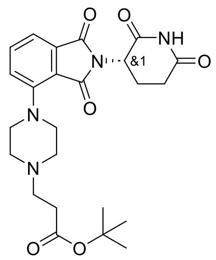

*Rac*-2-(2,6-dioxopiperidin-3-yl)-4-fluoroisoindoline-1,3-dione (300 mg, 1.09 mmol) and *tert*-butyl 3-(piperazin-1-yl)propanoate (1.0 eq.) were used to synthesise **4f** according to general procedure E. The reaction was stirred overnight. The crude was purified by RPFCC (30 g, 5-50% B in A over 10 CV) to afford **4f** as yellow solid (409 mg, 80%). **$^1$H NMR** (500 MHz, DMSO-$d_6$) δ 1.41 (s, 9H), 2.01–2.04 (m, 1H), 2.39 (t, *J* = 7.1 Hz, 2H), 2.52–2.63 (m, 8H), 2.87 (ddd, *J* = 16.8, 13.8, 5.4 Hz, 1H), 3.27 (s, 4H), 5.09 (dd, *J* = 12.8, 5.5 Hz, 1H), 7.32–7.38 (m, 2H), 7.66–7.73 (m, 1H), 11.09 (s, 1H). **MS** (m/z): $[M + H]^+$ calcd. for $C_{24}H_{31}N_4O_6$, 471.2; found 471.4.

**_Rac-tert_-butyl-1-(2-(2,6-dioxopiperidin-3-yl)-1,3-dioxoisoindolin-5-yl)piperidine-4-carboxylate (5a)**

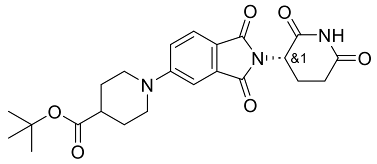

*Rac*-2-(2,6-Dioxopiperidin-3-yl)-5-fluoroisoindoline-1,3-dione (300 mg, 1.09 mmol) and *tert*-butyl piperidine-4-carboxylate hydrochloride (1.3 eq.) were used to synthesise **5a** according to general procedure E. The reaction was stirred overnight. The crude was purified by RPFCC (30 g, 10-95% B in A over 10 CV) to afford **5a** as a yellow/orange foam/film (331 mg, 46%). **$^1$H NMR** (500 MHz, Chloroform-*d*) δ 1.47 (s, 9H), 1.92–2.01 (m, 2H), 2.13 (ddt, *J* = 9.9, 4.4, 2.1 Hz, 1H), 2.20 (t, *J* = 8.1 Hz, 2H), 2.56 (tt, *J* = 9.1, 4.2 Hz, 1H), 2.71–2.92 (m, 3H), 3.19 (t, *J* = 9.7 Hz, 2H), 3.83 (dt, *J* = 12.7, 4.7 Hz, 2H), 4.94–4.99 (m, 1H), 7.51 (s, 2H), 7.77 (d, *J* = 9.0 Hz, 1H), 8.11 (s, 1H). **MS** (m/z): $[M + H]^+$ calcd. for $C_{23}H_{28}N_3O_6$, 442.20; found 442.42.

**_Rac-tert_-butyl-2-(1-(2-(2,6-dioxopiperidin-3-yl)-1,3-dioxoisoindolin-5-yl)piperidin-4-yl)acetate (5b)**

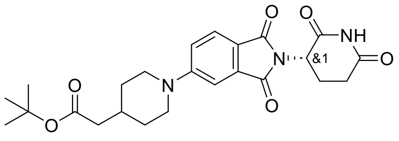

*Rac*-2-(2,6-dioxopiperidin-3-yl)-5-fluoroisoindoline-1,3-dione (436 mg, 1.58 mmol) and *tert*-butyl 2-(piperidin-4-yl)acetate (1.0 eq.) were used to synthesise **5b** according to general procedure E. The reaction was stirred for 2 h. The crude was purified by NPFCC (5 g, 0-100% EtOAc in heptane over 10 CV) to afford **5b** as a yellow foam/film (331 mg, 46%). **$^1$H NMR** (500 MHz, Chloroform-*d*) δ 1.46 (s, 9H), 1.72 (q, *J* = 11.1 Hz, 2H), 1.93 (d, *J* = 10.4 Hz, 2H), 2.15 (td, *J* = 5.0, 2.4 Hz, 1H), 2.25 (d, *J* = 7.1 Hz, 2H), 2.73–2.92 (m, 4H), 3.08–3.20 (m, 2H), 3.86 (d, *J* = 12.7 Hz, 2H), 4.96 (dd, *J* = 12.4, 5.3 Hz, 1H), 7.55 (s, 1H), 7.60 (s, 1H), 7.79 (d, *J* = 8.4 Hz, 1H), 8.10 (s, 1H). **MS** (m/z): $[M + H]^+$ calcd. for $C_{24}H_{30}N_3O_6$, 456.2; found 456.4.

**_Rac-tert_-butyl-3-(1-(2-(2,6-dioxopiperidin-3-yl)-1,3-dioxoisoindolin-5-yl)piperidin-4-yl)propanoate (5c)**

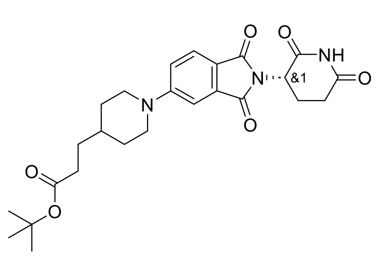

*Rac*-2-(2,6-dioxopiperidin-3-yl)-5-fluoroisoindoline-1,3-dione (300 mg, 1.09 mmol) and *tert*-butyl 3-(piperidin-4-yl)propanoate (1.0 eq.) were used to synthesise **5c** according to general procedure E. The reaction was stirred overnight. The crude was purified by NPFCC (25 g, 40-50% EtOAc in heptane over 10 CV) to afford **5c** as a yellow foam (451 mg, 88%). **$^1$H NMR** (500 MHz, Chloroform-*d*) δ 1.45 (s, 9H), 1.68 (t, *J* = 5.8 Hz, 3H), 1.97 (s, 4H), 2.15 (ddd, *J* = 7.3, 5.3, 2.2 Hz, 1H), 2.27–2.31 (m, 2H), 2.71–2.94 (m, 3H), 3.15–3.26 (m, 2H), 3.80 (d, *J* = 12.2 Hz, 2H), 4.97 (dd, *J* = 12.6, 5.4 Hz, 1H), 7.80 (s, 1H), 7.88 (d, *J* = 8.2 Hz, 1H), 8.08 (d, *J* = 7.4 Hz, 1H), 8.17 (s, 1H). **MS** (m/z): $[M + H]^+$ calcd. for $C_{25}H_{32}N_3O_6$, 470.2; found 470.4.

**_Rac_-4-(1-(2-(2,6-dioxopiperidin-3-yl)-1,3-dioxoisoindolin-5-yl)piperidin-4-yl)butanoic acid (5d)**

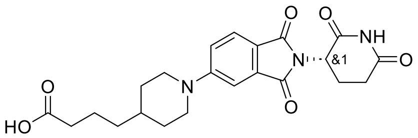

*Rac*-2-(2,6-dioxopiperidin-3-yl)-5-fluoroisoindoline-1,3-dione (300 mg, 1.09 mmol) and 4-(piperidin-4-yl)butanoic acid (1.2 eq.) were used to synthesise **5d** according to general procedure E. The reaction was stirred overnight. The crude product was purified by RPFCC (30 g, 10-95% B in A over 10 CV) to afford **5d** as a yellow/orange film/foam (325 mg, 70%). **$^1$H NMR** (500 MHz, DMSO-*d*$_6$) δ 1.14 (qd, *J* = 12.9, 4.1 Hz, 2H), 1.19–1.25 (m, 2H), 1.48–1.58 (m, 3H), 1.74 (d, *J* = 13.8 Hz, 2H), 1.97–2.04 (m, 1H), 2.20 (t, *J* = 7.4 Hz, 2H), 2.51–2.62 (m, 2H), 2.83–2.98 (m, 3H), 4.04 (d, *J* = 13.3 Hz, 2H), 5.06 (dd, *J* = 12.8, 5.4 Hz, 1H), 7.22 (dd, *J* = 8.7, 2.4 Hz, 1H), 7.30 (d, *J* = 2.3 Hz, 1H), 7.64 (d, *J* = 8.5 Hz, 1H), 11.08 (s, 1H), 12.00 (s, 1H). **MS** (m/z): [M + H]$^+$ calcd. for $C_{22}H_{26}N_3O_6$, 428.18; found 428.18.

**_Rac-tert_-butyl-2-(4-(2-(2,6-dioxopiperidin-3-yl)-1,3-dioxoisoindolin-5-yl)piperazin-1-yl)acetate (5e)**

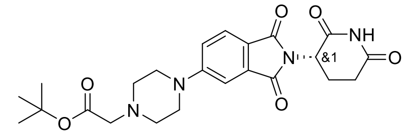

*Rac*-2-(2,6-dioxopiperidin-3-yl)-5-fluoroisoindoline-1,3-dione (300 mg, 1.09 mmol) and *tert*-butyl 2-(piperazin-1-yl)acetate (1.2 eq.) were used to synthesise **5e** according to general procedure E, and stirred for 4 h. The crude product was purified by NPFCC (10 g, 50-75% EtOAc in heptane over 10 CV) to afford **5e** as a yellow foam/film (286 mg, 58%). **$^1$H NMR** (500 MHz, Chloroform-*d*) δ 1.49 (s, 9H), 2.14 (ddt, *J* = 10.2, 5.2, 2.7 Hz, 1H), 2.74–2.91 (m, 3H), 3.22 (s, 4H), 3.52 (s, 2H), 3.73 (s, 4H), 4.95 (dd, *J* = 12.4, 5.3 Hz, 1H), 7.08 (dd, *J* = 8.5, 2.4 Hz, 1H), 7.30 (d, *J* = 2.4 Hz, 1H), 7.72 (d, *J* = 8.5 Hz, 1H), 8.19 (s, 1H). **MS** (m/z): [M + H]$^+$ calcd. for $C_{23}H_{29}N_4O_6$, 457.2; found 457.4.

**_Rac-tert_-butyl-3-(4-(2-(2,6-dioxopiperidin-3-yl)-1,3-dioxoisoindolin-5-yl)piperazin-1-yl)propanoate (5f)**

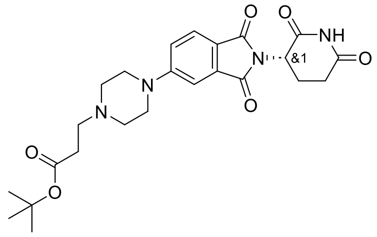

*Rac*-2-(2,6-dioxopiperidin-3-yl)-5-fluoroisoindoline-1,3-dione (300 mg, 1.09 mmol) and *tert*-butyl 3-(piperazin-1-yl)propanoate (1.0 eq.) was used to synthesise **5f** according to general procedure E. The reaction was stirred overnight. The crude product was purified by RPFCC (30 g, 20-50% B in A over 10 CV) to afford **5f** as a yellow foam/solid (459 mg, 90%). **$^1$H NMR** (500 MHz, Chloroform-*d*) δ 1.45 (s, 9H), 2.13 (ddt, *J* = 10.2, 5.2, 2.3 Hz, 1H), 2.73–2.90 (m, 5H), 3.03 (s, 4H), 3.09 (t, *J* = 7.1 Hz, 2H), 3.68 (t, *J* = 5.0 Hz, 4H), 4.95 (dd, *J* = 12.4, 5.3 Hz, 1H), 7.08 (dd, *J* = 8.5, 2.4 Hz, 1H), 7.31 (d, *J* = 2.3 Hz, 1H), 7.71 (d, *J* = 8.5 Hz, 1H), 8.38 (s, 1H). **MS** (m/z): [M + H]$^+$ calcd. for $C_{24}H_{31}N_4O_6$, 471.2; found 471.4.

**General procedure F**

*Rac*-2-(2,6-dioxopiperidin-3-yl)-4-hydroxyisoindoline-1,3-dione (1 eq.) or *rac*-2-(2,6-dioxopiperidin-3-yl)-5-hydroxyisoindoline-1,3-dione (1 eq.), the corresponding bromoalkane (1.1 eq.) and potassium hydrogen carbonate (1.65 eq.) were dissolved in DMF (5 mL) and stirred at 60 °C overnight. The amount of DMF was reduced by evaporation and the crude was purified by RPFCC to afford products **4h**-**4k** and **5g**-**5j**.

#### *Rac-tert*-butyl-2-((2-(2,6-dioxopiperidin-3-yl)-1,3-dioxoisoindolin-4-yl)oxy)acetate (4g)

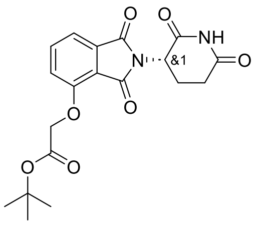


*Rac*-2-(2,6-dioxopiperidin-3-yl)-4-hydroxyisoindoline-1,3-dione (626 mg, 2.28 mmol), *tert*-butyl 2-bromoacetate (1.1 eq.) and potassium hydrogen carbonate (1.65 eq.) were dissolved in DMF (5 mL) and stirred at 60 °C overnight. The reaction mixture was diluted with EtOAc and washed with saturated ammonium chloride. The combined organic layer was dried over $MgSO_4$, filtered, concentrated, and the crude product was purified by NPFCC (25 g, 40-55% EtOAc in heptane over 10 CV) to afford **4g** as a white solid (369 mg, 42%). **$^1$H NMR** (500 MHz, Chloroform-*d*) δ 1.48 (s, 9H), 2.13 (ddt, *J* = 10.4, 5.4, 2.6 Hz, 1H), 2.70–2.94 (m, 3H), 4.79 (s, 2H), 4.97 (dd, *J* = 12.4, 5.3 Hz, 1H), 7.11 (d, *J* = 8.5 Hz, 1H), 7.52 (d, *J* = 7.3 Hz, 1H), 7.65–7.70 (m, 1H), 7.97 (s, 1H). **MS** (m/z): $[M - H]^-$ calcd. for $C_{19}H_{19}N_2O_7$, 387.1; found 387.1.

#### *Rac-tert*-butyl-5-((2-(2,6-dioxopiperidin-3-yl)-1,3-dioxoisoindolin-4-yl)oxy)pentanoate (4h)

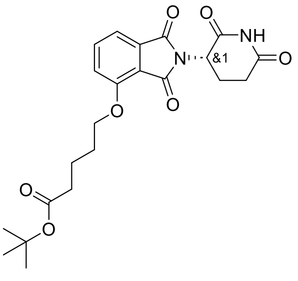


*Rac*-2-(2,6-dioxopiperidin-3-yl)-4-hydroxyisoindoline-1,3-dione (515 mg, 1.88 mmol) and *tert*-butyl 5-bromopentanoate (1.1 eq.) were used to synthesise **4h** according to general procedure F. The crude was purified by RPFCC (30 g, 30-50% B in A over 10 CV) to afford **4h** as a grey foam (662 mg, 82%). **$^1$H NMR** (500 MHz, Chloroform-*d*) δ 1.44 (s, 9H), 1.78–1.85 (m, 2H), 1.88–1.96 (m, 2H), 2.10–2.15 (m, 1H), 2.32 (t, *J* = 7.2 Hz, 2H), 2.69–2.93 (m, 3H), 4.19 (t, *J* = 6.4 Hz, 2H), 4.95 (dd, *J* = 12.4, 5.4 Hz, 1H), 7.21 (d, *J* = 8.5 Hz, 1H), 7.45 (d, *J* = 7.3 Hz, 1H), 7.65–7.70 (m, 1H), 8.00 (s, 1H). **MS** (m/z): $[M - H]^-$ calcd. for $C_{22}H_{25}N_2O_7$, 429.2; found 429.3.

#### *Rac-tert*-butyl-6-((2-(2,6-dioxopiperidin-3-yl)-1,3-dioxoisoindolin-4-yl)oxy)hexanoate (4i)

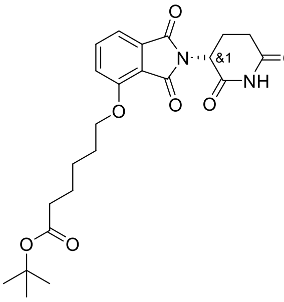


*Rac*-2-(2,6-dioxopiperidin-3-yl)-4-hydroxyisoindoline-1,3-dione (486 mg, 1.77 mmol) and *tert*-butyl 6-bromohexanoate (1.1 eq.) were used to synthesise **4i** according to general procedure F. The crude was purified by RPFCC (30 g, 40-70% B in A over 10 CV) to afford **4i** as a beige solid (587 mg, 75%). **$^1$H NMR** (500 MHz, Chloroform-*d*) δ 1.44 (s, 9H), 1.50–1.57 (m, 2H), 1.63–1.71 (m, 2H), 1.86–1.94 (m, 2H), 2.09–2.15 (m, 1H), 2.26 (t, *J* = 7.4 Hz, 2H), 2.69–2.93 (m, 3H), 4.18 (t, *J* = 6.5 Hz, 2H), 4.95 (dd, *J* = 12.3, 5.4 Hz, 1H), 7.21 (d, *J* = 8.5 Hz, 1H), 7.45 (d, *J* = 7.2 Hz, 1H), 7.67 (dd, *J* = 8.5, 7.3 Hz, 1H), 8.00 (s, 1H). **MS** (m/z): $[M - H]^-$ calcd. for $C_{23}H_{27}N_2O_7$, 443.2; found 443.4.

#### *Rac-tert*-butyl-(R)-7-((2-(2,6-dioxopiperidin-3-yl)-1,3-dioxoisoindolin-4-yl)oxy)heptanoate (4j)

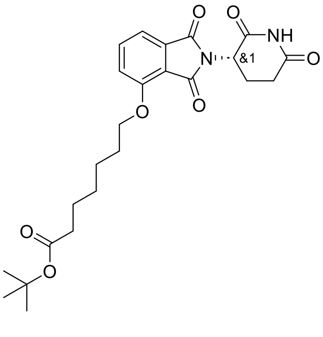


*Rac*-2-(2,6-dioxopiperidin-3-yl)-4-hydroxyisoindoline-1,3-dione (461 mg, 1.68 mmol) and *tert*-butyl 7-bromoheptanoate (1.1 eq.) were used to synthesise **4j** according to general procedure F. The crude was purified by RPFCC (30 g, 40-70% B in A over 10 CV) to afford **4j** as a white/light grey solid (576 mg, 75%). **$^1$H NMR** (500 MHz, Chloroform-*d*) δ 1.40 (p, *J* = 6.6 Hz, 2H), 1.44 (s, 9H), 1.50–1.56 (m, 2H), 1.62 (p, *J* = 7.4 Hz, 2H), 1.85–1.92 (m, 2H), 2.10–2.16 (m, 1H), 2.22 (t, *J* = 7.5 Hz, 2H), 2.70–2.93 (m, 3H), 4.17 (t, *J* = 6.6 Hz, 2H), 4.95 (dd, *J* = 12.3, 5.4 Hz, 1H), 7.21 (d, *J* = 8.5 Hz, 1H), 7.45 (d, *J* = 7.3 Hz, 1H), 7.64–7.70 (m, 1H), 7.99 (s, 1H). **MS** (m/z): $[M - H]^-$ calcd. for $C_{24}H_{29}N_2O_7$, 457.2; found 457.1.

#### ***Rac-tert*-butyl-8-((2-(2,6-dioxopiperidin-3-yl)-1,3-dioxoisoindolin-4-yl)oxy)octanoate (4k)**

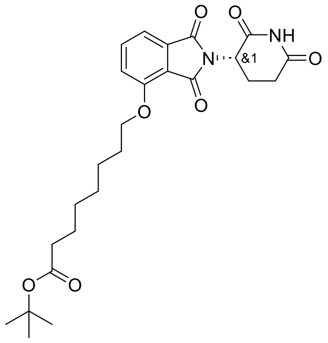

*Rac*-2-(2,6-dioxopiperidin-3-yl)-4-hydroxyisoindoline-1,3-dione (438 mg, 1.60 mmol) and *tert*-butyl 8-bromooctanoate (1.1 eq.) were used to synthesise **4k** according to general procedure F. The crude was purified by RPFCC (30 g, 40-80% B in A over 8 CV) to afford **4k** as a light green foam (577 mg, 77%). $^{1}$**H NMR** (500 MHz, Chloroform-*d*) δ 1.31–1.42 (m, 4H), 1.44 (s, 9H), 1.51 (p, *J* = 7.3 Hz, 2H), 1.55–1.62 (m, 2H), 1.84–1.91 (m, 2H), 2.10–2.15 (m, 1H), 2.21 (t, *J* = 7.5 Hz, 2H), 2.68–2.92 (m, 3H), 4.17 (td, *J* = 6.6, 1.2 Hz, 2H), 4.95 (dd, *J* = 12.3, 5.4 Hz, 1H), 7.21 (d, *J* = 8.5 Hz, 1H), 7.45 (d, *J* = 7.4 Hz, 1H), 7.67 (dd, *J* = 8.5, 7.3 Hz, 1H), 7.99 (s, 1H). **MS** (m/z): [M - H]$^-$ calcd. for $C_{25}H_{31}N_2O_7$, 471.2; found 471.3.

#### ***Rac-tert*-butyl-2-((2-(2,6-dioxopiperidin-3-yl)-1,3-dioxoisoindolin-5-yl)oxy)acetate (5g)**

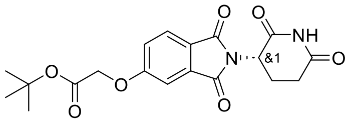

*Rac*-2-(2,6-dioxopiperidin-3-yl)-5-hydroxyisoindoline-1,3-dione (639 mg, 2.33 mmol) and *tert*-butyl 2-bromoacetate 1.1 eq.) were used to synthesise **5g** according to general procedure F. The crude was purified by RPFCC (30 g, 40-65% B in A over 7 CV) to afford **5g** as a white solid (802 mg, 89%). $^{1}$**H NMR** (500 MHz, Chloroform-*d*) δ 1.50 (s, 9H), 2.15 (ddd, *J* = 10.3, 5.2, 3.5 Hz, 1H), 2.70–2.94 (m, 3H), 4.65 (s, 2H), 4.96 (dd, *J* = 12.4, 5.4 Hz, 1H), 7.24 (dd, *J* = 8.3, 2.4 Hz, 1H), 7.30 (d, *J* = 2.3 Hz, 1H), 7.81 (d, *J* = 8.3 Hz, 1H), 7.96 (s, 1H). **MS** (m/z): [M - H]$^-$ calcd. for $C_{19}H_{19}N_2O_7$, 387.1; found 387.2.

#### ***Rac-tert*-butyl-5-((2-(2,6-dioxopiperidin-3-yl)-1,3-dioxoisoindolin-5-yl)oxy)pentanoate (5h)**

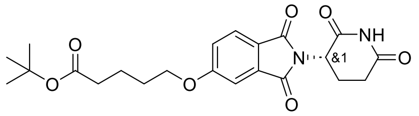

*Rac*-2-(2,6-dioxopiperidin-3-yl)-5-hydroxyisoindoline-1,3-dione (515 mg, 1.88 mmol) and *tert*-butyl 5-bromopentanoate (1.1 eq.) were used to synthesise **5h** according to general procedure F. The crude was purified by RPFCC (30 g, 45-70% B in A over 6 CV) to afford **5h** as a white foam (620 mg, 77%). $^{1}$**H NMR** (500 MHz, Chloroform-*d*) δ 1.45 (s, 9H), 1.78 (dt, *J* = 14.5, 7.0 Hz, 2H), 1.83–1.91 (m, 2H), 2.11–2.18 (m, 1H), 2.31 (t, *J* = 7.2 Hz, 2H), 2.70–2.94 (m, 3H), 4.09 (t, *J* = 6.1 Hz, 2H), 4.96 (dd, *J* = 12.4, 5.3 Hz, 1H), 7.18 (dd, *J* = 8.2, 2.3 Hz, 1H), 7.33 (d, *J* = 2.3 Hz, 1H), 7.77 (d, *J* = 8.2 Hz, 1H), 8.01 (s, 1H). **MS** (m/z): [M - H]$^-$ calcd. for $C_{22}H_{25}N_2O_7$, 429.2; found 429.1.

#### ***Rac-tert*-butyl-7-((2-(2,6-dioxopiperidin-3-yl)-1,3-dioxoisoindolin-5-yl)oxy)heptanoate (5i)**

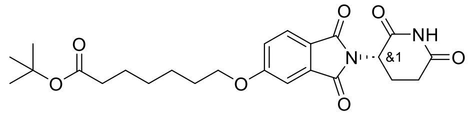

*Rac*-2-(2,6-Dioxopiperidin-3-yl)-5-hydroxyisoindoline-1,3-dione (461 mg, 1.68 mmol) and *tert*-butyl 7-bromoheptanoate (1.1 eq.) were used to synthesise **5i** according to general procedure **F**. The crude was purified by RPFCC (30 g, 40-70% B in A over 6 CV) to afford **5i** as a light green gum/film (584 mg, 76%). $^{1}$**H NMR** (500 MHz, Chloroform-*d*) δ 1.39 (p, *J* = 7.0 Hz, 2H), 1.44 (s, 9H), 1.46–1.53 (m, 2H), 1.62 (p, *J* = 7.4 Hz, 2H), 1.81–1.89 (m, 2H), 2.12–2.18 (m, 1H), 2.23 (t, *J* = 7.4 Hz, 2H), 2.70–2.94 (m, 3H), 4.07 (t, *J* = 6.4 Hz, 2H), 4.96 (dd, *J* = 12.4, 5.3 Hz, 1H), 7.18 (dd, *J* = 8.3, 2.3 Hz, 1H), 7.33 (d, *J* = 2.2 Hz, 1H), 7.77 (d, *J* = 8.3 Hz, 1H), 7.99 (s, 1H). *Tert*-butyl ester cleaved under LCMS conditions (pH =3) [M + H]$^+$ calculated for acid. **MS** (m/z): [M + H]$^+$ calcd. for $C_{20}H_{23}N_2O_7$, 403.2; found 403.4

#### ***Rac*-tert-butyl-8-((2-(2,6-dioxopiperidin-3-yl)-1,3-dioxoisoindolin-5-yl)oxy)octanoate (5j)**

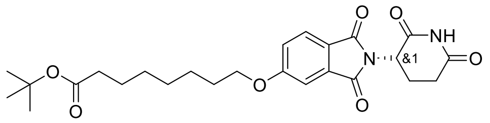

*Rac*-2-(2,6-dioxopiperidin-3-yl)-5-hydroxyisoindoline-1,3-dione (438 mg, 1.60 mmol) and *tert*-butyl 8-bromooctanoate (1.1 eq.) were used to synthesise **5j** according to general procedure F. The crude was purified by RPFCC (30 g, 50-80% B in A over 7 CV) to afford **5j** as a colourless sticky film (534 mg, 71%). **$^1$H NMR** (500 MHz, Chloroform-*d*) δ 1.37 (dtd, *J* = 12.0, 5.8, 2.5 Hz, 4H), 1.44 (s, 9H), 1.48 (q, *J* = 7.1 Hz, 2H), 1.57–1.64 (m, 2H), 1.83 (p, *J* = 6.7 Hz, 2H), 2.12–2.18 (m, 1H), 2.22 (t, *J* = 7.5 Hz, 2H), 2.70–2.95 (m, 3H), 4.07 (t, *J* = 6.5 Hz, 2H), 4.96 (dd, *J* = 12.4, 5.4 Hz, 1H), 7.18 (dd, *J* = 8.3, 2.3 Hz, 1H), 7.33 (d, *J* = 2.3 Hz, 1H), 7.77 (d, *J* = 8.3 Hz, 1H), 7.96 (s, 1H). **MS** (m/z): $[M - H]^-$ calcd. for $C_{25}H_{31}N_2O_7$, 471.2; found 471.3.

#### **General procedure G**

The corresponding *tert*-butyl ester (1.0 eq.) was dissolved in dioxane (2 mL) to which hydrochloric acid (dioxane, 4 M, 10 eq.) was added and stirred at 100 °C until complete deprotection monitored by LCMS. The reaction mixture was concentrated to afford products **6a**-**j**, **7a**-**i**. In some examples the yields reported are above 100%, probably due to HCl salt residue in product. Crude product was used in the next step without quantifying HCl salt amount.

#### ***Rac*-1-(2-(2,6-dioxopiperidin-3-yl)-1,3-dioxoisoindolin-4-yl)piperidine-4-carboxylic acid (6a)**

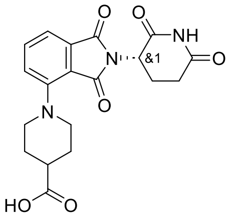

**4a** (365 mg, 0.83 mmol) was used to synthesise **6a** according to general procedure G. The reaction was stirred for 3 h. **6a** was obtained as a beige solid (421 mg, 121%). **$^1$H NMR** (500 MHz, DMSO-*d$_6$*) δ 1.76 (qd, *J* = 10.7, 7.7 Hz, 2H), 1.94 (d, *J* = 10.2 Hz, 2H), 1.99–2.06 (m, 1H), 2.45 (q, *J* = 3.5 Hz, 1H), 2.52–2.62 (m, 2H), 2.88 (ddd, *J* = 16.6, 13.8, 5.3 Hz, 1H), 2.92–3.01 (m, 2H), 3.63 (d, *J* = 12.7 Hz, 2H), 5.09 (dd, *J* = 12.8, 5.4 Hz, 1H), 7.34 (dd, *J* = 7.8, 3.9 Hz, 2H), 7.66–7.71 (m, 1H), 11.09 (s, 1H). **MS** (m/z): $[M + H]^+$ calcd. for $C_{19}H_{20}N_3O_6$, 386.1; found 386.1.

#### ***Rac*-2-(1-(2-(2,6-dioxopiperidin-3-yl)-1,3-dioxoisoindolin-4-yl)piperidin-4-yl)acetic acid (6b)**

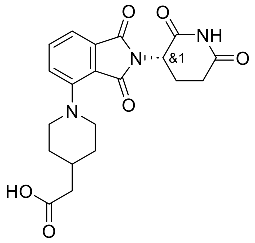

**4b** (210 mg, 0.46 mmol) was used to synthesise **6b** according to general procedure G. The reaction was stirred for 1 h. **6b** was obtained as a yellow solid (294 mg, 102%). **$^1$H NMR** (500 MHz, DMSO-*d$_6$*) δ 1.38–1.44 (m, 2H), 1.74–1.81 (m, 2H), 1.85 (ddd, *J* = 11.3, 7.7, 4.0 Hz, 1H), 1.98–2.04 (m, 1H), 2.21 (d, *J* = 7.0 Hz, 2H), 2.52–2.62 (m, 2H), 2.87 (t, *J* = 12.9 Hz, 3H), 3.67 (d, *J* = 11.3 Hz, 2H), 5.08 (dd, *J* = 12.8, 5.5 Hz, 1H), 7.32 (t, *J* = 7.1 Hz, 2H), 7.65–7.69 (m, 1H), 11.09 (s, 1H). **MS** (m/z): $[M + H]^+$ calcd. for $C_{20}H_{22}N_3O_6$, 400.3; found 400.2.

**_Rac_-3-(1-(2-(2,6-dioxopiperidin-3-yl)-1,3-dioxoisoindolin-4-yl)piperidin-4-yl)propanoic acid (6c)**

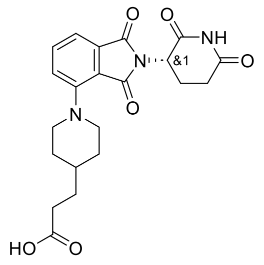

**4c** (442 mg, 0.94 mmol) was used to synthesise **6c** according to general procedure G. The reaction was stirred for 1 h. **6c** was obtained as a pale-yellow solid (435 mg, 103%). **$^1$H NMR** (500 MHz, DMSO-*d$_6$*) δ 1.32 (q, *J* = 10.2 Hz, 2H), 1.38–1.47 (m, 1H), 1.51 (q, *J* = 7.4 Hz, 2H), 1.76 (d, *J* = 12.8 Hz, 2H), 1.98–2.05 (m, 1H), 2.27 (t, *J* = 7.6 Hz, 2H), 2.51–2.62 (m, 2H), 2.79–2.92 (m, 3H), 3.65–3.72 (m, 2H), 5.09 (dd, *J* = 12.7, 5.5 Hz, 1H), 7.32 (t, *J* = 7.1 Hz, 2H), 7.65–7.70 (m, 1H), 11.09 (s, 1H). **MS** (m/z): $[M + H]^+$ calcd. for $C_{21}H_{24}N_3O_6$, 414.3; found 414.2.

**_Rac_-2-(4-(2-(2,6-dioxopiperidin-3-yl)-1,3-dioxoisoindolin-4-yl)piperazin-1-yl)acetic acid (6d)**

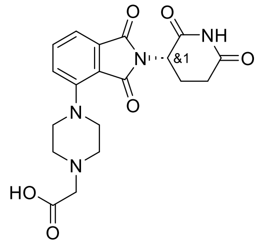

**4e** (260 mg, 0.57 mmol) was used to synthesise **6d** according to general procedure G. The reaction was stirred for 2 h. **6d** was obtained as a yellow powder (286 mg, 106%). **$^1$H NMR** (500 MHz, DMSO-*d$_6$*) δ 2.04 (dt, *J* = 7.3, 2.5 Hz, 1H), 2.51–2.63 (m, 2H), 2.88 (ddd, *J* = 17.0, 13.9, 5.4 Hz, 1H), 3.29–3.50 (m, 8H), 4.17 (s, 2H), 5.11 (dd, *J* = 12.8, 5.5 Hz, 1H), 7.41 (d, *J* = 8.5 Hz, 1H), 7.45 (d, *J* = 7.2 Hz, 1H), 7.74–7.79 (m, 1H), 11.11 (s, 1H). **MS** (m/z): $[M + H]^+$ calcd. for $C_{19}H_{21}N_4O_6$, 401.2; found 401.1.

**_Rac_-3-(4-(2-(2,6-dioxopiperidin-3-yl)-1,3-dioxoisoindolin-4-yl)piperazin-1-yl)propanoic acid (6e)**

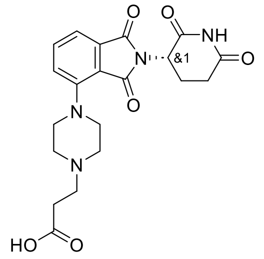

**4f** (399 mg, 0.85 mmol) was used to synthesise **6e** according to general procedure G. The reaction was stirred for 1h. **6e** was obtained as a yellow powder (386 mg, 93%). **$^1$H NMR** (500 MHz, DMSO-*d$_6$*) δ 2.03 (dtd, *J* = 13.1, 5.4, 2.4 Hz, 1H), 2.51–2.64 (m, 2H), 2.90 (t, *J* = 7.8 Hz, 2H), 3.16 (s, 4H), 3.39 (d, *J* = 7.4 Hz, 2H), 3.48–3.65 (m, 3H), 3.68–3.95 (m, 2H), 5.11 (dd, *J* = 12.8, 5.5 Hz, 1H), 7.42 (d, *J* = 8.4 Hz, 1H), 7.46 (d, *J* = 7.2 Hz, 1H), 7.76 (dd, *J* = 8.4, 7.2 Hz, 1H), 11.11 (s, 1H). **MS** (m/z): $[M + H]^+$ calcd. for $C_{20}H_{23}N_4O_6$, 415.2; found 415.3.

**_Rac_-1-(2-(2,6-dioxopiperidin-3-yl)-1,3-dioxoisoindolin-5-yl)piperidine-4-carboxylic acid (7a)**

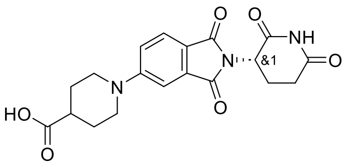

**5a** (370 mg, 0.81 mmol) was used to synthesise **7a** according to general procedure G. The reaction was stirred for 1 h. **7a** was obtained as a yellow solid (351 mg, 92%). **$^1$H NMR** (500 MHz, DMSO-*d$_6$*) δ 1.54–1.65 (m, 2H), 1.89 (d, *J* = 9.5 Hz, 2H), 1.97–2.04 (m, 1H), 2.52–2.61 (m, 3H), 2.88 (ddd, *J* = 16.5, 13.7, 5.4 Hz, 1H), 3.08 (t, *J* = 11.0 Hz, 2H), 3.97 (d, *J* = 13.3 Hz, 2H), 5.06 (dd, *J* = 12.8, 5.5 Hz, 1H), 7.25 (d, *J* = 11.1 Hz, 1H), 7.34 (s, 1H), 7.66 (d, *J* = 8.6 Hz, 1H), 11.08 (s, 1H). COOH not observed. **MS** (m/z): $[M + H]^+$ calcd. for $C_{19}H_{20}N_3O_6$, 386.1; found 386.2.

***Rac*-2-(1-(2-(2,6-dioxopiperidin-3-yl)-1,3-dioxoisoindolin-5-yl)piperidin-4-yl)acetic acid (7b)**

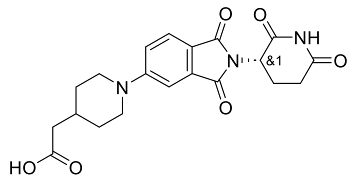


**5b** (325 mg, 0.71 mmol) was used to synthesise **7b** according to general procedure G. The reaction was stirred for 1 h. The crude was purified by preparative HPLC (acidic, 10-55% B in A over 25 min) to afford **7b** as a yellow solid (71 mg, 25%). **$^1$H NMR** (500 MHz, DMSO-*d$_6$)* δ 1.24 (td, *J* = 12.4, 3.8 Hz, 2H), 1.74 (d, *J* = 3.7 Hz, 2H), 1.95 (tt, *J* = 7.4, 3.9 Hz, 1H), 1.99–2.05 (m, 1H), 2.18 (d, *J* = 7.0 Hz, 2H), 2.52–2.62 (m, 2H), 2.84–2.90 (m, 1H), 2.96 (t, *J* = 11.4 Hz, 2H), 4.04 (d, *J* = 13.3 Hz, 2H), 5.06 (dd, *J* = 12.8, 5.4 Hz, 1H), 7.23 (d, *J* = 8.7 Hz, 1H), 7.30 (s, 1H), 7.64 (d, *J* = 8.5 Hz, 1H), 11.08 (s, 1H). **MS** (m/z): [M + H]$^+$ calcd. for $C_{20}H_{22}N_3O_6$, 400.2; found 400.3.

***Rac*-3-(1-(2-(2,6-dioxopiperidin-3-yl)-1,3-dioxoisoindolin-5-yl)piperidin-4-yl)propanoic acid (7c)**

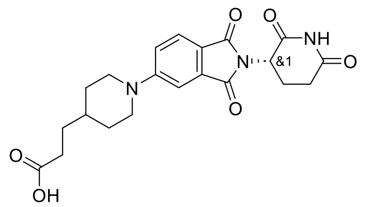


**5c** (445 mg, 0.95 mmol) was used to synthesise **7c** according to general procedure G. The reaction was stirred for 3 h. **7c** was obtained as a beige solid (462 mg, 108%). **$^1$H NMR** (500 MHz, DMSO-*d$_6$)* δ 1.16 (qd, *J* = 12.4, 3.9 Hz, 2H), 1.46 (q, *J* = 7.4 Hz, 2H), 1.54 (ddt, *J* = 10.4, 6.9, 3.8 Hz, 1H), 1.74 (d, *J* = 10.2 Hz, 2H), 1.98–2.03 (m, 1H), 2.25 (t, *J* = 7.6 Hz, 2H), 2.52–2.62 (m, 2H), 2.82–2.98 (m, 3H), 4.03 (d, *J* = 13.3 Hz, 2H), 5.06 (dd, *J* = 12.8, 5.4 Hz, 1H), 7.25 (dd, *J* = 8.6, 2.5 Hz, 1H), 7.32 (d, *J* = 2.5 Hz, 1H), 7.65 (d, *J* = 8.6 Hz, 1H), 11.08 (s, 1H). **MS** (m/z): [M + H]$^+$ calcd. for $C_{21}H_{24}N_3O_6$, 414.2; found 414.2.

***Rac*-2-(4-(2-(2,6-dioxopiperidin-3-yl)-1,3-dioxoisoindolin-5-yl)piperazin-1-yl)acetic acid (7d)**

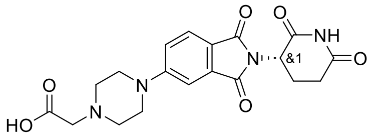


**5e** (280 mg, 0.61 mmol) was used to synthesise **7d** according to general procedure G. The reaction was stirred for 1 h. **7d** was obtained as a yellow powder (313 mg, 108%). **$^1$H NMR** (500 MHz, DMSO-*d$_6$)* δ 1.98–2.06 (m, 1H), 2.53–2.64 (m, 2H), 2.89 (ddd, *J* = 16.3, 13.7, 5.4 Hz, 1H), 3.23–3.62 (m, 8H), 4.21 (s, 2H), 5.09 (dd, *J* = 12.8, 5.4 Hz, 1H), 7.34 (d, *J* = 8.6 Hz, 1H), 7.47 (s, 1H), 7.76 (d, *J* = 8.6 Hz, 1H), 11.10 (s, 1H). **MS** (m/z): [M + H]$^+$ calcd. for $C_{19}H_{21}N_4O_6$, 401.2; found 401.2.

***Rac*-3-(4-(2-(2,6-dioxopiperidin-3-yl)-1,3-dioxoisoindolin-5-yl)piperazin-1-yl)propanoic acid (7e)**

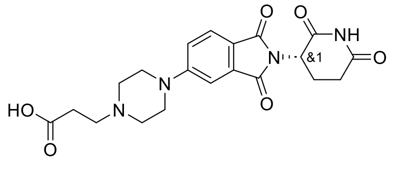


**5f** (431 mg, 0.92 mmol) was used to synthesise **7e** according to general procedure G. The reaction was stirred overnight. **7e** was obtained as a beige powder (430 mg, 96%). **$^1$H NMR** (500 MHz, DMSO-*d$_6$)* δ 2.03 (ddd, *J* = 12.9, 5.8, 3.4 Hz, 1H), 2.51–2.63 (m, 2H), 2.88 (ddd, *J* = 12.6, 7.8, 4.5 Hz, 3H), 3.15 (t, *J* = 10.6 Hz, 2H), 3.30–3.43 (m, 4H), 3.56 (d, *J* = 12.8 Hz, 2H), 4.22 (d, *J* = 14.0 Hz, 2H), 5.09 (dd, *J* = 12.9, 5.4 Hz, 1H), 7.35 (dd, *J* = 8.6, 2.3 Hz, 1H), 7.49 (d, *J* = 2.5 Hz, 1H), 7.75 (d, *J* = 8.5 Hz, 1H), 11.10 (s, 1H). **MS** (m/z): [M + H]$^+$ calcd. for $C_{20}H_{23}N_4O_6$, 415.2; found 415.4.

**_Rac_-2-((2-(2,6-dioxopiperidin-3-yl)-1,3-dioxoisoindolin-4-yl)oxy)acetic acid (6f)**

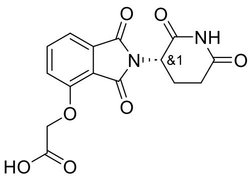

**4g** (366 mg, 0.94 mmol) was used to synthesise **6f** according to general procedure G. The reaction was stirred for 1 h. **6f** was obtained as a white solid (310 mg, 99%). **$^1$H NMR** (500 MHz, DMSO-*d$_6$*) δ 2.04 (ddd, *J* = 7.5, 5.9, 3.0 Hz, 1H), 2.51–2.63 (m, 2H), 2.89 (ddd, *J* = 16.9, 13.9, 5.4 Hz, 1H), 4.99 (s, 2H), 5.10 (dd, *J* = 12.8, 5.5 Hz, 1H), 7.39 (d, *J* = 8.6 Hz, 1H), 7.48 (d, *J* = 7.2 Hz, 1H), 7.77–7.82 (m, 1H), 11.12 (s, 1H), 13.25 (s, 1H). **MS** (m/z): $[M + H]^+$ calcd. for $C_{15}H_{13}N_2O_7$, 333.1; found 333.1.

**_Rac_-5-((2-(2,6-dioxopiperidin-3-yl)-1,3-dioxoisoindolin-4-yl)oxy)pentanoic acid (6g)**

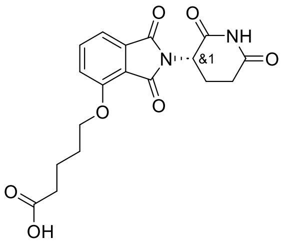

**4h** (657 mg, 1.53 mmol) was used to synthesise **6g** according to general procedure G. The reaction was stirred for 30 min. **6g** was obtained as a white solid (691 mg, 121%). **$^1$H NMR** (500 MHz, DMSO-*d$_6$*) δ 1.70 (q, *J* = 7.4 Hz, 2H), 1.74–1.82 (m, 2H), 1.99–2.05 (m, 1H), 2.31 (t, *J* = 7.3 Hz, 2H), 2.51–2.62 (m, 2H), 2.88 (ddd, *J* = 16.9, 13.9, 5.4 Hz, 1H), 4.21 (t, *J* = 6.2 Hz, 2H), 5.08 (dd, *J* = 12.8, 5.4 Hz, 1H), 7.44 (d, *J* = 7.2 Hz, 1H), 7.52 (d, *J* = 8.6 Hz, 1H), 7.81 (dd, *J* = 8.5, 7.3 Hz, 1H), 11.11 (s, 1H), 12.03 (s, 1H). **MS** (m/z): $[M + H]^+$ calcd. for $C_{18}H_{19}N_2O_7$, 375.1; found 375.1.

**_Rac_-6-((2-(2,6-dioxopiperidin-3-yl)-1,3-dioxoisoindolin-4-yl)oxy)hexanoic acid (6h)**

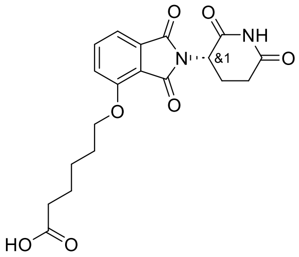

**4i** (583 mg, 1.31 mmol) was used to synthesise **6h** according to general procedure G. The reaction was stirred for 30 min. **6h** was obtained as a colourless sticky gum (635 mg, 125%). **$^1$H NMR** (500 MHz, DMSO-*d$_6$*) δ 1.42–1.50 (m, 2H), 1.58 (p, *J* = 7.4 Hz, 2H), 1.76 (p, *J* = 6.8 Hz, 2H), 1.99–2.06 (m, 1H), 2.23 (t, *J* = 7.4 Hz, 2H), 2.51–2.62 (m, 2H), 2.88 (ddd, *J* = 17.0, 13.9, 5.4 Hz, 1H), 4.20 (t, *J* = 6.4 Hz, 2H), 5.08 (dd, *J* = 12.8, 5.4 Hz, 1H), 7.44 (d, *J* = 7.2 Hz, 1H), 7.51 (d, *J* = 8.5 Hz, 1H), 7.81 (dd, *J* = 8.5, 7.2 Hz, 1H), 11.11 (s, 1H), 12.01 (s, 1H). **MS** (m/z): $[M + H]^+$ calcd. for $C_{19}H_{21}N_2O_7$, 389.1; found 389.1.

**_Rac_-7-((2-(2,6-dioxopiperidin-3-yl)-1,3-dioxoisoindolin-4-yl)oxy)heptanoic acid (6i)**

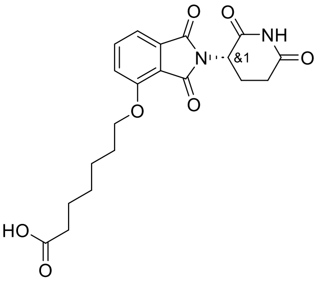

**4j** (573 mg, 1.25 mmol) was used to synthesise **6i** according to general procedure G. The reaction was stirred for 30 min. **6i** was obtained as a white gum (633 mg, 126%). **$^1$H NMR** (500 MHz, DMSO-*d$_6$*) δ 1.35 (q, *J* = 8.8 Hz, 2H), 1.42–1.55 (m, 4H), 1.75 (p, *J* = 6.6 Hz, 2H), 1.99–2.05 (m, 1H), 2.21 (t, *J* = 7.3 Hz, 2H), 2.51–2.62 (m, 2H), 2.88 (ddd, *J* = 16.9, 13.9, 5.5 Hz, 1H), 4.20 (t, *J* = 6.4 Hz, 2H), 5.08 (dd, *J* = 12.8, 5.4 Hz, 1H), 7.44 (d, *J* = 7.2 Hz, 1H), 7.51 (d, *J* = 8.6 Hz, 1H), 7.80 (dd, *J* = 8.5, 7.2 Hz, 1H), 11.11 (s, 1H), 11.98 (s, 1H). **MS** (m/z): $[M + H]^+$ calcd. for $C_{20}H_{23}N_2O_7$, 403.1; found 403.2.

***Rac*-8-((2-(2,6-dioxopiperidin-3-yl)-1,3-dioxoisoindolin-4-yl)oxy)octanoic acid (6j)**

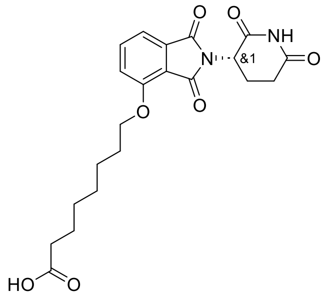

**4k** (573 mg, 1.21 mmol) was used to synthesise **6j** according to general procedure G. The reaction was stirred for 30 min. **6j** was obtained as a white gum (692 mg, 137%). **$^1$H NMR** (500 MHz, DMSO-*d$_6$*) δ 1.24–1.38 (m, 4H), 1.41–1.55 (m, 4H), 1.75 (p, *J* = 6.5 Hz, 2H), 2.03 (ddd, *J* = 9.9, 6.3, 2.3 Hz, 1H), 2.20 (t, *J* = 7.3 Hz, 2H), 2.51–2.62 (m, 2H), 2.88 (ddd, *J* = 16.9, 13.9, 5.4 Hz, 1H), 4.20 (t, *J* = 6.4 Hz, 2H), 5.08 (dd, *J* = 12.8, 5.4 Hz, 1H), 7.44 (d, *J* = 7.2 Hz, 1H), 7.51 (d, *J* = 8.5 Hz, 1H), 7.80 (dd, *J* = 8.5, 7.3 Hz, 1H), 11.11 (s, 1H), 11.96 (s, 1H). **MS** (m/z): [M + H]$^+$ calcd. for $C_{21}H_{25}N_2O_7$, 417.2; found 417.4.

***Rac*-2-((2-(2,6-dioxopiperidin-3-yl)-1,3-dioxoisoindolin-5-yl)oxy)acetic acid (7f)**

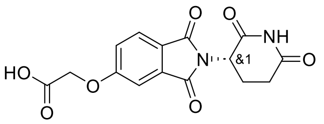

**5g** (799 mg, 2.06 mmol) was used to synthesise **7f** according to general procedure G. The reaction was stirred for 30 min. **7f** was obtained as a white solid (869 mg, 127%). **$^1$H NMR** (500 MHz, DMSO-*d$_6$*) δ 2.01–2.08 (m, 1H), 2.52–2.63 (m, 2H), 2.89 (ddd, *J* = 16.9, 13.8, 5.4 Hz, 1H), 4.96 (s, 2H), 5.12 (dd, *J* = 12.9, 5.4 Hz, 1H), 7.36 (dd, *J* = 8.3, 2.3 Hz, 1H), 7.42 (d, *J* = 2.3 Hz, 1H), 7.85 (d, *J* = 8.3 Hz, 1H), 11.12 (s, 1H), 13.21 (s, 1H). **MS** (m/z): [M + H]$^+$ calcd. for $C_{15}H_{13}N_2O_7$, 333.1; found 333.1.

***Rac*-5-((2-(2,6-dioxopiperidin-3-yl)-1,3-dioxoisoindolin-5-yl)oxy)pentanoic acid (7g)**

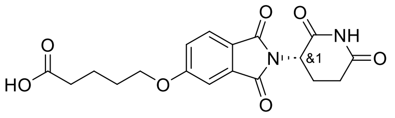

**5h** (618, 1.44 mmol) was used to synthesise **7g** according to general procedure G. The reaction was stirred for 30 min. **7g** was obtained as a white gum (653 mg, 122%). **$^1$H NMR** (500 MHz, DMSO-*d$_6$*) δ 1.66 (p, *J* = 7.6 Hz, 2H), 1.74–1.81 (m, 2H), 2.01–2.07 (m, 1H), 2.30 (t, *J* = 7.3 Hz, 2H), 2.51–2.62 (m, 2H), 2.89 (ddd, *J* = 16.8, 13.8, 5.4 Hz, 1H), 4.18 (t, *J* = 6.3 Hz, 2H), 5.12 (dd, *J* = 12.8, 5.4 Hz, 1H), 7.35 (dd, *J* = 8.3, 2.3 Hz, 1H), 7.43 (d, *J* = 2.3 Hz, 1H), 7.83 (d, *J* = 8.3 Hz, 1H), 11.12 (s, 1H), 12.06 (s, 1H). **MS** (m/z): [M + H]$^+$ calcd. for $C_{18}H_{19}N_2O_7$, 375.1 found 375.1.

***Rac*-7-((2-(2,6-dioxopiperidin-3-yl)-1,3-dioxoisoindolin-5-yl)oxy)heptanoic acid (7h)**

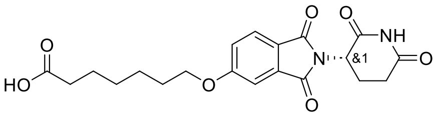

**5i** (580 mg, 1.26 mmol) was used to synthesise **7h** according to general procedure G. The reaction was stirred for 30 min. **7h** was obtained as a white gum (713 mg, 140%). **$^1$H NMR** (500 MHz, DMSO-*d$_6$*) δ 1.34 (q, *J* = 8.1 Hz, 2H), 1.43 (p, *J* = 7.1 Hz, 2H), 1.52 (p, *J* = 7.4 Hz, 2H), 1.71–1.78 (m, 2H), 2.01–2.07 (m, 1H), 2.21 (t, *J* = 7.3 Hz, 2H), 2.51–2.62 (m, 2H), 2.89 (ddd, *J* = 16.8, 13.8, 5.4 Hz, 1H), 4.16 (t, *J* = 6.5 Hz, 2H), 5.12 (dd, *J* = 12.8, 5.4 Hz, 1H), 7.34 (dd, *J* = 8.3, 2.3 Hz, 1H), 7.42 (d, *J* = 2.3 Hz, 1H), 7.83 (d, *J* = 8.3 Hz, 1H), 11.12 (s, 1H), 11.99 (s, 1H). **MS** (m/z): [M + H]$^+$ calcd. for $C_{20}H_{23}N_2O_7$, 403.2; found 403.2.

***Rac*-8-((2-(2,6-dioxopiperidin-3-yl)-1,3-dioxoisoindolin-5-yl)oxy)octanoic acid (7i)**

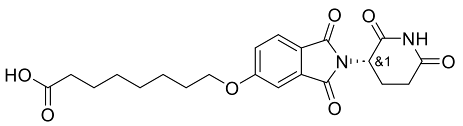

**5j** (532 mg, 1.13 mmol) was used to synthesise **7i** according to general procedure G. The reaction was stirred for 30 min. **7i** was obtained as a white sticky solid (630 mg, 134%). **$^1$H NMR** (500 MHz, DMSO-*d$_6$*) δ 1.26–1.37 (m, 4H), 1.41 (q, *J* = 7.4 Hz, 2H), 1.50 (p, *J* = 7.4 Hz, 2H), 1.74 (p, *J* = 6.6 Hz, 2H), 2.01–2.08 (m, 1H), 2.20 (t, *J* = 7.4 Hz, 2H), 2.51–2.62 (m, 2H), 2.89 (ddd, *J* = 16.9, 13.8, 5.4 Hz, 1H), 4.16 (t, *J* = 6.5 Hz, 2H), 5.12 (dd, *J* = 12.8, 5.4 Hz,

1H), 7.34 (dd, *J* = 8.3, 2.3 Hz, 1H), 7.42 (d, *J* = 2.3 Hz, 1H), 7.82 (d, *J* = 8.3 Hz, 1H), 11.12 (s, 1H), 11.98 (s, 1H). **MS** (m/z): $[M + H]^+$ calcd. for $C_{21}H_{25}N_2O_7$, 417.2; found 417.2.

#### General procedure H

SNAP2 ligand (1.0 eq.) and the corresponding acid (1.0 eq.) were dissolved in DCM (0.5 mL) followed by addition of T3P (2 eq.) and DIPEA (4-10 eq.). The reaction mixture was stirred at room temperature until no further progression monitored by LCMS. The crude was purified by RPFCC to afford the corresponding PROTAC.

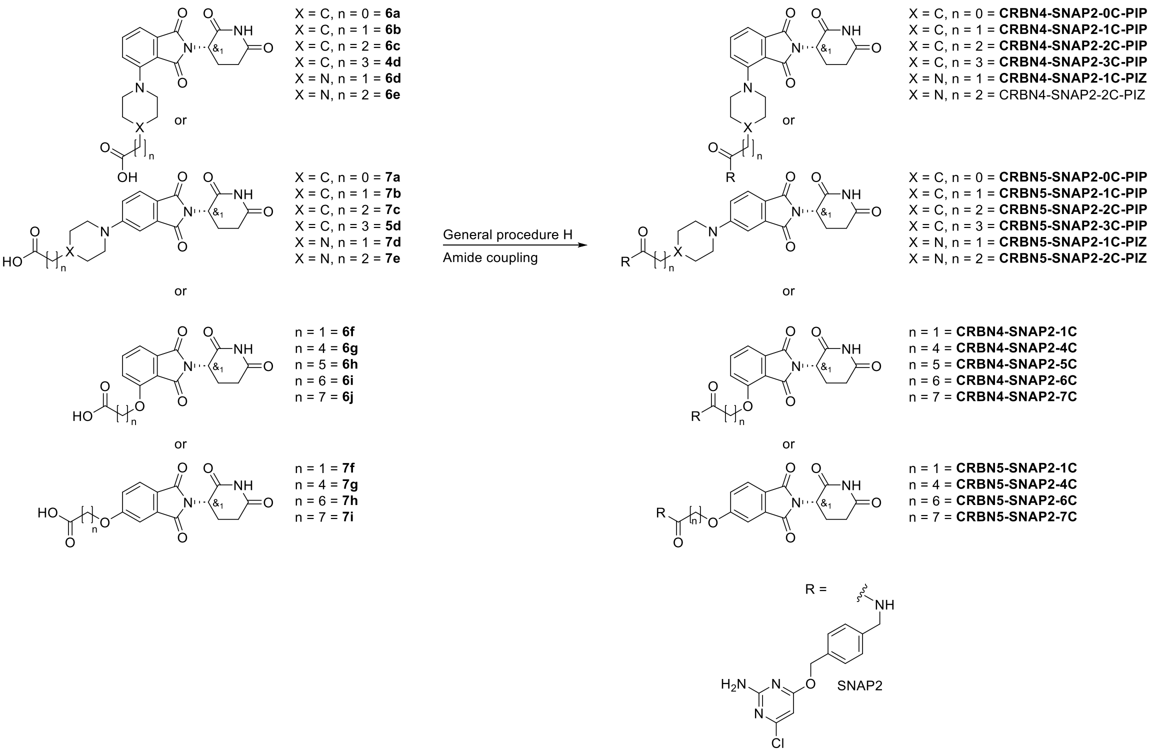


**Scheme S3**. Synthesis overview of CRBN-SNAP2-PROTACs

#### ***Rac-N*-(4-(((2-amino-6-chloropyrimidin-4-yl)oxy)methyl)benzyl)-1-(2-(2,6-dioxopiperidin-3-yl)-1,3-dioxoisoindolin-4-yl)piperidine-4-carboxamide (CRBN4-SNAP2-0C-PIP)**

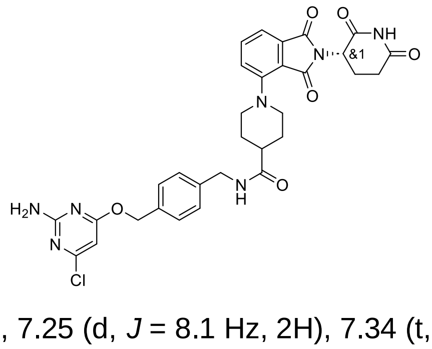

SNAP2 ligand (5 mg, 0.02 mmol), **6a** (8 mg, 1.0 eq.) and DIPEA (4 eq.) were used to synthesise **CRBN4-SNAP2-0C-PIP** according to general procedure H. The reaction was stirred for 1 h. The crude was purified by RPFCC (6 g, 45-60% B in A over 6 CV) to afford **CRBN4-SNAP2-0C-PIP** as a yellow fluffy solid (4.7 mg, 39%). **$^1$H NMR** (500 MHz, DMSO-*d$_6$*) δ 1.82 (q, *J* = 4.6 Hz, 4H), 1.99–2.06 (m, 1H), 2.34–2.43 (m, 1H), 2.52–2.62 (m, 2H), 2.83–2.94 (m, 3H), 3.72 (d, *J* = 11.7 Hz, 2H), 4.28 (d, *J* = 5.8 Hz, 2H), 5.10 (dd, *J* = 12.7, 5.4 Hz, 1H), 5.29 (s, 2H), 6.13 (s, 1H), 7.11 (s, 2H), 7.25 (d, *J* = 8.1 Hz, 2H), 7.34 (t, *J* = 8.4 Hz, 2H), 7.39 (d, *J* = 8.2 Hz, 2H), 7.68 (dd, *J* = 8.4, 7.1 Hz, 1H), 8.40 (t, *J* = 6.0 Hz, 1H), 11.09 (s, 1H). **MS** (m/z): $[M + H]^+$ calcd. for $C_{31}H_{31}ClN_7O_6$, 632.202; found 632.203.

#### ***Rac-N*-(4-(((2-amino-6-chloropyrimidin-4-yl)oxy)methyl)benzyl)-2-(1-(2-(2,6-dioxopiperidin-3-yl)-1,3-dioxoisoindolin-4-yl)piperidin-4-yl)acetamide (CRBN4-SNAP2-1C-PIP)**

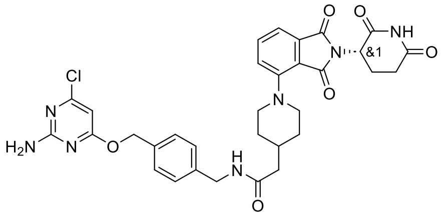


SNAP2 ligand (5 mg, 0.02 mmol), **6b** (8.2 mg, 1.0 eq.) and DIPEA (4 eq.) were used to synthesise **CRBN4-SNAP2-1C-PIP** according to general procedure H. The reaction was stirred for 1 h. The crude was purified by RPFCC (6 g, 45-60% B in A over 6 CV) to afford **CRBN4-SNAP2-1C-PIP** as a yellow fluffy solid (4.6 mg, 38%). **$^1$H NMR** (500 MHz, DMSO-*d$_6$*) δ 1.39 (q, *J* = 11.4 Hz, 2H), 1.74 (d, *J* = 12.6 Hz, 2H), 1.90 (s, 1H), 1.99–2.05 (m, 1H), 2.14 (d, *J* = 7.0 Hz, 2H), 2.55–2.65 (m, 2H), 2.86 (t, *J* = 10.8 Hz, 3H), 3.66 (d, *J* = 11.7 Hz, 2H), 4.27 (d, *J* = 5.8 Hz, 2H), 5.09 (dd, *J* = 12.8, 5.5 Hz, 1H), 5.29 (s, 2H), 6.13 (s, 1H), 7.11 (s, 2H), 7.26 (d, *J* = 7.8 Hz, 2H), 7.33 (d, *J* = 7.9 Hz, 2H), 7.39 (d, *J* = 7.7 Hz, 2H), 7.67 (t, *J* = 7.8 Hz, 1H), 8.38 (t, *J* = 5.9 Hz, 1H), 11.09 (s, 1H). **MS** (m/z): [M + H]$^+$ calcd. for $C_{32}H_{33}ClN_7O_6$, 646.218; found 646.217.

#### ***Rac-N*-(4-(((2-amino-6-chloropyrimidin-4-yl)oxy)methyl)benzyl)-3-(1-(2-(2,6-dioxopiperidin-3-yl)-1,3-dioxoisoindolin-4-yl)piperidin-4-yl)propenamide (CRBN4-SNAP2-2C-PIP)**

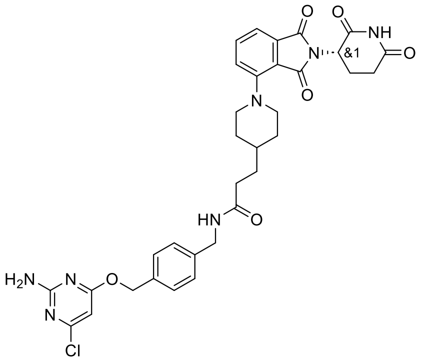


SNAP2 ligand (10 mg, 0.04 mmol), **6c** (17 mg, 1.0 eq.) and DIPEA (6 eq.) were used to synthesise **CRBN4-SNAP2-2C-PIP** according to general procedure H. The reaction was stirred for 3 h. The crude was purified by RPFCC (6 g, 50-70% B in A over 6 CV) to afford **CRBN4-SNAP2-2C-PIP** as a yellow fluffy solid (16.7 mg, 67%). **$^1$H NMR** (500 MHz, DMSO-*d$_6$*) δ 1.33 (t, *J* = 11.8 Hz, 3H), 1.53 (q, *J* = 7.1 Hz, 2H), 1.76 (d, *J* = 11.8 Hz, 2H), 1.99–2.06 (m, 1H), 2.20 (t, *J* = 7.6 Hz, 2H), 2.53–2.62 (m, 2H), 2.77–2.92 (m, 3H), 3.68 (d, *J* = 11.2 Hz, 2H), 4.26 (d, *J* = 5.9 Hz, 2H), 5.09 (dd, *J* = 12.7, 5.4 Hz, 1H), 5.28 (s, 2H), 6.10 (s, 1H), 7.10 (s, 2H), 7.25 (d, *J* = 7.9 Hz, 2H), 7.32 (dd, *J* = 7.8, 4.1 Hz, 2H), 7.39 (d, *J* = 7.9 Hz, 2H), 7.67 (dd, *J* = 8.5, 7.1 Hz, 1H), 8.35 (t, *J* = 6.0 Hz, 1H), 11.09 (s, 1H). **MS** (m/z): [M + H]$^+$ calcd. for $C_{33}H_{35}ClN_7O_6$, 660.234; found 660.231.

#### ***Rac-N*-(4-(((2-amino-6-chloropyrimidin-4-yl)oxy)methyl)benzyl)-4-(1-(2-(2,6-dioxopiperidin-3-yl)-1,3-dioxoisoindolin-4-yl)piperidin-4-yl)butanamide (CRBN4-SNAP2-3C-PIP)**

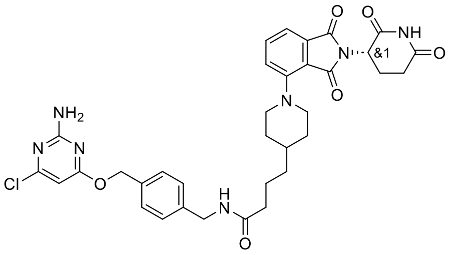


SNAP2 ligand (10 mg, 0.04 mmol) and **4d** (17.5 mg, 1.0 eq.) were used to synthesise **CRBN4-SNAP2-3C-PIP** according to general procedure H. The reaction was stirred overnight. The crude was purified by RPFCC (6 g, 50-75% B in A over 8 CV) to afford **CRBN4-SNAP2-3C-PIP** as a yellow fluffy solid (13.4 mg, 53%). **$^1$H NMR** (500 MHz, DMSO-*d$_6$*) δ 1.21–1.35 (m, 4H), 1.42 (s, 1H), 1.54–1.62 (m, 2H), 1.75 (d, *J* = 12.3 Hz, 2H), 1.99–2.05 (m, 1H), 2.14 (t, *J* = 7.3 Hz, 2H), 2.53–2.62 (m, 2H), 2.80–2.92 (m, 3H), 3.68 (d, *J* = 11.7 Hz, 2H), 4.25 (d, *J* = 5.9 Hz, 2H), 5.09 (dd, *J* = 12.7, 5.4 Hz, 1H), 5.28 (s, 2H), 6.12 (s, 1H), 7.11 (s, 2H), 7.25 (d, *J* = 7.9 Hz, 2H), 7.32 (t, *J* = 7.4 Hz, 2H), 7.38 (d, *J* = 7.9 Hz, 2H), 7.67 (dd, *J* = 8.5, 7.1 Hz, 1H), 8.32 (t, *J* = 5.9 Hz, 1H), 11.09 (s, 1H). **MS** (m/z): [M + H]$^+$ calcd. for $C_{34}H_{37}ClN_7O_6$, 674.249; found 674.251.

#### *Rac-N*-(4-(((2-amino-6-chloropyrimidin-4-yl)oxy)methyl)benzyl)-2-(4-(2-(2,6-dioxopiperidin-3-yl)-1,3-dioxoisoindolin-4-yl)piperazin-1-yl)acetamide (CRBN4-SNAP2-1C-PIZ)

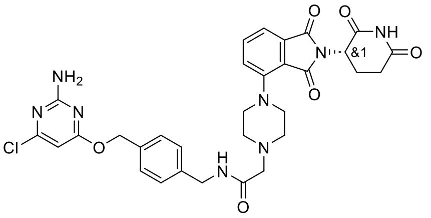

SNAP2 ligand (10 mg, 0.04 mmol) and **6d** (17.9 mg, 0.04 mmol) and DIPEA (10 eq.) were used to synthesise **CRBN4-SNAP2-1C-PIZ** according to general procedure H. The reaction was stirred overnight. The crude was purified by RPFCC (6 g, 30-45% B in A over 6 CV) to afford **CRBN4-SNAP2-1C-PIZ** as a yellow fluffy solid (5.9 mg, 24%). **$^1$H NMR** (500 MHz, DMSO-*d$_6$*) δ 1.99–2.06 (m, 1H), 2.52–2.82 (m, 6H), 2.87 (ddd, *J* = 16.8, 13.8, 5.4 Hz, 1H), 3.09 (s, 2H), 3.36 (s, 4H), 4.32 (d, *J* = 6.1 Hz, 2H), 5.09 (dd, *J* = 12.7, 5.4 Hz, 1H), 5.29 (s, 2H), 6.13 (s, 1H), 7.04–7.18 (m, 2H), 7.28 (d, *J* = 7.9 Hz, 2H), 7.34–7.41 (m, 4H), 7.71 (t, *J* = 7.8 Hz, 1H), 8.41 (s, 1H), 11.10 (s, 1H). **MS** (m/z): [M + H]$^+$ calcd. for $C_{31}H_{32}ClN_8O_6$, 647.213; found 647.212.

#### *Rac-N*-(4-(((2-amino-6-chloropyrimidin-4-yl)oxy)methyl)benzyl)-3-(4-(2-(2,6-dioxopiperidin-3-yl)-1,3-dioxoisoindolin-4-yl)piperazin-1-yl)propenamide (CRBN4-SNAP2-2C-PIZ)

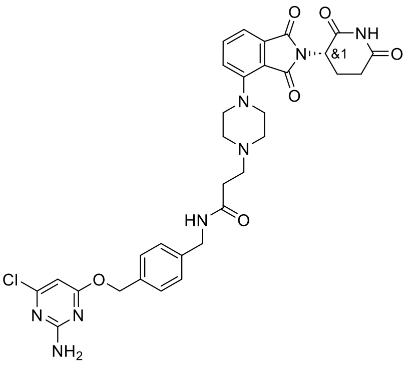

SNAP2 ligand (10 mg, 0.04 mmol), **6e** (18.4 mg, 0.04 mmol) and DIPEA (6 eq.) were used to synthesise **CRBN4-SNAP2-2C-PIZ** according to general procedure H. The reaction was stirred overnight. The crude was purified by RPFCC (6 g, 35-50% B in A over 6 CV) to afford **CRBN4-SNAP2-2C-PIZ** as a yellow fluffy solid (11.4 mg, 46%). **$^1$H NMR** (500 MHz, DMSO-*d$_6$*) δ 2.00–2.06 (m, 1H), 2.48 (s, 2H), 2.53–2.62 (m, 2H), 2.88 (ddd, *J* = 17.0, 13.8, 5.5 Hz, 1H), 3.37 (s, 10H), 4.29 (d, *J* = 5.7 Hz, 2H), 5.10 (dd, *J* = 12.8, 5.5 Hz, 1H), 5.28 (s, 2H), 6.10 (s, 1H), 7.11 (s, 2H), 7.29 (d, *J* = 7.9 Hz, 2H), 7.38 (d, *J* = 8.0 Hz, 4H), 7.72 (t, *J* = 7.7 Hz, 1H), 8.55 (s, 1H), 11.10 (s, 1H). **MS** (m/z): [M + H]$^+$ calcd. for $C_{32}H_{34}ClN_8O_6$, 661.229; found 661.230.

#### *Rac-N*-(4-(((2-amino-6-chloropyrimidin-4-yl)oxy)methyl)benzyl)-1-(2-(2,6-dioxopiperidin-3-yl)-1,3-dioxoisoindolin-5-yl)piperidine-4-carboxamide (CRBN5-SNAP2-0C-PIP)

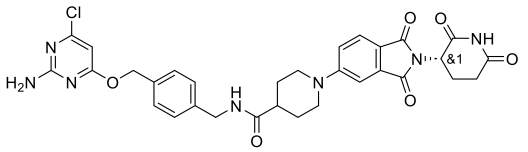

SNAP2 ligand (10 mg, 0.04 mmol), **7a** (15.9 mg, 0.04 mmol) and DIPEA (4 eq.) were used to synthesise **CRBN5-SNAP2-0C-PIP** according to general procedure H. The reaction was stirred for 1 h. The crude was purified by RPFCC (6 g, 45-60% B in A over 6 CV) to afford **CRBN5-SNAP2-0C-PIP** as a yellow fluffy solid (10.0 mg, 42%). **$^1$H NMR** (500 MHz, DMSO-*d$_6$*) δ 1.63 (qd, *J* = 12.6, 3.9 Hz, 2H), 1.80 (dd, *J* = 13.7, 3.7 Hz, 2H), 1.98–2.04 (m, 1H), 2.52–2.61 (m, 3H), 2.88 (ddd, *J* = 16.5, 13.7, 5.3 Hz, 1H), 2.97–3.05 (m, 2H), 4.08 (d, *J* = 13.2 Hz, 2H), 4.26 (d, *J* = 5.9 Hz, 2H), 5.07 (dd, *J* = 12.8, 5.4 Hz, 1H), 5.28 (s, 2H), 6.13 (s, 1H), 7.07–7.15 (m, 2H), 7.22–7.27 (m, 3H), 7.33 (d, *J* = 2.3 Hz, 1H), 7.36–7.40 (m, 2H), 7.66 (d, *J* = 8.5 Hz, 1H), 8.39 (t, *J* = 6.0 Hz, 1H), 11.08 (s, 1H). **$^{13}$C NMR** (126 MHz, DMSO-*d$_6$*) δ 173.9, 172.9, 170.3, 170.1, 167.6, 167.0, 162.8, 160.0, 154.8, 139.7, 134.6, 134.1, 128.4, 127.1, 125.0, 117.7, 117.6, 107.9, 94.4, 67.3, 48.7, 46.9, 41.7, 41.6, 31.0, 27.6, 22.2. **MS** (m/z): [M + H]$^+$ calcd. for $C_{31}H_{31}ClN_7O_6$, 632.202; found 632.203.

***Rac-N*-(4-(((2-amino-6-chloropyrimidin-4-yl)oxy)methyl)benzyl)-2-(1-(2-(2,6-dioxopiperidin-3-yl)-1,3-dioxoisoindolin-5-yl)piperidin-4-yl)acetamide (CRBN5-SNAP2-1C-PIP)**

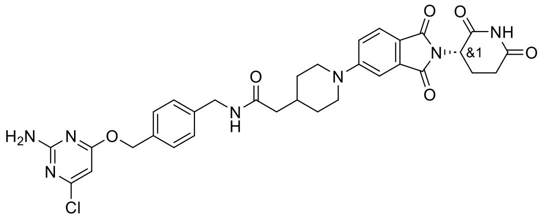

SNAP2 ligand (5 mg, 0.02 mmol), **7b** (8.2 mg, 0.02 mmol) and DIPEA (4 eq.) were used to synthesise **CRBN5-SNAP2-1C-PIP** according to general procedure H. The reaction was stirred for 1 h. The crude was purified by RPFCC (6 g, 45-60% B in A over 6 CV) to afford **CRBN5-SNAP2-1C-PIP** as a yellow fluffy solid (5.0 mg, 41%). **$^1$H NMR** (500 MHz, DMSO-*d$_6$*) δ 1.21 (q, *J* = 11.6 Hz, 2H), 1.71 (d, *J* = 12.5 Hz, 2H), 1.98–2.03 (m, 2H), 2.09 (d, *J* = 7.1 Hz, 2H), 2.53–2.61 (m, 2H), 2.88 (ddd, *J* = 16.7, 13.8, 5.3 Hz, 1H), 2.96 (t, *J* = 11.4 Hz, 2H), 4.03 (d, *J* = 13.1 Hz, 2H), 4.26 (d, *J* = 5.9 Hz, 2H), 5.06 (dd, *J* = 12.8, 5.4 Hz, 1H), 5.29 (s, 2H), 6.13 (s, 1H), 7.07–7.15 (m, 2H), 7.21–7.27 (m, 3H), 7.31 (d, *J* = 2.3 Hz, 1H), 7.37–7.40 (m, 2H), 7.65 (d, *J* = 8.5 Hz, 1H), 8.35 (t, *J* = 5.9 Hz, 1H), 11.08 (s, 1H). **$^{13}$C NMR** (126 MHz, DMSO-*d$_6$*) δ 172.8, 170.9, 170.3, 170.1, 167.7, 167.0, 162.8, 160.0, 154.9, 139.7, 134.7, 134.1, 128.4, 127.3, 125.0, 117.6, 117.4, 107.8, 94.4, 67.3, 48.7, 47.3, 42.1, 41.8, 32.9, 31.0, 30.8, 22.2. **MS** (m/z): [M + H]$^+$ calcd. for $C_{32}H_{33}ClN_7O_6$, 646.218; found 646.220.

***Rac-N*-(4-(((2-amino-6-chloropyrimidin-4-yl)oxy)methyl)benzyl)-3-(1-(2-(2,6-dioxopiperidin-3-yl)-1,3-dioxoisoindolin-5-yl)piperidin-4-yl)propenamide (CRBN5-SNAP2-2C-PIP)**

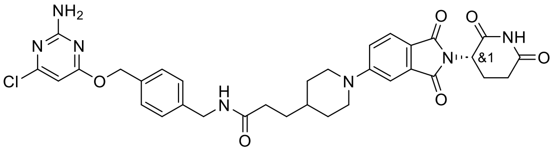

SNAP2 ligand (10 mg, 0.04 mmol), **7c** (17 mg, 0.04 mmol) and DIPEA (6 eq.) were used to synthesise **CRBN5-SNAP2-2C-PIP** according to general procedure H. The reaction was stirred for 3 h. The crude was purified by RPFCC (6 g, 50-70% B in A over 6 CV) to afford **CRBN5-SNAP2-2C-PIP** as a yellow fluffy solid (10.5 mg, 42%). **$^1$H NMR** (500 MHz, DMSO-*d$_6$*) δ 1.14 (q, *J* = 11.8 Hz, 2H), 1.48 (d, *J* = 7.5 Hz, 3H), 1.73 (d, *J* = 12.7 Hz, 2H), 1.97–2.04 (m, 1H), 2.17 (t, *J* = 7.2 Hz, 2H), 2.53–2.62 (m, 2H), 2.88 (dtd, *J* = 16.1, 13.1, 4.0 Hz, 3H), 4.04 (d, *J* = 13.1 Hz, 2H), 4.25 (d, *J* = 5.9 Hz, 2H), 5.06 (dd, *J* = 12.8, 5.4 Hz, 1H), 5.28 (s, 2H), 6.11 (s, 1H), 7.10 (s, 2H), 7.20–7.27 (m, 3H), 7.30 (d, *J* = 2.3 Hz, 1H), 7.37–7.40 (m, 2H), 7.64 (d, *J* = 8.5 Hz, 1H), 8.34 (t, *J* = 6.0 Hz, 1H), 11.08 (s, 1H). **MS** (m/z): [M + H]$^+$ calcd. for $C_{33}H_{35}ClN_7O_6$, 660.234; found 660.234.

***Rac-N*-(4-(((2-amino-6-chloropyrimidin-4-yl)oxy)methyl)benzyl)-4-(1-(2-(2,6-dioxopiperidin-3-yl)-1,3-dioxoisoindolin-5-yl)piperidin-4-yl)butanamide (CRBN5-SNAP2-3C-PIP)**

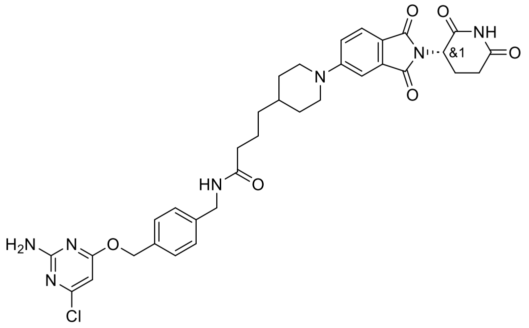

SNAP2 ligand (10 mg, 0.04 mmol), **5d** (17.5 mg, 0.04 mmol) and DIPEA (6 eq.) were used to synthesise **CRBN5-SNAP2-3C-PIP** according to general procedure H. The reaction was stirred overnight. The crude was purified by RPFCC (6 g, 50-75% B in A over 8 CV) to afford **CRBN5-SNAP2-3C-PIP** as a yellow fluffy solid (10.6 mg, 42%). **$^1$H NMR** (500 MHz, DMSO-*d$_6$*) δ 1.08–1.24 (m, 4H), 1.47–1.60 (m, 3H), 1.73 (d, *J* = 12.8 Hz, 2H), 2.00 (ddd, *J* = 11.2, 6.0, 3.6 Hz, 1H), 2.12 (t, *J* = 7.4 Hz, 2H), 2.53–2.61 (m, 2H), 2.83–2.97 (m, 3H), 4.03 (d, *J* = 12.9 Hz, 2H), 4.24 (d, *J* = 5.9 Hz, 2H), 5.06 (dd, *J* = 12.8, 5.4 Hz, 1H), 5.28 (s, 2H), 6.12 (s, 1H), 7.11 (s, 2H), 7.20–7.27

(m, 3H), 7.30 (d, *J* = 2.3 Hz, 1H), 7.36–7.40 (m, 2H), 7.64 (d, *J* = 8.6 Hz, 1H), 8.31 (t, *J* = 5.9 Hz, 1H), 11.08 (s, 1H). **MS** (m/z): [M + H]$^+$ calcd. for $C_{34}H_{37}ClN_7O_6$, 674.249; found 674.248.

#### ***Rac-N*-(4-(((2-amino-6-chloropyrimidin-4-yl)oxy)methyl)benzyl)-2-(4-(2-(2,6-dioxopiperidin-3-yl)-1,3-dioxoisoindolin-5-yl)piperazin-1-yl)acetamide (CRBN5-SNAP2-1C-PIZ)**

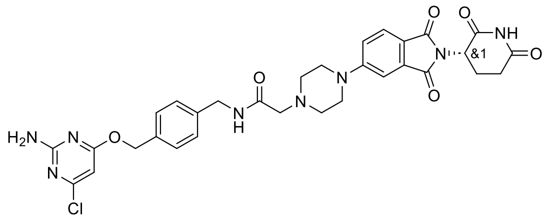

SNAP2 ligand (10 mg, 0.04 mmol), **7d** (17.9 mg, 0.04 mmol) and DIPEA (10 eq.) were used to synthesise **CRBN5-SNAP2-1C-PIZ** according to general procedure H. The reaction was stirred overnight. The crude was purified by RPFCC (6 g, 30-43% B in A over 6 CV) to afford **CRBN5-SNAP2-1C-PIZ** as a yellow fluffy solid (9.7 mg, 40%). **$^1$H NMR** (500 MHz, DMSO-*d$_6$*) δ 2.00–2.04 (m, 1H), 2.52–2.76 (m, 6H), 2.84–2.93 (m, 1H), 3.10 (s, 2H), 3.51 (d, *J* = 9.7 Hz, 4H), 4.32 (d, *J* = 6.1 Hz, 2H), 5.07 (dd, *J* = 12.8, 5.4 Hz, 1H), 5.29 (s, 2H), 6.13 (s, 1H), 7.11 (s, 2H), 7.25–7.31 (m, 3H), 7.38 (t, *J* = 7.6 Hz, 3H), 7.69 (d, *J* = 8.5 Hz, 1H), 8.45 (s, 1H), 11.09 (s, 1H). **$^{13}$C NMR** (126 MHz, DMSO-*d$_6$*) δ 172.8, 170.9, 170.3, 170.1, 167.7, 167.0, 162.8, 160.0, 154.9, 139.7, 134.7, 134.1, 128.4, 127.3, 125.0, 117.6, 117.4, 107.8, 94.4, 67.3, 48.7, 47.3, 42.1, 41.8, 32.9, 31.0, 30.8, 22.2. **MS** (m/z): [M + H]$^+$ calcd. for $C_{31}H_{32}ClN_8O_6$, 647.213; found 647.217.

#### ***Rac-N*-(4-(((2-amino-6-chloropyrimidin-4-yl)oxy)methyl)benzyl)-3-(4-(2-(2,6-dioxopiperidin-3-yl)-1,3-dioxoisoindolin-5-yl)piperazin-1-yl)propenamide (CRBN5-SNAP2-2C-PIZ)**

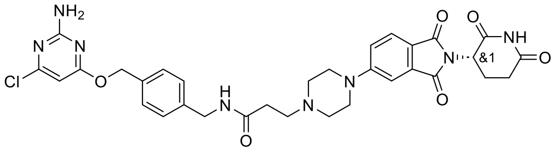

SNAP2 ligand (10 mg, 0.04 mmol), **7e** (18.41 mg, 0.04 mmol) and DIPEA (6 eq.) were used to synthesise **CRBN5-SNAP2-2C-PIZ** according to general procedure H. The reaction was stirred overnight. The crude was purified by RPFCC (6 g, 35-50% B in A over 8 CV) to afford **CRBN5-SNAP2-2C-PIZ** as a yellow fluffy solid (9.2 mg, 37%). **$^1$H NMR** (500 MHz, DMSO-*d$_6$*) δ 1.99–2.04 (m, 1H), 2.53–2.60 (m, 4H), 2.88 (ddd, *J* = 17.0, 13.8, 5.4 Hz, 1H), 3.40 (s, 10H), 4.28 (d, *J* = 5.8 Hz, 2H), 5.07 (dd, *J* = 12.9, 5.5 Hz, 1H), 5.27 (s, 2H), 6.09 (s, 1H), 7.08 (s, 2H), 7.28 (d, *J* = 8.1 Hz, 3H), 7.37 (d, *J* = 8.1 Hz, 3H), 7.69 (d, *J* = 8.5 Hz, 1H), 8.45 (s, 1H), 11.08 (s, 1H). **MS** (m/z): [M + H]$^+$ calcd. for $C_{32}H_{34}ClN_8O_6$, 661.229; found 661.229.

#### ***Rac-N*-(4-(((2-amino-6-chloropyrimidin-4-yl)oxy)methyl)benzyl)-2-((2-(2,6-dioxopiperidin-3-yl)-1,3-dioxoisoindolin-4-yl)oxy)acetamide (CRBN4-SNAP2-1C)**

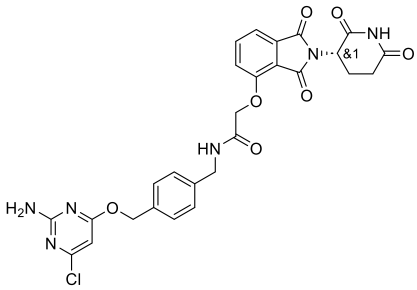

SNAP2 ligand (10 mg, 0.04 mmol), **6f** (12.6 mg, 0.04 mmol) and DIPEA (4 eq.) were used to synthesise **CRBN4-SNAP2-1C** according to general procedure H. The reaction was stirred overnight. The crude was purified by RPFCC (6 g, 35-50% B in A over 6 CV) to afford **CRBN4-SNAP2-1C** as a white fluffy solid (4.7 mg, 21%). **$^1$H NMR** (500 MHz, DMSO-*d$_6$*) δ 2.02–2.07 (m, 1H), 2.55–2.62 (m, 2H), 2.85–2.93 (m, 1H), 4.35 (d, *J* = 6.0 Hz, 2H), 4.81 (s, 2H), 5.13 (dd, *J* = 12.8, 5.4 Hz, 1H), 5.28 (s, 2H), 6.13 (s, 1H), 7.11 (s, 2H), 7.27 (d, *J* = 8.0 Hz, 2H), 7.35–7.42 (m, 3H), 7.47 (d, *J* = 2.3 Hz, 1H), 7.87 (d, *J* = 8.3 Hz, 1H), 8.78 (t, *J* = 6.0 Hz, 1H), 11.13 (s, 1H). **MS** (m/z): [M + H]$^+$ calcd. for $C_{27}H_{24}ClN_6O_7$, 579.140; found 579.140.

***Rac-N*-(4-(((2-amino-6-chloropyrimidin-4-yl)oxy)methyl)benzyl)-5-((2-(2,6-dioxopiperidin-3-yl)-1,3-dioxoisoindolin-4-yl)oxy)pentanamide (CRBN4-SNAP2-4C)**

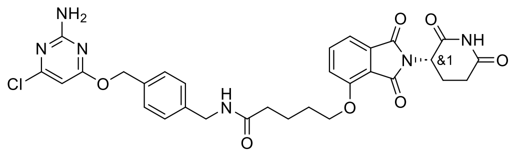

SNAP2 ligand (10 mg, 0.04 mmol), **6g** (14.14 mg, 0.04 mmol) and DIPEA (6 eq.) were used to synthesise **CRBN4-SNAP2-4C** according to general procedure H. The reaction was stirred for 3 h. The crude was purified by RPFCC (6 g, 45-70% B in A over 6 CV) to afford **CRBN4-SNAP2-4C** as a white fluffy solid (10.6 mg, 45%). **$^1$H NMR** (500 MHz, DMSO-$d_6$) δ 1.69–1.80 (m, 4H), 1.99–2.05 (m, 1H), 2.23 (t, *J* = 7.0 Hz, 2H), 2.51–2.61 (m, 2H), 2.88 (ddd, *J* = 16.8, 13.8, 5.4 Hz, 1H), 4.21 (t, *J* = 6.0 Hz, 2H), 4.26 (d, *J* = 5.9 Hz, 2H), 5.07 (dd, *J* = 12.8, 5.5 Hz, 1H), 5.28 (s, 2H), 6.12 (s, 1H), 7.10 (s, 2H), 7.25 (d, *J* = 8.1 Hz, 2H), 7.37 (d, *J* = 8.2 Hz, 2H), 7.44 (d, *J* = 7.2 Hz, 1H), 7.50 (d, *J* = 8.5 Hz, 1H), 7.81 (dd, *J* = 8.5, 7.2 Hz, 1H), 8.32 (s, 1H), 11.10 (s, 1H). **MS** (m/z): $[M + H]^+$ calcd. for $C_{30}H_{30}ClN_6O_7$, 621.187; found 621.185.

***Rac-N*-(4-(((2-amino-6-chloropyrimidin-4-yl)oxy)methyl)benzyl)-6-((2-(2,6-dioxopiperidin-3-yl)-1,3-dioxoisoindolin-4-yl)oxy)hexanamide (CRBN4-SNAP2-5C)**

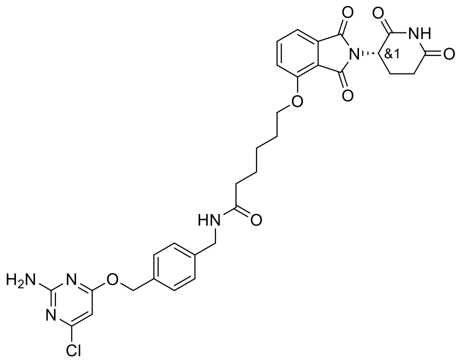

SNAP2 ligand (10 mg, 0.04 mmol), **6h** (14.7 mg, 0.04 mmol) and DIPEA (6 eq.) were used to **CRBN4-SNAP2-5C** according to general procedure H. The reaction was stirred for 3 h. The crude was purified by RPFCC (6 g, 40-55% B in A over 8 CV) to afford **CRBN4-SNAP2-5C** as a white fluffy solid (5.6 mg, 23%). **$^1$H NMR** (500 MHz, DMSO-$d_6$) δ 1.43–1.51 (m, 2H), 1.60 (p, *J* = 7.5 Hz, 2H), 1.76 (p, *J* = 6.7 Hz, 2H), 2.02 (d, *J* = 7.2 Hz, 1H), 2.16 (t, *J* = 7.4 Hz, 1H), 2.33 (d, *J* = 7.4 Hz, 1H), 2.52–2.62 (m, 2H), 2.83–2.92 (m, 1H), 4.17–4.22 (m, 2H), 4.25 (d, *J* = 5.8 Hz, 2H), 5.07 (dd, *J* = 12.8, 5.4 Hz, 1H), 5.27 (s, 2H), 6.11 (s, 1H), 7.09 (s, 2H), 7.24 (d, *J* = 8.0 Hz, 2H), 7.37 (d, *J* = 8.0 Hz, 2H), 7.44 (d, *J* = 7.2 Hz, 1H), 7.51 (d, *J* = 8.6 Hz, 1H), 7.80 (dd, *J* = 8.6, 7.2 Hz, 1H), 8.30 (t, *J* = 5.8 Hz, 1H), 11.09 (s, 1H). **MS** (m/z): $[M + H]^+$ calcd. for $C_{31}H_{32}ClN_6O_7$, 635.202; found 635.202.

***Rac-N*-(4-(((2-amino-6-chloropyrimidin-4-yl)oxy)methyl)benzyl)-7-((2-(2,6-dioxopiperidin-3-yl)-1,3-dioxoisoindolin-4-yl)oxy)heptanamide (CRBN4-SNAP2-6C)**

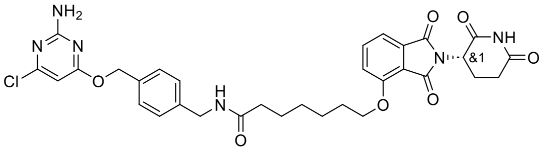

SNAP2 ligand (10 mg, 0.04 mmol), **6i** (15.2 mg, 0.04 mmol) and DIPEA (6 eq.) were used to synthesise **CRBN4-SNAP2-6C** according to general procedure H. The reaction was stirred for 3 h. The crude was purified by RPFCC (6 g, 40-55% B in A over 8 CV) to afford **CRBN4-SNAP2-6C** as a white solid (9.5 mg, 39%). **$^1$H NMR** (500 MHz, DMSO-$d_6$) δ 1.33 (q, *J* = 7.7 Hz, 2H), 1.46 (p, *J* = 7.4 Hz, 2H), 1.54 (q, *J* = 7.6 Hz, 2H), 1.74 (t, *J* = 7.4 Hz, 2H), 1.99–2.05 (m, 1H), 2.14 (t, *J* = 7.4 Hz, 2H), 2.52–2.62 (m, 2H), 2.84–2.92 (m, 1H), 4.19 (t, *J* = 6.3 Hz, 2H), 4.24 (d, *J* = 5.9 Hz, 2H), 5.07 (dd, *J* = 12.8, 5.4 Hz, 1H), 5.27 (s, 2H), 6.12 (s, 1H), 7.09 (s, 2H), 7.24 (d, *J* = 7.9 Hz, 2H), 7.37 (d, *J* = 8.1 Hz, 2H), 7.44 (d, *J* = 7.3 Hz, 1H), 7.50 (d, *J* = 8.5 Hz, 1H), 7.80 (dd, *J* = 8.6, 7.2 Hz, 1H), 8.28 (t, *J* = 5.9 Hz, 1H), 11.09 (s, 1H). **MS** (m/z): $[M + H]^+$ calcd. for $C_{32}H_{34}ClN_6O_7$, 649.218; found 649.219.

**Rac-N-(4-(((2-amino-6-chloropyrimidin-4-yl)oxy)methyl)benzyl)-8-((2-(2,6-dioxopiperidin-3-yl)-1,3-dioxoisoindolin-4-yl)oxy)octanamide (CRBN4-SNAP2-7C)**

SNAP2 ligand (10 mg, 0.04 mmol), **6j** (15.7 mg, 0.04 mmol) and DIPEA (6 eq.) were used to synthesise **CRBN4-SNAP2-7C** according to general procedure H. The reaction was stirred for 3 h. The crude was purified by RPFCC (6 g, 45-65% B in A over 8 CV) to afford **CRBN4-SNAP2-7C** as a white fluffy solid (11.2 mg, 45%).

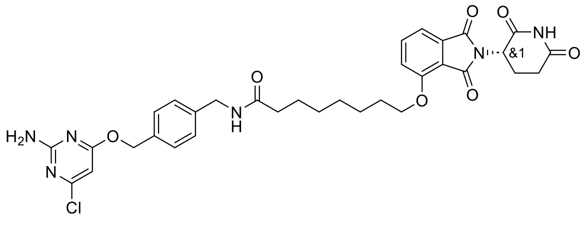

**$^1$H NMR** (500 MHz, DMSO-$d_6$) δ 1.24–1.37 (m, 4H), 1.43 (q, $J$ = 7.5 Hz, 2H), 1.53 (p, $J$ = 7.4 Hz, 2H), 1.74 (p, $J$ = 6.6 Hz, 2H), 1.98–2.05 (m, 1H), 2.13 (t, $J$ = 7.4 Hz, 2H), 2.52–2.63 (m, 2H), 2.87 (ddd, $J$ = 16.8, 13.8, 5.4 Hz, 1H), 4.19 (t, $J$ = 6.4 Hz, 2H), 4.24 (d, $J$ = 5.9 Hz, 2H), 5.07 (dd, $J$ = 12.8, 5.4 Hz, 1H), 5.27 (s, 2H), 6.11 (s, 1H), 7.09 (s, 2H), 7.20–7.28 (m, 2H), 7.34–7.40 (m, 2H), 7.43 (d, $J$ = 7.2 Hz, 1H), 7.50 (d, $J$ = 8.6 Hz, 1H), 7.80 (dd, $J$ = 8.5, 7.2 Hz, 1H), 8.28 (t, $J$ = 5.9 Hz, 1H), 11.10 (s, 1H). **MS** (m/z): [M + H]$^+$ calcd. for $C_{33}H_{36}ClN_6O_7$, 663.233; found 663.235.

**Rac-N-(4-(((2-amino-6-chloropyrimidin-4-yl)oxy)methyl)benzyl)-2-((2-(2,6-dioxopiperidin-3-yl)-1,3-dioxoisoindolin-5-yl)oxy)acetamide (CRBN5-SNAP2-1C)**

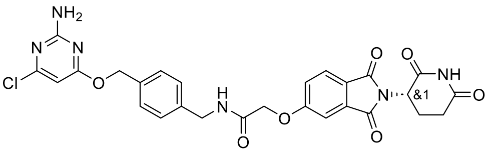

SNAP2 ligand (10 mg, 0.04 mmol), **7f** (12.6 mg, 0.04 mmol) and DIPEA (4 eq.) were used to synthesise **CRBN5-SNAP2-1C** according to general procedure H. The reaction was stirred overnight. The crude was purified by RPFCC (6 g, 30-50% B in A over 8 CV) to afford **CRBN5-SNAP2-1C** as a white fluffy solid (11.1 mg, 51%). **$^1$H NMR** (500 MHz, DMSO-$d_6$) δ 2.03 (ddd, $J$ = 12.8, 6.3, 2.3 Hz, 1H), 2.53–2.62 (m, 2H), 2.89 (ddd, $J$ = 16.9, 13.8, 5.4 Hz, 1H), 4.36 (d, $J$ = 6.1 Hz, 2H), 4.87 (s, 2H), 5.11 (dd, $J$ = 12.9, 5.4 Hz, 1H), 5.29 (s, 2H), 6.14 (s, 1H), 7.12 (s, 2H), 7.28 (d, $J$ = 7.9 Hz, 2H), 7.39 (dd, $J$ = 8.2, 5.2 Hz, 3H), 7.50 (d, $J$ = 7.3 Hz, 1H), 7.80 (dd, $J$ = 8.5, 7.2 Hz, 1H), 8.54 (t, $J$ = 6.1 Hz, 1H), 11.12 (s, 1H). **MS** (m/z): [M + H]$^+$ calcd. for $C_{27}H_{24}ClN_6O_7$, 579.140; found 579.143.

**Rac-N-(4-(((2-amino-6-chloropyrimidin-4-yl)oxy)methyl)benzyl)-5-((2-(2,6-dioxopiperidin-3-yl)-1,3-dioxoisoindolin-5-yl)oxy)pentanamide (CRBN5-SNAP2-4C)**

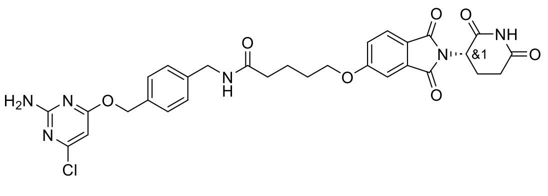

SNAP2 ligand (10 mg, 0.04 mmol), **7g** (14.1 mg, 0.04 mmol) and DIPEA (6 eq.) were used to synthesise **CRBN5-SNAP2-4C** according to general procedure H. The reaction was stirred for 3 h. The crude was purified by RPFCC (6 g, 45-80% B in A over 8 CV) to afford **CRBN5-SNAP2-4C** as a white fluffy solid (5.1 mg, 22%). **$^1$H NMR** (500 MHz, DMSO-$d_6$) δ 1.66–1.78 (m, 4H), 2.01–2.07 (m, 1H), 2.22 (t, $J$ = 7.2 Hz, 2H), 2.51–2.62 (m, 2H), 2.89 (ddd, $J$ = 16.9, 13.8, 5.4 Hz, 1H), 4.18 (t, $J$ = 6.2 Hz, 2H), 4.26 (d, $J$ = 5.9 Hz, 2H), 5.12 (dd, $J$ = 12.8, 5.4 Hz, 1H), 5.28 (s, 2H), 6.12 (s, 1H), 7.10 (s, 2H), 7.25 (d, $J$ = 8.0 Hz, 2H), 7.33–7.39 (m, 3H), 7.42 (d, $J$ = 2.3 Hz, 1H), 7.82 (s, 1H), 8.34 (t, $J$ = 5.9 Hz, 1H), 11.11 (s, 1H). **MS** (m/z): [M + H]$^+$ calcd. for $C_{30}H_{30}ClN_6O_7$, 621.187; found 621.186.

#### ***Rac-N*-(4-(((2-amino-6-chloropyrimidin-4-yl)oxy)methyl)benzyl)-7-((2-(2,6-dioxopiperidin-3-yl)-1,3-dioxoisoindolin-5-yl)oxy)heptanamide (CRBN5-SNAP2-6C)**

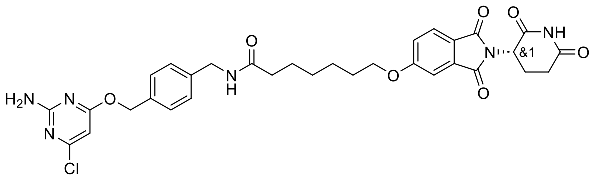

SNAP2 ligand (10 mg, 0.04 mmol), **7h** (15.2 mg, 0.04 mmol) and DIPEA (6 eq.) were used to synthesise **CRBN5-SNAP2-6C** according to general procedure H. The reaction was stirred for 3 h. The crude was purified by RPFCC (6 g, 40-55% B in A over 8 CV) to afford **CRBN5-SNAP2-6C** as a white solid (6.8 mg, 28%). **$^1$H NMR** (500 MHz, DMSO-*d$_6$*) δ 1.33 (q, *J* = 7.7 Hz, 2H), 1.42 (p, *J* = 7.1 Hz, 2H), 1.55 (p, *J* = 7.3 Hz, 2H), 1.74 (t, *J* = 7.4 Hz, 2H), 2.01–2.07 (m, 1H), 2.14 (t, *J* = 7.4 Hz, 2H), 2.52–2.62 (m, 2H), 2.85–2.93 (m, 1H), 4.16 (t, *J* = 6.4 Hz, 2H), 4.25 (d, *J* = 5.9 Hz, 2H), 5.11 (dd, *J* = 12.9, 5.4 Hz, 1H), 5.27 (s, 2H), 6.11 (s, 1H), 7.09 (s, 2H), 7.24 (d, *J* = 7.9 Hz, 2H), 7.32–7.39 (m, 3H), 7.42 (d, *J* = 2.3 Hz, 1H), 7.82 (d, *J* = 8.3 Hz, 1H), 8.29 (s, 1H), 11.10 (s, 1H). **MS** (m/z): $[M + H]^+$ calcd. for $C_{32}H_{34}ClN_6O_7$, 649.218; found 649.218.

#### ***Rac*-N-(4-(((2-amino-6-chloropyrimidin-4-yl)oxy)methyl)benzyl)-8-((2-(2,6-dioxopiperidin-3-yl)-1,3-dioxoisoindolin-5-yl)oxy)octanamide (CRBN5-SNAP2-7C)**

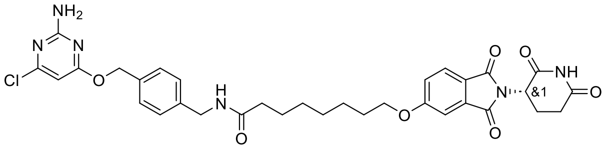

SNAP2 ligand (10 mg, 0.04 mmol), **7i** (15.7 mg, 0.04 mmol) and DIPEA (6 eq.) were used to synthesise **CRBN5-SNAP2-7C** according to general procedure H. The reaction was stirred for 3 h. The crude was purified by RPFCC (6 g, 45-65% B in A over 8 CV) to afford **CRBN5-SNAP2-7C** as a white fluffy solid (13.1 mg, 52%). **$^1$H NMR** (500 MHz, DMSO-*d$_6$*) δ 1.25–1.37 (m, 4H), 1.41 (p, *J* = 7.0 Hz, 2H), 1.53 (p, *J* = 7.4 Hz, 2H), 1.74 (p, *J* = 6.7 Hz, 2H), 2.01–2.08 (m, 1H), 2.13 (t, *J* = 7.4 Hz, 2H), 2.52–2.63 (m, 2H), 2.89 (ddd, *J* = 17.0, 13.8, 5.4 Hz, 1H), 4.16 (t, *J* = 6.5 Hz, 2H), 4.24 (d, *J* = 6.1 Hz, 2H), 5.11 (dd, *J* = 12.8, 5.4 Hz, 1H), 5.27 (s, 2H), 6.11 (s, 1H), 7.09 (s, 2H), 7.24 (d, *J* = 8.1 Hz, 2H), 7.34 (dd, *J* = 8.4, 2.3 Hz, 1H), 7.36–7.39 (m, 2H), 7.42 (d, *J* = 2.3 Hz, 1H), 7.82 (d, *J* = 8.3 Hz, 1H), 8.29 (t, *J* = 6.0 Hz, 1H), 11.11 (s, 1H). **MS** (m/z): $[M + H]^+$ calcd. for $C_{33}H_{36}ClN_6O_7$, 663.233; found 663.235.

### SNAP ligand variations

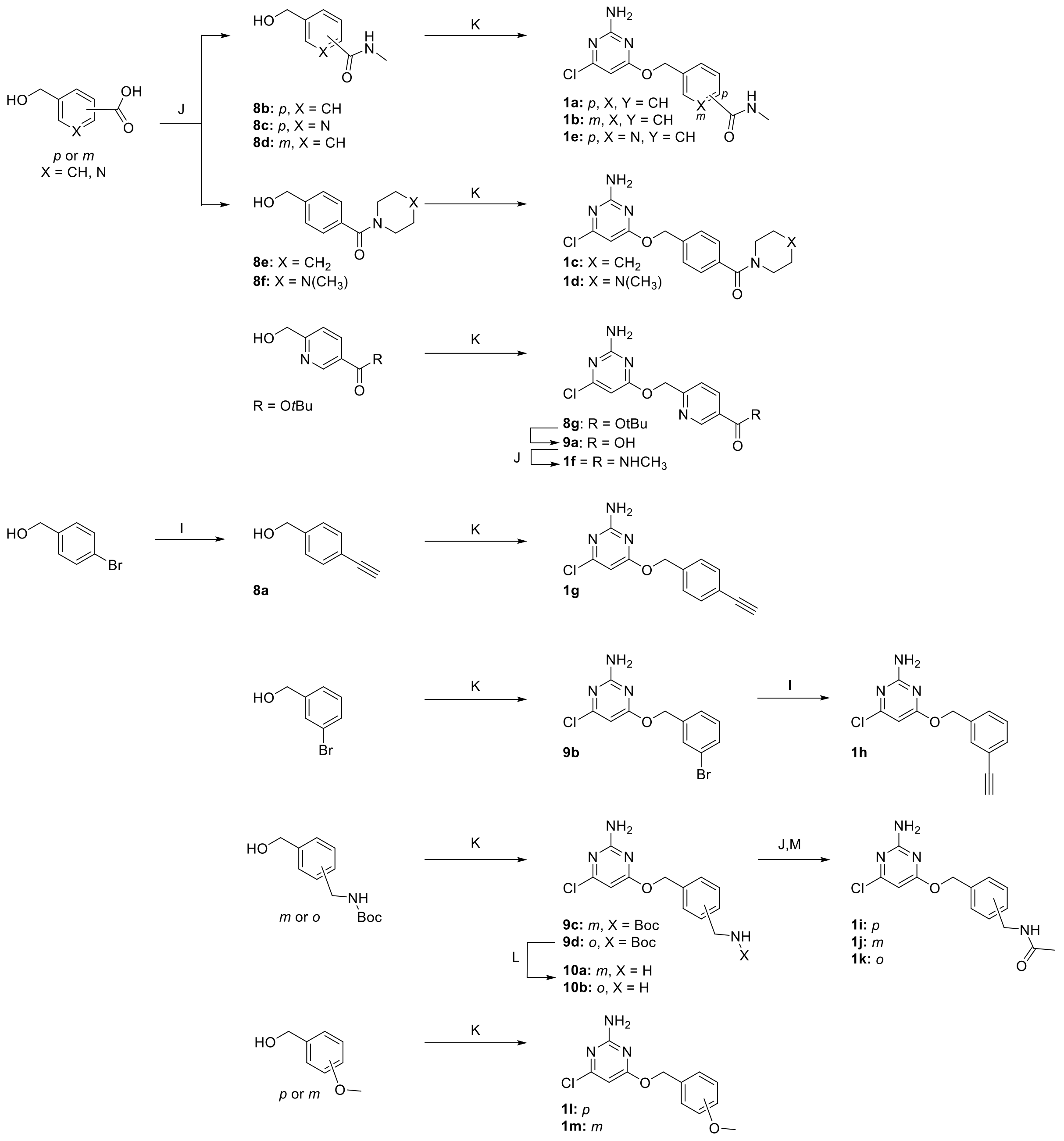

**Scheme S4**. Overview of synthetic route for SNAP ligand variations.

#### General procedure I

The desired aryl bromide (1.0 eq.), CuI (0.2 eq.) and $Pd(PPh_3)_4$ (0.1 eq.) in a sealed vessel under argon were dissolved in THF (5 mL) before addition of $Et_3N$ (2.0 eq.) and trimethylsilyl acetylene (2.0 eq.) were stirred at 70 °C for 16 h. The reaction mixture was filtered through a pad of celite and concentrated before redissolving in MeOH (5 mL) and addition of potassium carbonate (1 eq.). The reaction mixture was stirred at room temperature for 2 h. The reaction mixture was concentrated and partitioned between ammonium chloride and EtOAc, the layers separated, and the aqueous phase extracted twice with EtOAc. The combined organic phase was dried over $MgSO_4$, filtered, concentrated, and purified by NPFCC to afford products **8a**, **1h**.

**General procedure J**

The desired acid (1.0 eq.), the desired amine (1.1-3 eq.) and HATU (1 eq.) were dissolved in DCM (1 mL) before addition of DIPEA (3-5 eq.). The reaction mixture was stirred at room temperature until completion monitored by LCMS. The residue was purified by RPFCC to afford products **8b-f**, **1f**, **1i**.

**General procedure K**

The desired alcohol (1.0 eq.) and 4,6-dichloropyrimidin-2-amine (1.2 eq.) were dissolved in THF (3 mL) before addition of KHMDS (1.2 eq.). The reaction mixture was stirred at 40 °C until completion or until no further conversion was observed, monitored by LCMS. The reaction mixture was quenched by addition of ammonium chloride solution and purified by NPFCC or RPFCC afford products **1a**-**e**, **1g**, **1l**-**m**, **8g**, **9b**-**d**.

**General procedure L**

The desired Boc-protected amine (1.0 eq.) was dissolved in DCM (0.5 mL) before addition of HCl (4 M in dioxane, 10 eq.). The reaction mixture was stirred at room temperature for 2 h. The reaction mixture was concentrated to afford products **10a**-**b** to be used directly in the next step without further purification.

**General procedure M**

The desired amine (1.0 eq.), PyOxim (1.5 eq.) and OXYMA Pure (0.5 eq.) were dissolved in DMF (1 mL) before addition of acetic acid (1.5 eq.) and DIPEA (5 eq.). The reaction mixture was stirred at room temperature for 16 h until completion monitored by LCMS. The residue was purified on RPFCC to afford products **1j**-**k**.

**(4-Ethynylphenyl)methanol (8a)**

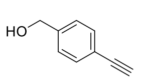


(4-Bromophenyl)methanol (400 mg, 2.14 mmol) was used to synthesise **8a** according to general procedure I. The crude product was purified by NPFCC (12 g, 0–17% EtOAc in heptane over 35 CV) to afford **8a** as a brown oil (67 mg, 24%); **$^1$H NMR** (500 MHz, Chloroform-*d*) δ 3.09 (s, 1H), 5.09 (s, 2H), 7.30 (d, *J* = 8.2 Hz, 2H), 7.48 (d, *J* = 8.2 Hz, 2H); **$^{13}$C NMR** (126 MHz, Chloroform-*d*) δ 136.7, 132.4, 128.1, 122.1, 83.3, 77.8, 65.8; **MS** (m/z): $[M + H]^+$ calcd. for $C_9H_8O$, 133.1; found 133.0.

**4-(Hydroxymethyl)-*N*-methylbenzamide (8b)**

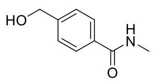


4-(Hydroxymethyl)benzoic acid (250 mg, 1.64 mmol) and methylamine hydrochloride (1.1 eq.) were used to synthesise **8b** according to general procedure J with DIPEA (4 eq.) and stirred for 1 h. The crude product was purified by RPFCC twice (12 g, 100% A over 10 CV, 0–3 B in A over 5 CV then by 12 g, 100% 0.1% ammonia in water over 10 CV, 0–10% B in 0.1% ammonia in water over 12 CV) to afford **8b** as a white solid (107 mg, 39%); **$^1$H NMR** (500 MHz, DMSO-d$_6$) δ 2.77 (d, *J* = 4.6 Hz, 3H), 4.54 (d, *J* = 5.5 Hz, 2H), 5.29 (t, *J* = 5.7 Hz, 1H), 7.35–7.4 (m, 2H), 7.77–7.81 (m, 2H), 8.38 (q, *J* = 4.6 Hz, 1H); **$^{13}$C NMR** (126 MHz, DMSO-*d$_6$*) δ 166.5, 145.7, 132.9, 126.9, 126.0, 62.5, 26.2; **MS** (m/z): $[M + H]^+$ calcd. for $C_9H_{11}NO_2$, 166.1; found 166.2.

**5-(Hydroxymethyl)-*N*-methylpicolinamide (8c)**

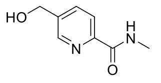


5-(Hydroxymethyl)picolinic acid (250 mg, 1.63 mmol) and methylamine hydrochloride (1.1 eq.) were used to synthesise **8c** according to general procedure J with DIPEA (4 eq.) and stirred for 2 h. The crude product was purified by RPFCC (12 g, 100% 0.1% A over 10 CV, 0–5 B in A over 10 CV) to afford **8c** as a (429 mg, 45 w%, 71%) to be used directly in the next step without further purification; **$^1$H NMR**

(500 MHz, DMSO-$d_6$) δ 2.81 (d, *J* = 4.9 Hz, 3H), 4.61 (s, 2H), 5.47 (s, 1H), 7.89 (dd, *J* = 8.0, 2.1 Hz, 1H), 7.99 (d, *J* = 8.0 Hz, 1H), 8.56 (d, *J* = 2.1 Hz, 1H), 8.75 (q, *J* = 4.7 Hz, 1H); **$^{13}$C NMR** (126 MHz, DMSO-$d_6$) δ 164.4, 148.8, 146.7, 140.6, 135.6, 121.3, 60.4, 40.4, 26.0; ; **MS** (m/z): [M + H]$^+$ calcd. for $C_8H_{11}N_2O_2$, 167.1; found 167.0

**3-(Hydroxymethyl)-*N*-methylbenzamide (8d)**

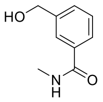

3-(Hydroxymethyl)benzoic acid (100 mg, 0.66 mmol) and methylamine hydrochloride (1.1 eq.) were used to synthesise **8d** according to general procedure J with DIPEA (4 eq.) and stirred overnight. The crude product was purified by RPFCC (12 g, 100% A over 5 CV, 0–5% B in A over 15 CV) to afford **8d** as a white solid (70 mg, 65%); **$^1$H NMR** (500 MHz, DMSO-$d_6$) δ 2.77 (d, *J* = 4.6 Hz, 3H), 4.53 (d, *J* = 5.7 Hz, 2H), 5.28 (t, *J* = 5.7 Hz, 1H), 7.39 (t, *J* = 7.6 Hz, 1H), 7.45 (dt, *J* = 7.6, 1.4 Hz, 1H), 7.66–7.7 (m, 1H), 7.77–7.81 (m, 1H), 8.38–8.44 (m, 1H); **$^{13}$C NMR** (126 MHz, DMSO-$d_6$) δ 166.7, 142.7, 134.4, 129.0, 128.0, 125.3, 125.2, 62.7, 26.3; **MS** (m/z): [M + H]$^+$ calcd. for $C_8H_{10}N_2O_2$, 167.1; found 167.0.

**(4-(Hydroxymethyl)phenyl)(piperidin-1-yl)methanone (8e)**

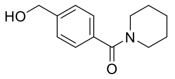

4-(Hydroxymethyl)benzoic acid (161 mg, 1.06 mmol) and piperidine hydrochloride (3 eq.) were used to synthesise **8e** according to general procedure J with DIPEA (5 eq.) and stirred overnight. The crude product was purified by RPFCC (12, 5–50% B in A over 16 CV) to afford **8e** as a colourless film/gum (196 mg, 84%); **$^1$H NMR** (500 MHz, DMSO-$d_6$) δ 1.36–1.57 (m, 4H), 1.61 (q, *J* = 8.7 Hz, 2H), 3.27 (s, 2H), 3.56 (s, 2H), 4.52 (s, 2H), 5.27 (s, 1H), 7.31 (d, *J* = 8.1 Hz, 2H), 7.36 (d, *J* = 8.1 Hz, 2H); **$^{13}$C NMR** (126 MHz, DMSO-$d_6$) δ 169.0, 143.8, 134.8, 126.5, 126.2, 62.5, 48.0, 42.3, 26.0, 25.3, 24.1; **MS** (m/z): [M + H]$^+$ calcd. for $C_{13}H_{17}NO_2$, 220.1; found 220.3.

**(4-(Hydroxymethyl)phenyl)(4-methylpiperazin-1-yl)methanone (8f)**

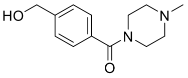

4-(Hydroxymethyl)benzoic acid (250 mg, 1.64 mmol) and 1-methylpiperazine (3 eq.) were used to synthesise **8f** according to general procedure J with DIPEA (3 eq.) and stirred overnight. The crude product was purified by RPFCC (30–60% B in A over 20 CV) to afford **8f** as a colourless film/gum (109 mg, 28%); **$^1$H NMR** (500 MHz, DMSO-$d_6$) δ 2.18 (s, 3H), 2.21–2.4 (m, 4H), 3.34–3.34 (m, 2H), 3.59 (s, 2H), 4.52 (d, *J* = 2.7 Hz, 2H), 5.28 (s, 1H), 7.33 (d, *J* = 8.1 Hz, 2H), 7.37 (d, *J* = 8.1 Hz, 2H); **$^{13}$C NMR** (126 MHz, DMSO-$d_6$) δ 169.0, 144.1, 134.2, 126.8, 126.2, 62.5, 54.7, 47.2, 45.6, 41.5; **MS** (m/z): [M + H]$^+$ calcd. for $C_{13}H_{18}N_2O_2$, 235.1; found 235.3.

***Tert*-butyl 6-(((2-amino-6-chloropyrimidin-4-yl)oxy)methyl)nicotinate (8g)**

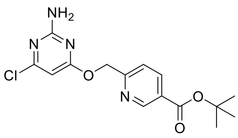

*Tert*-butyl 6-(hydroxymethyl)nicotinate was used to synthesize **8g** according to general procedure K. The reaction was stirred at 50 °C for 4 h and then stirred at room temperature overnight. The organics were concentrated, and the residue partitioned between ammonium chloride and DCM, the layers separated, and the aqueous phase extracted with DCM. The combined organic phase was dried over $MgSO_4$, filtered, concentrated and purified by RPFCC (6 g, 5–95% B in A over 15 CV) to afford **8g** as a white solid (142 mg, 59%); **$^1$H NMR** (500 MHz, DMSO-$d_6$) δ 1.56 (9H, d), 5.48 (2H, s), 6.26 (1H, d), 7.10 (2H, s), 7.55 (1H, d), 8.25 (1H, d), 9.00 (1H, s). **$^{13}$C NMR** (151 MHz, DMSO-$d_6$) δ 27.73, 39.92, 40.06, 67.38, 81.69, 94.46, 121.01, 126.11, 137.57, 149.60, 160.21, 160.32, 162.72, 163.68, 169.98. **MS** (m/z): [M + H]$^+$ calcd. for $C_{15}H_{17}ClN_4O_3$, 336.1; found 337.3.

**6-(((2-amino-6-chloropyrimidin-4-yl)oxy)methyl)nicotinic acid (9a)**

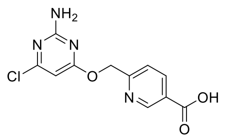

**8g** (140 mg, 0.42 mmol) was dissolved in DCM (2 mL) before addition of TFA (2.1 mL, 27 mmol) and the reaction stirred at room temperature overnight. The volatiles were removed under reduced pressure, co-evaporating with toluene (2x) and EtOAc (2x) to remove TFA to give **9a** (165 mg, 101 %) as a yellow solid, to be used directly in the next step without further purification; **$^1$H NMR** (500 MHz, DMSO-*d$_6$*) δ 5.49 (d, *J* = 1.9 Hz, 2H), 6.26 (d, *J* = 2.0 Hz, 1H), 7.09–7.14 (m, 2H), 7.55 (d, *J* = 8.2 Hz, 1H), 8.29 (d, *J* = 8.1 Hz, 1H), 9.04 (s, 1H). **$^{13}$C NMR** (151 MHz, DMSO-*d$_6$*) δ 67.40, 94.47, 121.05, 125.66, 137.89, 149.89, 160.21, 160.27, 162.74, 166.04, 170.00. **MS** (m/z): $[M + H]^+$ calcd. for $C_{11}H_9ClN_4O_3$, 280.1; found 281.1.

**4-((3-Bromobenzyl)oxy)-6-chloropyrimidin-2-amine (9b)**

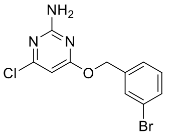

(3-Bromophenyl)methanol (38 mg, 0.20 mmol) was used to synthesise **9b** according to general procedure K and stirred for 3 h. The organics were concentrated and the residue partitioned between ammonium chloride and DCM, the layers separated and the aqueous phase extracted with DCM. The combined organic phase was dried over $MgSO_4$, filtered, concentrated and purified by NPFCC (5 g, 10–18% EtOAc in heptane over 12 CV) to afford **9b** as a white solid (31 mg, 49%); **$^1$H NMR** (500 MHz, Chloroform-*d*) δ 5.31 (s, 2H), 5.33 (s, 2H), 6.21 (s, 1H), 7.28 (t, *J* = 7.8 Hz, 1H), 7.35 (d, *J* = 7.8 Hz, 1H), 7.50 (d, *J* = 7.9 Hz, 1H), 7.58 (s, 1H); **$^{13}$C NMR** (126 MHz, Chloroform-*d*) δ 170.8, 162.3, 161.1, 138.4, 131.4, 131.0, 130.3, 126.5, 122.7, 97.4, 67.3; **MS** (m/z): $[M + H]^+$ calcd. for $C_{11}H_9BrClN_3O$, 314.0 ; found 314.1.

***Tert*-Butyl (3-(((2-amino-6-chloropyrimidin-4-yl)oxy)methyl)benzyl)carbamate (9c)**

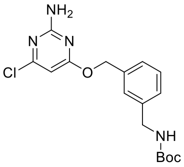

*Tert*-butyl (3-(hydroxymethyl)benzyl)carbamate (48 mg, 0.20 mmol) was used to synthesise **9c** according to general procedure K and stirred overnight. The crude product was purified by RPFCC (6 g, 45–62% B in A over 10 CV) to afford **9c** as a colourless film (45 mg, 61%); **$^1$H NMR** (500 MHz, Chloroform-*d*) δ 1.46 (s, 9H), 4.33 (d, *J* = 5.6 Hz, 2H), 4.88 (s, 1H), 5.14 (s, 2H), 5.30 (s, 2H), 6.16 (s, 1H), 7.24 (s, 1H), 7.29 (d, *J* = 7.7 Hz, 1H), 7.32–7.36 (m, 2H); **$^{13}$C NMR** (126 MHz, Chloroform-*d*) δ 171.0, 162.2, 160.9, 156.1, 136.5, 129.0, 127.5, 127.2, 97.5, 68.4, 44.6, 28.5; **MS** (m/z): $[M + H]^+$ calcd. for $C_{17}H_{21}ClN_4O_3$, 365.1; found 365.3.

***Tert*-Butyl (2-(((2-amino-6-chloropyrimidin-4-yl)oxy)methyl)benzyl)carbamate (9d)**

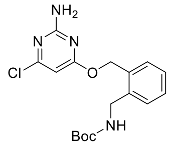

*Tert*-butyl (2-(hydroxymethyl)benzyl)carbamate (48 mg, 0.20 mmol) was used to synthesise **9d** according to general procedure K and stirred overnight. The crude product was purified by RPFCC (6 g, 45–60 % B in A over 15 CV) to afford **9d** as a colourless film (46 mg, 62%); **$^1$H NMR** (500 MHz, Chloroform-*d*) δ 1.46 (s, 9H), 4.32 (s, 1H), 4.48 (d, *J* = 5.3 Hz, 2H), 5.35 (s, 2H), 5.38 (s, 2H), 6.15 (s, 1H), 7.31 (dd, *J* = 9.8, 4.7 Hz, 3H), 7.42 (d, *J* = 6.7 Hz, 1H); **$^{13}$C NMR** (126 MHz, Chloroform-*d*) δ 170.6, 162.2, 160.8, 155.7, 136.8, 134.3, 129.6, 129.0, 128.8, 128.1, 97.4, 65.6, 42.6, 28.6; **MS** (m/z): $[M + H]^+$ calcd. for $C_{17}H_{21}ClN_4O_3$, 365.1; found 365.3.

**4-((3-(Aminomethyl)benzyl)oxy)-6-chloropyrimidin-2-amine (10a)**

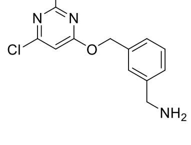

**9c** (45 mg, 0.12 mmol) was used to synthesise **10a** according to general procedure L. The crude product was concentrated to afford **10a** as a white solid (45 mg, 121%) to be used directly in the next step without further purification; **$^1$H NMR** (500 MHz, DMSO-*d$_6$*) δ 4.02 (q, *J* = 5.9 Hz, 2H), 4.77 (s, 2H), 5.58 (s, 1H), 7.17 (s, 2H), 7.4–7.5 (m, 3H), 7.55 (s, 1H), 8.43 (s, 3H); **$^{13}$C**

**NMR** (126 MHz, DMSO-*d6*) δ 162.0, 159.3, 155.5, 138.0, 134.6, 129.4, 129.0, 129.0, 128.9, 99.4, 45.9, 42.0; **MS** (m/z): $[M + H]^+$ calcd. for $C_{12}H_{13}ClN_4O$, 265.1; found 265.3.

#### 4-((2-(Aminomethyl)benzyl)oxy)-6-chloropyrimidin-2-amine (10b)

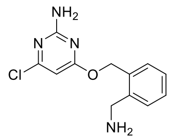

**9d** (46 mg, 0.13 mmol) was used to synthesise **10b** according to general procedure L. The crude product was concentrated to afford **10b** as a white solid (49 mg, 128%) to be used directly in the next step without further purification; $^1$**H NMR** (500 MHz, DMSO-*d6*) δ 4.16 (q, *J* = 5.9 Hz, 2H), 4.94 (s, 2H), 5.58 (s, 1H), 7.16 (s, 2H), 7.40 (td, *J* = 7.4, 1.2 Hz, 1H), 7.45 (td, *J* = 7.5, 1.5 Hz, 1H), 7.51 (dd, *J* = 7.5, 1.3 Hz, 1H), 7.54 (d, *J* = 7.0 Hz, 1H), 8.49 (s, 3H); $^{13}$**C NMR** (126 MHz, DMSO-*d6*) δ 162.0, 159.3, 155.5, 136.1, 133.0, 130.6, 129.6, 129.2, 128.9, 99.4, 44.0, 38.5; **MS** (m/z): $[M + H]^+$ calcd. for $C_{12}H_{13}ClN_4O$, 265.1; found 265.3.

#### 4-(((2-Amino-6-chloropyrimidin-4-yl)oxy)methyl)-*N*-methylbenzamide (1a)

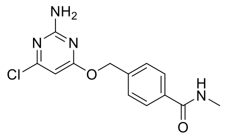

**8b** (50 mg, 0.30 mmol) was used to synthesise **1a** according to general procedure K and stirred for 2d. The organics were concentrated and the residue partitioned between ammonium chloride and DCM, the layers separated and the aqueous phase extracted with DCM. The combined organic phase was dried over $MgSO_4$, filtered, concentrated and purified by NPFCC (5 g, 50–90% EtOAc in heptane over 10 CV) to afford **1a** as a white solid (31 mg, 35%); $^1$**H NMR** (500 MHz, DMSO-*d6*) δ 2.78 (d, *J* = 4.5 Hz, 3H), 5.37 (s, 2H), 6.18 (s, 1H), 7.13 (s, 2H), 7.47–7.52 (m, 2H), 7.81–7.85 (m, 2H), 8.44 (q, *J* = 4.4 Hz, 1H); $^{13}$**C NMR** (126 MHz, DMSO-*d6*) δ 170.2, 166.3, 162.8, 160.1, 139.2, 134.2, 127.9, 127.2, 94.4, 66.8, 40.1, 40.0, 39.9, 39.9, 39.8, 39.7, 39.6, 39.5, 39.4, 39.4, 39.2, 39.0, 26.3; **MS** (m/z): $[M + H]^+$ calcd. for $C_{13}H_{13}ClN_4O_2$, 293.081; found 293.080.

#### 3-(((2-Amino-6-chloropyrimidin-4-yl)oxy)methyl)-*N*-methylbenzamide (1b)

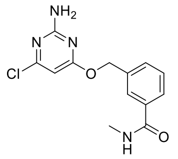

**8d** (70 mg, 0.42 mmol) was used to synthesise **1b** according to general procedure K and stirred overnight. The organics were concentrated and the residue partitioned between ammonium chloride and DCM, the layers separated and the aqueous phase extracted with DCM. The combined organic phase was dried over $MgSO_4$, filtered, concentrated and purified by NPFCC (5 g, 25–35% EtOAc in heptane over 8 CV, 35–100 % EtOAc in heptane over 5 CV, 100% EtOAc over 5 CV) to afford **1b** as a white solid (68 mg, 55%); $^1$**H NMR** (500 MHz, DMSO-*d6*) δ 2.78 (d, *J* = 4.6 Hz, 3H), 5.36 (s, 2H), 6.18 (s, 1H), 7.13 (s, 2H), 7.47 (t, *J* = 7.7 Hz, 1H), 7.55–7.59 (m, 1H), 7.79 (dt, *J* = 7.7, 1.4 Hz, 1H), 7.87 (t, *J* = 1.5 Hz, 1H), 8.46 (d, *J* = 4.4 Hz, 1H); $^{13}$**C NMR** (126 MHz, DMSO-*d6*) δ 170.3, 166.3, 162.8, 160.1, 136.5, 134.7, 130.8, 128.5, 126.9, 126.7, 94.4, 67.1, 26.3; **MS** (m/z): $[M + H]^+$ calcd. for $C_{13}H_{14}ClN_4O_2$, 293.081; found 293.080.

#### (4-(((2-Amino-6-chloropyrimidin-4-yl)oxy)methyl)phenyl)(piperidin-1-yl)methanone (1c)

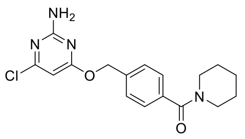

**8e** (45 mg, 0.20 mmol) was used to synthesise **1c** according to general procedure K and stirred for 2 d. The crude product was purified by RPFCC (6 g, 30–70% B in A over 14 CV) to afford **1c** as a white solid (36 mg, 52%); $^1$**H NMR** (500 MHz, DMSO-*d6*) δ 1.45 (s, 4H), 1.58–1.62 (m, 2H), 3.25 (s, 2H), 3.57 (s, 2H), 5.35 (s, 2H), 6.17 (s, 1H), 7.14 (s, 2H), 7.37 (d, *J* = 8.1 Hz, 2H), 7.48 (d, *J* = 8.1 Hz, 2H); $^{13}$**C NMR** (126 MHz, DMSO-*d6*) δ 170.3, 168.6, 162.8, 160.1, 137.3, 136.2, 128.1, 126.8, 94.4, 66.9, 48.0, 42.3, 25.9, 25.3, 24.0; **MS** (m/z): $[M + H]^+$ calcd. for $C_{17}H_{20}ClN_4O_2$, 347.128; found 347.128.

**(4-(((2-Amino-6-chloropyrimidin-4-yl)oxy)methyl)phenyl)(4-methylpiperazin-1-yl)methanone (1d)**

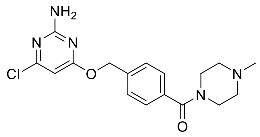

**8f** (48 mg, 0.20 mmol) was used to synthesise **1d** according to general procedure K and stirred overnight. The crude product was purified by RPFCC (6 g, 5–17% B in A over 10 CV) to afford **1d** as a colourless film/gum (30 mg, 36%); **$^1$H NMR** (500 MHz, DMSO-*d$_6$*) δ 2.19 (s, 3H), 2.22–2.43 (m, 4H), 3.31 (s, 2H), 3.60 (s, 2H), 5.35 (s, 2H), 6.17 (s, 1H), 7.14 (s, 2H), 7.39 (d, *J* = 8.1 Hz, 2H), 7.49 (d, *J* = 8.1 Hz, 2H), 8.15 (s, 1H); **$^{13}$C NMR** (126 MHz, DMSO-*d$_6$*) δ 170.3, 168.7, 163.3, 162.8, 137.6, 135.6, 128.1, 127.1, 94.4, 66.9, 54.6, 54.2, 47.0, 45.6, 41.4; **MS** (m/z): [M + H]$^+$ calcd. for $C_{17}H_{21}ClN_5O_2$, 362.138; found 362.139.

**5-(((2-Amino-6-chloropyrimidin-4-yl)oxy)methyl)-*N*-methylpicolinamide (1e)**

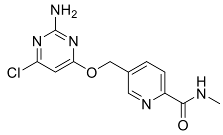

**8c** (75 mg, 45 w%, 0.20 mmol) was used to synthesise **1e** according to general procedure K and stirred overnight. The organics were concentrated and the residue partitioned between ammonium chloride and DCM, the layers separated and the aqueous phase extracted with DCM. The combined organic phase was dried over $MgSO_4$, filtered, concentrated and purified by NPFCC (5 g, 0–100% EtOAc in heptane over 50 CV) to afford **1e** as an off-white film/gum (16 mg, 27%); **$^1$H NMR** (500 MHz, DMSO-*d$_6$*) δ 2.81 (d, *J* = 4.9 Hz, 3H), 5.43 (s, 2H), 6.20 (s, 1H), 7.16 (s, 2H), 8.01–8.06 (m, 2H), 8.72 (s, 1H), 8.75–8.81 (m, 1H); **$^{13}$C NMR** (126 MHz, DMSO-*d$_6$*) δ 170.0, 164.1, 162.7, 160.2, 149.8, 148.3, 137.6, 134.6, 121.5, 94.4, 64.7, 26.0; **MS** (m/z): [M + H]$^+$ calcd. for $C_{12}H_{13}ClN_5O_2$, 294.076; found 294.076.

**6-(((2-amino-6-chloropyrimidin-4-yl)oxy)methyl)-N-methylnicotinamide (1f)**

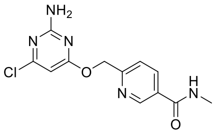

**9a** (53 mg, 0.13 mmol) was used to synthesise **1f** according to general procedure J. The crude product was purified by RPFCC (6 g, 20–45% B in A over 10 CV). The product was lyophilised to give a fluffy pale yellow powder which was triturated with diethyl ether to yield **1f** as a pale yellow solid (22 mg, 56 %); **$^1$H NMR** (500 MHz, DMSO-*d$_6$*) δ 2.80 (3H, d), 3.29 (2H, s), 5.46 (1H, s), 7.11 (2H, s), 7.52 (1H, d), 8.19 (1H, d), 8.63 (1H, s), 8.95 (1H, s). **$^{13}$C NMR** (151 MHz, DMSO-*d$_6$*) δ 26.18, 40.06, 67.49, 94.48, 120.94, 129.09, 135.69, 147.89, 158.42, 160.17, 162.75, 164.88, 170.06. **MS** (m/z): [M - H]$^+$ calcd. for $C_{12}H_{11}ClN_5O_2$, 292.061; found 292.060.

**4-Chloro-6-((4-ethynylbenzyl)oxy)pyrimidin-2-amine (1g)**

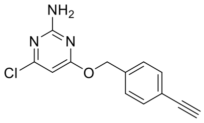

**8a** (67 mg, 0.51 mmol) was used to synthesise **1g** according to general procedure K and stirred overnight. The organics were concentrated and the residue partitioned between ammonium chloride and DCM, the layers separated and the aqueous phase extracted with DCM. The combined organic phase was dried over $MgSO_4$, filtered, concentrated and purified by NPFCC (5 g, 0–27% EtOAc in heptane over 30 CV) and by RPFCC (6 g, 30–80% B in A over 13 CV) to afford **1g** as a white solid/gum (28 mg, 22%); **$^1$H NMR** (500 MHz, DMSO-*d$_6$*) δ 4.21 (s, 1H), 5.33 (s, 2H), 6.17 (s, 1H), 7.13 (s, 2H), 7.41–7.46 (m, 2H), 7.47–7.51 (m, 2H); **$^{13}$C NMR** (126 MHz, DMSO-*d$_6$*) δ 170.2, 162.8, 160.1, 137.2, 131.8, 128.3, 121.4, 94.4, 83.2, 81.2, 66.8; **MS** (m/z): [M + H]$^+$ calcd. for $C_{13}H_{11}ClN_3O$, 260.059; found 260.060.

#### 4-Chloro-6-((3-ethynylbenzyl)oxy)pyrimidin-2-amine (1h)

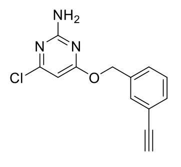

**9b** (31 mg, 0.10 mmol) was used to synthesise **1h** according to general procedure I. The crude product was purified by NPFCC (5 g, 0–15% EtOAc in heptane over 45 CV) to afford **1h** as a white solid/film (8.1 mg, 32 %); **$^1$H NMR** (500 MHz, Chloroform-*d*) δ 3.09 (s, 1H), 5.20 (s, 2H), 5.30 (s, 2H), 6.18 (s, 1H), 7.35 (dt, *J* = 15.1, 7.7 Hz, 2H), 7.46 (d, *J* = 7.4 Hz, 1H), 7.52 (s, 1H); **$^{13}$C NMR** (126 MHz, Chloroform-*d*) δ 170.9, 162.2, 161.0, 136.5, 132.0, 131.7, 128.7, 128.4, 122.6, 97.5, 83.4, 77.7, 67.7; **MS** (m/z): [M + H]$^+$ calcd. for $C_{13}H_{11}ClN_3O$, 260.059; found 260.060.

#### *N*-(4-(((2-Amino-6-chloropyrimidin-4-yl)oxy)methyl)benzyl)acetamide (1i)

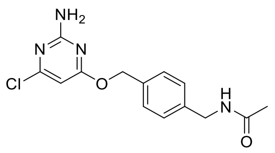

SNAP2 ligand (52.9 mg, 0.20 mmol) and acetic acid (1.1 eq.) was used to synthesise **1i** according to general procedure J using DIPEA (4 eq.) and stirred for 2 h. The crude product was purified by RPFCC (6 g, 5–40% B in A over 10 CV, 40–95% B in A over 4 CV) to afford **1i** as a fluffy white solid (55 mg, 89 %); **$^1$H NMR** (500 MHz, Methanol-*d$_4$*) δ 1.98 (s, 3H), 4.35 (s, 2H), 5.34 (s, 2H), 6.10 (s, 1H), 7.25–7.32 (m, 2H), 7.37–7.42 (m, 2H); **$^{13}$C NMR** (126 MHz, Methanol-*d$_4$*) δ 173.1, 172.4, 164.5, 161.8, 140.0, 136.8, 129.6, 128.7, 96.6, 69.0, 43.9, 22.5; **MS** (m/z): [M + H]$^+$ calcd. for $C_{14}H_{16}ClN_4O_2$, 307.096; found 307.096.

#### *N*-(3-(((2-Amino-6-chloropyrimidin-4-yl)oxy)methyl)benzyl)acetamide (1j)

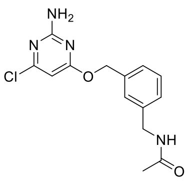

**10a** (65 mg, 0.22 mmol) was used to synthesise **1j** according to general procedure M. The crude product was purified by RPFCC (6g, 20–45% B in A over 10 CV) followed by trituration with diethylether (2 x 3 mL) to afford **1j** as a white sticky solid (31 mg, 47%); **$^1$H NMR** (500 MHz, DMSO-*d$_6$*) δ 1.86 (s, 3H), 4.25 (d, *J* = 6.0 Hz, 2H), 5.30 (s, 2H), 6.15 (s, 1H), 7.12 (s, 2H), 7.21 (d, *J* = 7.2 Hz, 1H), 7.28–7.36 (m, 3H), 8.36 (t, *J* = 5.5 Hz, 1H); **$^{13}$C NMR** (126 MHz, DMSO-*d$_6$*) δ 170.3, 169.2, 162.8, 160.1, 139.9, 136.2, 128.5, 127.2, 127.1, 126.8, 94.4, 67.4, 42.0, 22.6, **MS** (m/z): [M + H]$^+$ calcd. for $C_{14}H_{16}ClN_4O_2$, 307.096; found 307.096.

#### *N*-(2-(((2-Amino-6-chloropyrimidin-4-yl)oxy)methyl)benzyl)acetamide (1k)

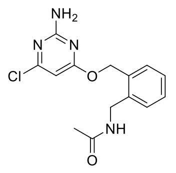

**10b** (65 mg, 0.22 mmol) was used to synthesise **1k** according to general procedure M. The crude product was purified by RPFCC (6 g, 20–40% B in 0.1% ammonia in water over 12 CV) followed by NPFCC (5 g, 50–100 % EtOAc in heptane over 10 CV, 100 % EtOAc over 5 CV) to afford **1k** as a colourless film (3.4 mg, 5 %); **$^1$H NMR** (500 MHz, Acetone-*d$_6$*) δ 1.95 (s, 3H), 4.59 (d, *J* = 5.9 Hz, 2H), 5.40 (s, 2H), 6.09 (s, 1H), 6.50 (s, 2H), 7.26 (td, *J* = 7.4, 1.5 Hz, 1H), 7.30 (td, *J* = 7.5, 1.5 Hz, 1H), 7.34–7.37 (m, 1H), 7.44 (dd, *J* = 7.4, 1.2 Hz, 1H), 7.51 (s, 1H); **$^{13}$C NMR** (126 MHz, Acetone-*d$_6$*) δ 171.5, 170.0, 164.1, 161.5, 139.0, 135.2, 130.4, 129.2, 128.7, 127.9, 96.0, 66.1, 40.8, 22.9; **MS** (m/z): [M + H]$^+$ calcd. for $C_{14}H_{16}ClN_4O_2$, 307.096;

#### 4-Chloro-6-((4-methoxybenzyl)oxy)pyrimidin-2-amine (1l)

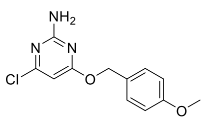

(4-Methoxyphenyl)methanol (50 mg, 0.36 mmol) was used to synthesise **1l** according to general procedure K and stirred overnight. The organics were concentrated and the residue partitioned between ammonium chloride and DCM, the layers separated and the aqueous phase extracted with DCM. The combined organic phase was dried over $MgSO_4$, filtered, concentrated and purified by NPFCC (5 g, 15–30% EtOAc in heptane over 15 CV) to afford **1l** as a white solid (79 mg, 82%); **$^1$H NMR** (500 MHz, Chloroform-*d*) δ 3.75 (s, 3H), 5.23 (s, 2H), 6.11 (s, 1H), 6.9–6.96 (m, 2H), 7.11 (s, 2H), 7.35–7.42 (m, 2H); **$^{13}$C NMR** (126 MHz, Chloroform-*d*) δ 170.4, 162.8, 160.0,

159.2, 130.4, 130.3, 128.1, 113.8, 113.8, 113.8, 94.4, 67.3, 55.1; **MS** (m/z): $[M + H]^+$ calcd. for $C_{12}H_{13}ClN_3O_2$, 266.070; found 266.071.

**4-Chloro-6-((3-methoxybenzyl)oxy)pyrimidin-2-amine (1m)**

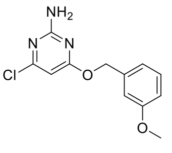

(3-Methoxyphenyl)methanol (28 mg, 0.20 mmol) was used to synthesise **1m** according to general procedure K and stirred for 2 d. The crude product was purified by RPFCC (6 g, 30–65% B in A over 10 CV) to afford **1m** as a white solid (33 mg, 62%); $^1$**H NMR** (500 MHz, DMSO-*d*$_6$) δ 3.75 (s, 3H), 5.28 (s, 2H), 6.16 (s, 1H), 6.87–6.93 (m, 1H), 6.97–7.02 (m, 2H), 7.12 (s, 2H), 7.29 (t, *J* = 8.0 Hz, 1H); $^{13}$**C NMR** (126 MHz, DMSO-*d*$_6$) δ 170.3, 162.8, 160.1, 159.3, 137.8, 129.6, 120.3, 113.8, 113.6, 94.4, 67.3, 55.1; **MS** (m/z): $[M + H]^+$ calcd. for $C_{12}H_{13}ClN_3O_2$, 266.070; found 266.071.

**Negative control compounds for VHL and CRBN PROTACs**

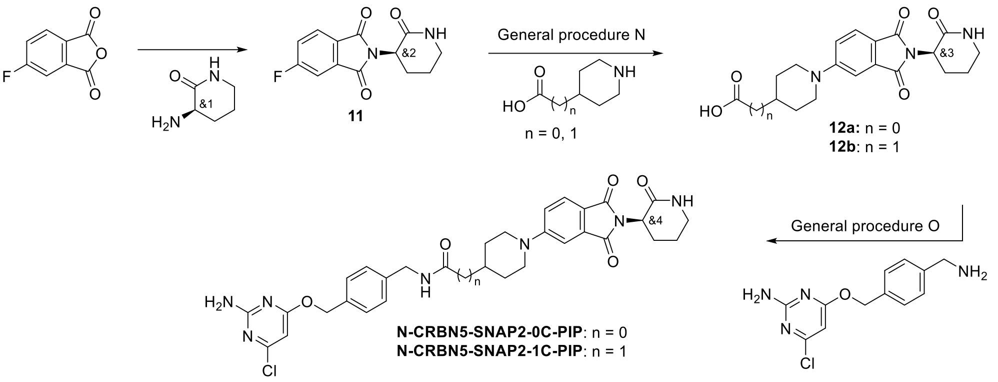


**Scheme S5**. Overview synthesis of CRBN negative control compounds.

**General procedure N – SNAR**

The corresponding aryl fluoride (1.0 eq.) and the corresponding amine (1.0-1.2 eq.) were dissolved in DMSO (1 mL) and DIPEA (2.2 eq.) was added and the solution was stirred at 100 °C overnight. The reaction mixture was concentrated and purified by RPFCC to afford products **12a-b**.

**General procedure O – Amide coupling**

The corresponding acid (1.0 eq.), SNAP2 ligand (1.0-1.5 eq.) and HATU (1.0 eq.) were dissolved in DCM (1 mL) and DIPEA (4.0 eq.) was added and the solution stirred at room temperature until completion monitored by LCMS. The reaction mixture was concentrated and purified by RPFCC for afford the desired PROTACs.

***Rac*-5-fluoro-2-(2-oxopiperidin-3-yl)isoindoline-1,3-dione (11)**

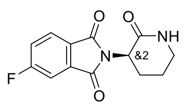

5-Fluoroisobenzofuran-1,3-dione (700 mg, 4.21 mmol), *rac*-(*R*)-3-aminopiperidin-2-one (505 mg, 4.42 mmol) and sodium acetate (1037 mg, 12.64 mmol) were dissolved in acetic acid (5 mL) and stirred at 80 °C for 20 min. The reaction was neutralised by addition of $K_2CO_3$ solution (50 mL) followed by addition of EtOAc and a rough separation of the phases. The aqueous phase was extracted with EtOAc (x4). The combined organic phase was dried over $MgSO_4$ filtered and solvent was removed under reduced pressure. the same round bottom flask before loading concentrated and loaded on celite. The crude product was purified by RPFCC (30 g, 5-50% B in A over 12 CV) to afford **11** as a white solid (364 mg, 28%). $^1$**H NMR** (500 MHz, DMSO-*d*$_6$) δ

1.00–1.12 (m, 2H), 1.15–1.22 (m, 1H), 1.38 (qd, *J* = 12.1, 4.8 Hz, 1H), 2.33–2.44 (m, 2H), 3.78 (dd, *J* = 12.0, 6.3 Hz, 1H), 6.86 (ddd, *J* = 9.4, 8.2, 2.3 Hz, 1H), 6.97 (dd, *J* = 7.5, 2.3 Hz, 1H), 7.05 (d, *J* = 3.1 Hz, 1H), 7.14 (dd, *J* = 8.2, 4.5 Hz, 1H). **MS** (m/z): $[M + H]^+$ calcd. for $C_{13}H_{11}FN_2O_3$, 263.1; found 263.0.

#### ***Rac*-1-(1,3-dioxo-2-(2-oxopiperidin-3-yl)isoindolin-5-yl)piperidine-4-carboxylic acid (12a)**

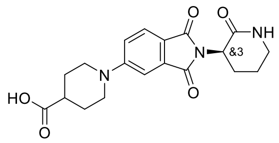


**11** (150 mg, 0.57 mmol) and piperidine-4-carboxylic acid (1.2 eq.) were used to synthesise **12a** according to general procedure N. The crude product was purified by RPFCC (12 g, 5-11% B in 0.1% ammonia in water over 9 CV) to afford **12a** as a yellow/orange solid (185 mg, 83%). $^1$**H NMR** (500 MHz, DMSO-$d_6$) δ 1.53–1.63 (m, 2H), 1.86 (dt, *J* = 17.8, 4.4 Hz, 4H), 1.95 (ddd, *J* = 10.7, 5.8, 2.2 Hz, 1H), 2.20 (qd, *J* = 12.1, 3.8 Hz, 1H), 2.4–2.47 (m, 1H), 3.05 (ddd, *J* = 13.5, 11.3, 2.9 Hz, 2H), 3.17–3.26 (m, 2H), 3.92 (dt, *J* = 13.1, 4.0 Hz, 2H), 4.52 (dd, *J* = 11.9, 6.2 Hz, 1H), 7.21 (dd, *J* = 8.6, 2.4 Hz, 1H), 7.28 (d, *J* = 2.3 Hz, 1H), 7.61 (d, *J* = 8.5 Hz, 1H), 7.80 (d, *J* = 3.2 Hz, 1H). **MS** (m/z): $[M + H]^+$ calcd. for $C_{19}H_{21}N_3O_5$, 372.2; found 372.2.

#### ***Rac*-2-(1-(1,3-dioxo-2-(2-oxopiperidin-3-yl)isoindolin-5-yl)piperidin-4-yl)acetic acid (12b)**

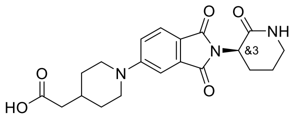


**11** (150 mg, 0.57 mmol) and 2-(piperidin-4-yl)acetic acid (1.0 eq.) were used to synthesise **12b** according to general procedure N. The crude product was purified by RPFCC (12 g, 5-15% B in 0.1% ammonia in water over 8 CV) to afford **12b** as a yellow/orange solid (190 mg, 83%). $^1$**H NMR** (500 MHz, DMSO-$d_6$) δ 1.13–1.24 (m, 2H), 1.7–1.78 (m, 2H), 1.8–1.98 (m, 4H), 2.09 (d, *J* = 6.9 Hz, 2H), 2.20 (qd, *J* = 12.1, 3.7 Hz, 1H), 2.93 (td, *J* = 12.8, 2.6 Hz, 2H), 3.22 (qd, *J* = 12.2, 6.3 Hz, 2H), 4.00 (dt, *J* = 13.3, 3.3 Hz, 2H), 4.51 (dd, *J* = 11.9, 6.2 Hz, 1H), 7.20 (dd, *J* = 8.6, 2.4 Hz, 1H), 7.26 (d, *J* = 2.3 Hz, 1H), 7.60 (d, *J* = 8.5 Hz, 1H), 7.80 (d, *J* = 3.1 Hz, 1H). **MS** (m/z): $[M + H]^+$ calcd. for $C_{20}H_{23}N_3O_5$, 386.2; found 386.4.

#### ***Rac-N*-(4-(((2-amino-6-chloropyrimidin-4-yl)oxy)methyl)benzyl)-1-(1,3-dioxo-2-(2-oxopiperidin-3-yl)isoindolin-5-yl)piperidine-4-carboxamide (N-CRBN5-SNAP2-0C-PIP)**

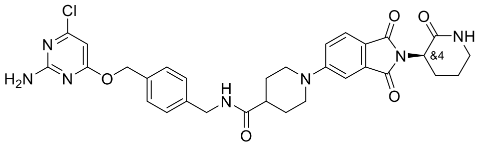


**12a** (50 mg, 0.12 mmol) and SNAP2 ligand (1.2 eq.) were used to synthesise **N-CRBN5-SNAP2-0C-PIP** according to general procedure O. The reaction was stirred for 1 h. The crude product was purified by RPFCC (6 g, 30-43% B in A over 15 CV) to afford **N-CRBN5-SNAP2-0C-PIP** as a yellow solid (52 mg, 65%). $^1$**H NMR** (500 MHz, DMSO-$d_6$) δ 1.58–1.68 (m, 2H), 1.76–1.82 (m, 2H), 1.83–1.99 (m, 3H), 2.20 (qd, *J* = 12.1, 3.9 Hz, 1H), 2.51–2.52 (m, 1H), 2.99 (td, *J* = 12.9, 2.7 Hz, 2H), 3.15–3.25 (m, 2H), 4.06 (d, *J* = 13.1 Hz, 2H), 4.26 (d, *J* = 5.9 Hz, 2H), 4.52 (dd, *J* = 11.9, 6.2 Hz, 1H), 5.28 (s, 2H), 6.13 (s, 1H), 7.11 (s, 2H), 7.23 (dd, *J* = 9.4, 3.1 Hz, 3H), 7.29 (d, *J* = 2.3 Hz, 1H), 7.38 (d, *J* = 8.0 Hz, 2H), 7.62 (d, *J* = 8.5 Hz, 1H), 7.80 (d, *J* = 3.1 Hz, 1H), 8.38 (t, *J* = 6.0 Hz, 1H). **MS** (m/z): $[M + H]^+$ calcd. for $C_{31}H_{33}ClN_7O_5$, 618.223; found 618.221.

**_Rac-N_-(4-(((2-amino-6-chloropyrimidin-4-yl)oxy)methyl)benzyl)-2-(1-(1,3-dioxo-2-(2-oxopiperidin-3-yl)isoindolin-5-yl)piperidin-4-yl)acetamide (N-CRBN5-SNAP2-1C-PIP)**

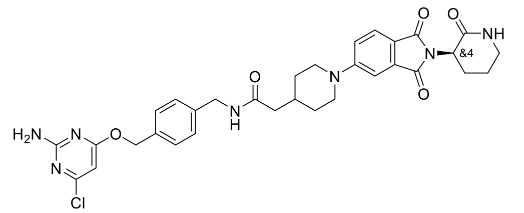

**12b** (50 mg, 0.12 mmol) and SNAP2 ligand (1.0 eq.) were used to synthesise **N-CRBN5-SNAP2-1C-PIP** according to general procedure O. The reaction was stirred for 1 h. The crude product was purified by RPFCC (12 g, 35-45% B in A over 12 CV) to afford **N-CRBN5-SNAP2-1C-PIP** as a yellow solid (35 mg, 45%). **$^1$H NMR** (500 MHz, DMSO-*d$_6$*) δ 1.16–1.27 (m, 2H), 1.71 (d, *J* = 12.6 Hz, 2H), 1.83–2.03 (m, 4H), 2.09–2.11 (m, 2H), 2.20 (qd, *J* = 12.1, 3.9 Hz, 1H), 2.95 (td, *J* = 13.0, 2.6 Hz, 2H), 3.14–3.26 (m, 2H), 4.01 (d, *J* = 12.9 Hz, 2H), 4.26 (d, *J* = 5.9 Hz, 2H), 4.52 (dd, *J* = 11.9, 6.2 Hz, 1H), 5.29 (s, 2H), 6.13 (s, 1H), 7.11 (s, 2H), 7.21 (dd, *J* = 8.7, 2.4 Hz, 1H), 7.23–7.29 (m, 3H), 7.37–7.41 (m, 2H), 7.61 (d, *J* = 8.6 Hz, 1H), 7.80 (d, *J* = 3.2 Hz, 1H), 8.35 (t, *J* = 5.9 Hz, 1H). **MS** (m/z): [M + H]$^+$ calcd. for $C_{32}H_{35}ClN_7O_5$, 632.239; found 632.235.

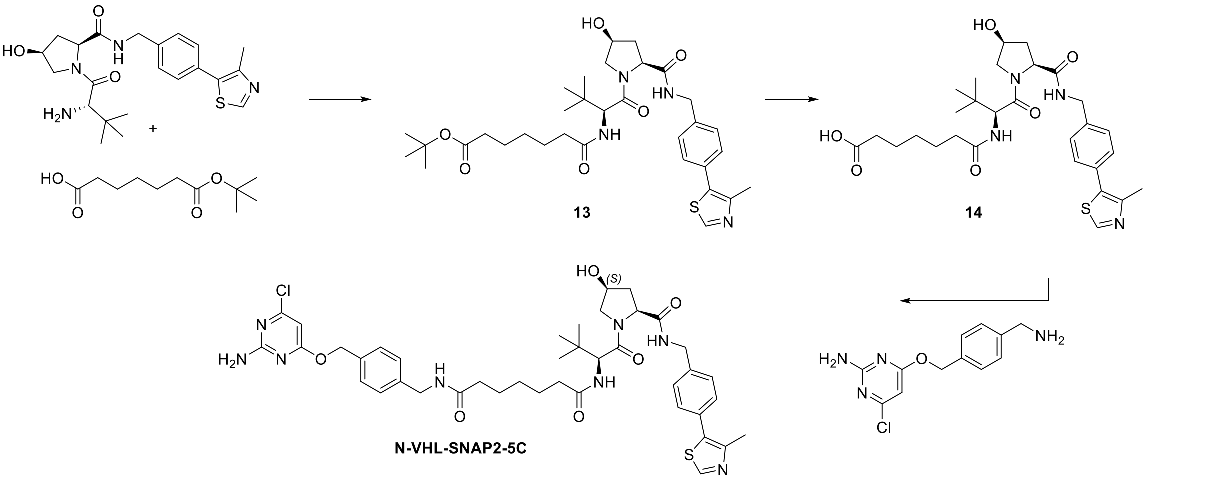


**Scheme S6**. Overview synthesis of VHL negative control compounds.

**_Tert_-butyl 7-(((*S*)-1-((2*S*,4*S*)-4-hydroxy-2-((4-(4-methylthiazol-5-yl)benzyl)carbamoyl)pyrrolidin-1-yl)-3,3-dimethyl-1-oxobutan-2-yl)amino)-7-oxoheptanoate (13)**

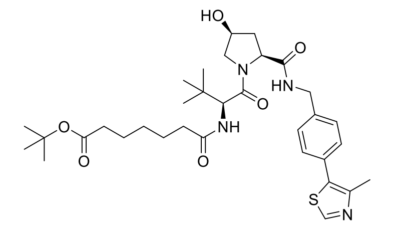

(2*S*,4*S*)-1-((*S*)-2-Amino-3,3-dimethylbutanoyl)-4-hydroxy-*N*-(4-(4-methylthiazol-5-yl)benzyl)pyrrolidine-2-carboxamide hydrochloride (135 mg, 0.29 mmol) and 7-(*tert*-butoxy)-7-oxoheptanoic acid were dissolved in DCM (1 mL) and T3P (2 eq.) and DIPEA (6 eq.) added. The solution was stirred at room temperature for 1 h. The reaction mixture was concentrated and purified by RPFCC (12 g, 40-50% B in A over 10 CV) to afford **13** as a colourless gum (106 mg, 58 %). **$^1$H NMR** (500 MHz, Chloroform-*d*) δ 0.91 (s, 9H), 1.28–1.37 (m, 2H), 1.43 (s, 9H), 1.53–1.68 (m, 5H), 2.17–2.23 (m, 5H), 2.37 (d, *J* = 14.2 Hz, 1H), 2.56 (s, 3H), 3.81 (dd, *J* = 11.0, 1.5 Hz, 1H), 3.93 (dd, *J* = 11.0, 4.2 Hz, 1H), 4.31 (dd, *J* = 15.0, 5.0 Hz, 1H), 4.48 (s, 1H), 4.51 (d, *J* = 9.1 Hz, 1H), 4.66 (dd, *J* = 14.9, 7.1 Hz, 1H), 4.72 (d, *J* = 9.0 Hz, 1H), 5.94 (d, *J* = 9.1 Hz, 1H), 7.34–7.41 (m, 4H), 7.45 (t, *J* = 6.1 Hz, 1H), 8.79 (s, 1H). **MS** (m/z): [M + H]$^+$ calcd. for $C_{33}H_{48}N_4O_6S$, 629.3; found 629.5.

#### 7-(((*S*)-1-((2*S*,4*S*)-4-Hydroxy-2-((4-(4-methylthiazol-5-yl)benzyl)carbamoyl)pyrrolidin-1-yl)-3,3-dimethyl-1-oxobutan-2-yl)amino)-7-oxoheptanoic acid (14)

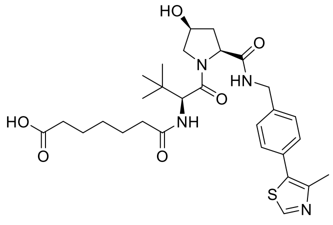

**13** (105 mg, 0.17 mmol) was dissolved in DCM (1 mL) and hydrochloric acid (dioxane, 4 M, 30 eq.) was added. The reaction mixture was stirred at room temperature for 20 min. After concentrating at low temperature methyl ester was formed as indicated by LCMS and NMR analysis. To the resulting gum was added LiOH (10 eq.) in water (1 mL). The reaction mixture was stirred at room temperature for 3 h. The reaction mixture was concentrated at low temperature and redissolved in acetone and filtered to afford **14** as a colourless gum/oil (79 mg, 82 %). **$^1$H NMR** (500 MHz, DMSO-*d*$_6$) δ 0.94 (s, 9H), 1.2–1.26 (m, 2H), 1.46 (h, *J* = 7.0 Hz, 4H), 1.72 (dt, *J* = 12.4, 6.2 Hz, 1H), 2.07–2.25 (m, 4H), 2.28–2.36 (m, 1H), 2.44 (s, 3H), 3.63–3.74 (m, 1H), 3.93 (dd, *J* = 10.1, 5.7 Hz, 1H), 4.17–4.29 (m, 2H), 4.36 (dd, *J* = 8.5, 6.2 Hz, 1H), 4.39–4.47 (m, 2H), 5.48 (s, 1H), 7.39 (q, *J* = 8.3 Hz, 4H), 7.87 (d, *J* = 8.8 Hz, 1H), 8.69 (t, *J* = 6.1 Hz, 1H), 8.99 (s, 1H). **MS** (m/z): $[M + H]^+$ calcd. for $C_{29}H_{40}N_4O_6S$, 573.3; found 573.5.

#### *N*$^1$-(4-(((2-Amino-6-chloropyrimidin-4-yl)oxy)methyl)benzyl)-*N*$^7$-((*S*)-1-((2*S*,4*S*)-4-hydroxy-2-((4-(4-methylthiazol-5-yl)benzyl)carbamoyl)pyrrolidin-1-yl)-3,3-dimethyl-1-oxobutan-2-yl)heptanediamide (N-VHL-SNAP2-5C)

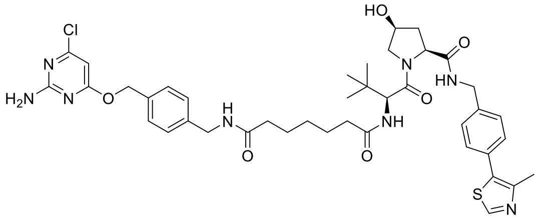

**14** (40 mg, 0.07 mmol), SNAP2 ligand (1.2 eq.) and HATU (1.0 eq.) were dissolved in a mixture of DCM:DMF (10:3, 1.3 mL) and DIPEA (3.0 eq.) was added. The reaction mixture was stirred at room temperature overnight. The reaction mixture was concentrated and purified by RPFCC (6 g, 30-47% B in A over 15 CV) to afford **N-VHL-SNAP2-5C** as a colourless film (13.0 mg, 23 %). **$^1$H NMR** (500 MHz, Chloroform-*d*) δ 0.92 (s, 9H), 1.32 (tt, *J* = 8.0, 6.1 Hz, 2H), 1.58–1.67 (m, 4H), 2.12–2.23 (m, 5H), 2.30 (d, *J* = 14.1 Hz, 1H), 2.50 (s, 3H), 3.78–3.84 (m, 1H), 3.89 (dd, *J* = 11.0, 4.1 Hz, 1H), 4.29 (dd, *J* = 15.0, 5.0 Hz, 1H), 4.34–4.47 (m, 3H), 4.50 (d, *J* = 9.0 Hz, 1H), 4.64 (dd, *J* = 14.9, 7.0 Hz, 1H), 4.70 (d, *J* = 9.0 Hz, 1H), 5.23–5.38 (m, 4H), 5.64 (s, 1H), 5.89–5.95 (m, 1H), 6.12 (s, 1H), 6.16 (d, *J* = 8.9 Hz, 1H), 7.22–7.26 (m, 2H), 7.32–7.37 (m, 6H), 7.69 (dd, *J* = 7.0, 5.0 Hz, 1H), 8.68 (s, 1H). **MS** (m/z): $[M + H]^+$ calcd. for $C_{41}H_{52}ClN_8O_6S$, 819.342; found 819.341.
